# CDSeqR: fast complete deconvolution for gene expression data from bulk tissues

**DOI:** 10.1101/2021.01.30.428954

**Authors:** Kai Kang, Caizhi David Huang, Yuanyuan Li, David M. Umbach, Leping Li

## Abstract

**Background:** Biological tissues consist of heterogenous populations of cells. Because gene expression patterns from bulk tissue samples reflect the contributions from all cells in the tissue, understanding the contribution of individual cell types to the overall gene expression in the tissue is fundamentally important. We recently developed a computational method, CDSeq, that can simultaneously estimate both sample-specific cell-type proportions and cell-type-specific gene expression profiles using only bulk RNA-Seq counts from multiple samples. Here we present an R implementation of CDSeq (CDSeqR) with significant performance improvement over the original implementation in MATLAB and an added new function to aid cell type annotation. The R package would be of interest for the broader R community.

**Result:** We developed a novel strategy to substantially improve computational efficiency in both speed and memory usage. In addition, we designed and implemented a new function for annotating the CDSeq estimated cell types using single-cell RNA sequencing (scRNA-seq) data. This function allows users to readily interpret and visualize the CDSeq estimated cell types. In addition, this new function further allows the users to annotate CDSeq-estimated cell types using marker genes. We carried out additional validations of the CDSeqR software using synthetic, real cell mixtures, and real bulk RNA-seq data from the Cancer Genome Atlas (TCGA) and The Genotype-Tissue Expression (GTEx) project.

**Conclusions:** The existing bulk RNA-seq repositories, such as TCGA and GTEx, provide enormous resources for better understanding changes in transcriptomics and human diseases. They are also potentially useful for studying cell-cell interactions in the tissue microenvironment. Bulk level analyses neglect tissue heterogeneity, however, and hinder investigation of a cell-type-specific expression. The CDSeqR package may aid *in silico* dissection of bulk expression data, enabling researchers to recover cell-type-specific information.

## Background

Many biological samples present heterogeneity; namely, their cellular composition varies phenotypically and functionally. The cell-type proportions can exhibit high variability cross samples from the same tissue type, potentially indicating the occurrence of various physiological and pathological activities in the samples. Therefore, knowing the cellular composition of heterogeneous samples is important to better understand the roles of distinct cell populations. For example, dissection of tumor-infiltrating immune cells holds promise for gaining deeper insights into the underlying mechanisms of cancers, for discovering predictive biomarkers, as well as for the development of novel treatment strategies [1,2]. Although single-cell RNA sequencing has become popular, in addition to its high cost and sparse read counts, isolating cells from solid tissues especially frozen tissues remains challenging. Moreover, the inevitable sparsity of RNA molecules sequenced from frozen samples further challenges computational techniques. Therefore, bulk RNA-seq data are still routinely generated in many research projects. The heterogeneous nature of bulk RNA-seq, however, hampers efforts to study how individual cell types in the tissue microenvironment influence the underlying biology of interest. Computational deconvolution methods [3,4,13–16,5–12] are complementary to single-cell analysis and are very useful tools for extracting cell-type-specific signals from bulk measurements.

This work describes an R package (CDSeqR) implementation of CDSeq (complete deconvolution using sequencing data) [5] that we recently developed in MATLAB. CDSeq was designed to simultaneously estimate the cell-type-specific gene expression profiles (GEPs) and the sample-specific cell-type proportions using bulk RNA-seq data without the need of reference GEPs (i.e. expression of pure cell lines or scRNA-seq data). Here, we not only implemented a user-friendly R version for CDSeq but also substantially improved its computational efficiency in both speed and memory usage. Specially, we designed a dimension Reduction-Recover approach that is automatically parallelized to facilitate computation. In addition, we implemented a new functionality for annotating the CDSeq-estimated cell types using scRNA-seq data. Moreover, we employed a recently published deconvolution benchmarking pipeline [17] to thoroughly test CDSeqR using both cell mixtures [18] and simulated data [19–21]. Throughout the manuscript, we refer to “CDSeqR” as the package and “CDSeq” as the deconvolution procedure.

## Implementation

### Statistical model

CDSeq employs a hierarchical Bayesian modeling strategy to account for the mixing process of observed RNA reads from different cell types. Specifically, the gene expression of any particular cell type, in the form of read counts, is modeled by a Dirichlet multinomial random variable and the bulk measure is therefore a weighted sum of all cell types (i.e., Dirichlet multinomial random variables) constituting the bulk tissue. Then, CDSeq uses a Gibbs sampler [5,22] for parameter estimation. A detailed description of the model and solver can be found in [5].

### A dimension reduce-recover strategy for fast deconvolution

A practical challenge for our original MATLAB implementation of CDSeq was the high computational demand, especially for large datasets. The Gibbs sampler employed for parameter estimation requires storage of several large vectors whose sizes are data-dependent; therefore, it can be computationally demanding especially for large datasets. We propose a novel strategy for alleviating this issue. Our strategy, named dimension Reduce-Recover, consists of two steps: dimension reduction and dimension recovery. This approach was based on the following assumption: cell proportions in each sample are invariant with respect to the genes used in deconvolution. Specifically, for a bulk RNA-seq sample, the proportions of cell types can be estimated using multiple subsets of randomly selected genes. Therefore, for a bulk RNA-Seq dataset, we should be able to obtain the estimated cell-type proportions for each sample in the dataset regardless of the subset of genes considered. This idea allows us to calculate multiple independent estimates of cell-type proportions using many subsets of randomly selected genes processed in parallel. The final estimate of the cell-type proportions is the average of the independent estimates. We then estimate the cell-type specific gene expression profiles by post-multiplying the input data matrix by a generalized inverse of the matrix of final estimated cell-type proportions (Supplementary Methods). We have carried out thorough testing and demonstrated that this approach substantially speeds up computation without sacrificing the performance (see Supplementary Note for details, and Supplementary Fig. S9). We further evaluated the effects of the size of the gene list and the block number on the performance when using the Reduce-Recovery strategy (Supplementary Fig. S9).

### Cell type assignment function in CDSeqR

Bulk tissue RNA-seq samples usually present a great degree of heterogeneity. For example, cancer cells in a tumor exhibit a spectrum of phenotypic and transcriptomic stages [23]. This spectrum makes the assignment of identified cell types to known cell types challenging. In our original MATLAB version, the cell-type assignment was based on correlation analysis using a user-provided pure cell line GEPs. While this option remains available, the increasing number of publicly available single cell data sets, e.g. [21,24]., provide alternative GEPs for cell-type assignment. Thus, in CDSeqR, we designed and implemented a new function to perform cell-type assignment for the downstream processing of CDSeq estimated cell-type-specific GEPs. In addition, there may be cases where reliable reference datasets from either pure cell line or single cell RNA-seq may not be available. Annotation of CDSeq estimated cell types using cell type marker genes can also be performed in CDSeqR.

Specifically, this new function, *cellTypeAssignSCRNA*, embodies three approaches to annotate the CDSeq-estimated cell types. First, it employs correlation analysis to measure the similarity between the CDSeq-estimated cell-type-specific GEPs and the summarized cell-type specific gene expression profiles from scRNA-seq data. When scRNA-seq reference data are provided, *cellTypeAssignSCRNA* first sums up the read counts of all cells from the same cell type to form the aggregated cell-type specific gene expression profiles for each cell type in the scRNA-seq data. Next, *cellTypeAssignSCRNA* performs pair-wise correlation analysis between the CDSeq estimated GEPs and the scRNA-seq GEPs and assign the CDSeq estimated cell type to the cell type in the scRNA-seq data with which it has the highest correlation. A threshold for the correlation can be set *a priori.* Any CDSeq estimated cell types that failed to achieve the correlation threshold would be considered unannotated. Note that it is possible that more than one CDSeq estimated cell types can match the same cell type.

Second, cell annotation can also be achieved by clustering both the user-provided scRNA-seq reference data with the pseudo scRNA-seq data generated from the CDSeq estimated cell-type specific gene expression profiles. Under the assumption that the total read count of each CDSeq estimated cell type follows a negative binomial distribution, *cellTypeAssignSCRNA* generates the total synthetic read count for each CDSeq cell type using the CDSeq estimated GEPs for the cell using the R function *rnbinom* (*n*, *size*, *mu*) where *n* can be any integer greater than or equal to 1 (See Supplementary Figures S11-S16) and, by default, *mu* (mean) and *size* (dispersion) are estimated from the reference scRNA-seq. These default parameters in *rnbinom* can be replaced by user-provided parameters. Next, *cellTypeAssignSCRNA* generates the synthetic single-cell read counts for each CDSeq cell type using the CDSeq-estimated GEPs and the R function *rmultinom*(*n*, *size*, *prob*), with *n* being the number of pseudo cells per CDSeq estimated cell type (same as the n in *rnbinom*), *size* set equal to the total read count for the cell type as generated by the preceding call to *rnbinom*, and *prob* provided by CDSeq-estimated cell-type-specific gene expression profile expressed as a multinomial parameter vector (i.e., expression levels expressed as proportions that sum to one). The resulting synthetic scRNA-seq data can then subsequently be combined with user provided scRNA-seq data for clustering analysis and cell type annotation using the Seurat package [25,26]. The clustering algorithm implemented in Seurat is a graph-based approach based on the outcomes of principal components analysis. The number of clusters cannot be set explicitly but is affected by a resolution parameter; the larger the resolution parameter is the more clusters will be returned. The cell types identified by CDSeq from bulk RNA-seq data were clustered with the individual cells from the scRNA-seq data. We then assigned CDSeq-identified cell types to the cell type from the scRNA-seq data to which the majority of the cells in the cluster belong (Fig 1).

**Figure 1.**
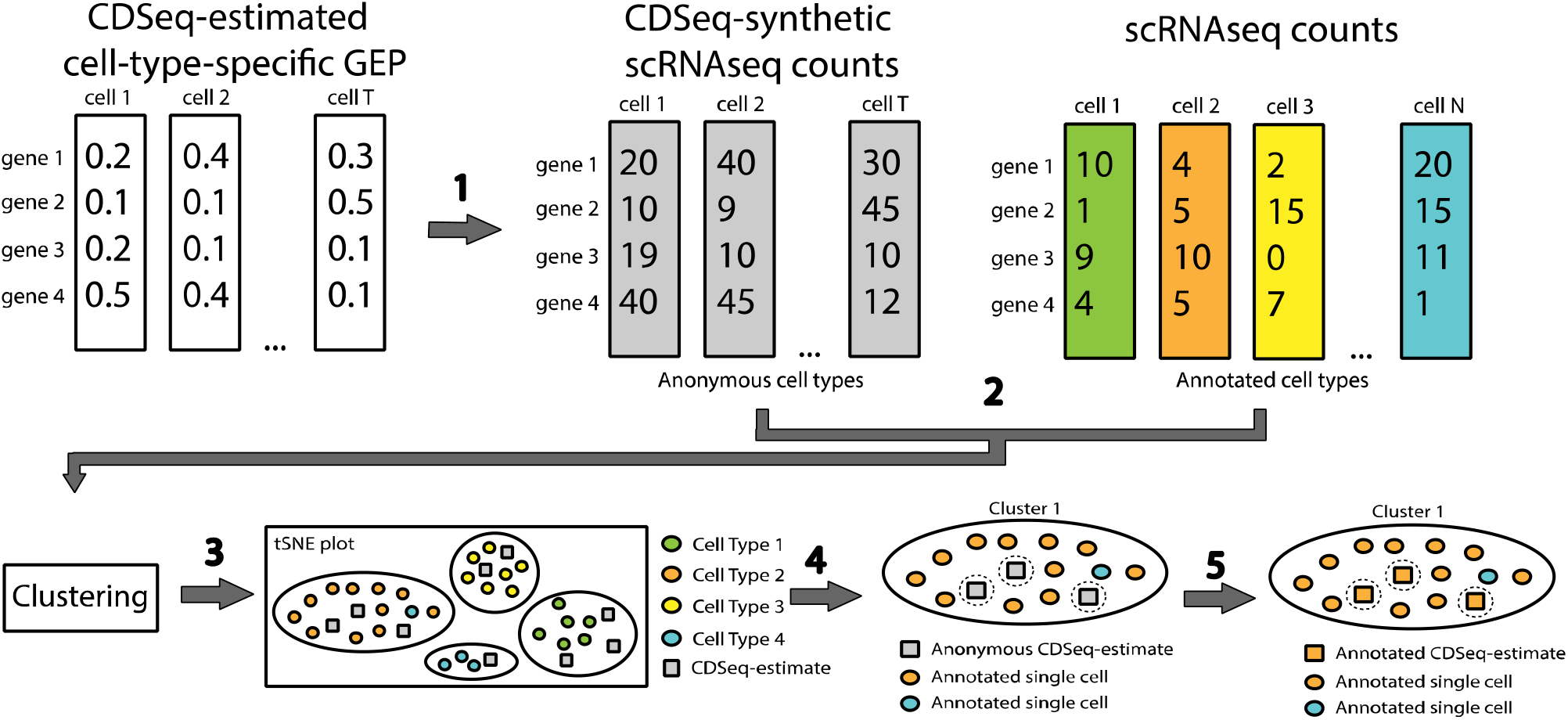
Schematic illustration of our cell-type assignment approach. It consists of five steps: 1) generating synthetic single-cell read counts (gray rectangles) for the CDSeq-estimated cell-type-specific GEPs (white rectangles); 2) combining the pseudo scRNA-seq data with user-provided scRNA-seq data (colored rectangles), then performing clustering using Seurat; 3) visualizing the result using tSNE plot to show the 2D embedding of the clustering outcomes with differently colored circles denoting distinctive cell types in the single-cell data and gray squares denoting the CDSeq-identified cell types; 4) annotating each cluster to the cell type to which the majority of the cells in the cluster belong; 5) assigning the CDSeq-identified cell types accordingly as in step 4.

Lastly, in the case where the reference data are not available, *cellTypeAssignSCRNA* carried out clustering analysis of the pseudo single-cell data generated from the CDSeq-estimated GEPs using Seurat [25,26] and identifies the marker genes in each cluster for cell type annotation.

The marker genes for each cluster are identified by comparing their expression levels in the cluster with those in all other clusters. We show that this approach works reasonably well (Fig. 4 and Supplementary Fig. S17, S18, S19 and S21) using synthetic mixtures of human brain cells [21].

**Figure 2.**
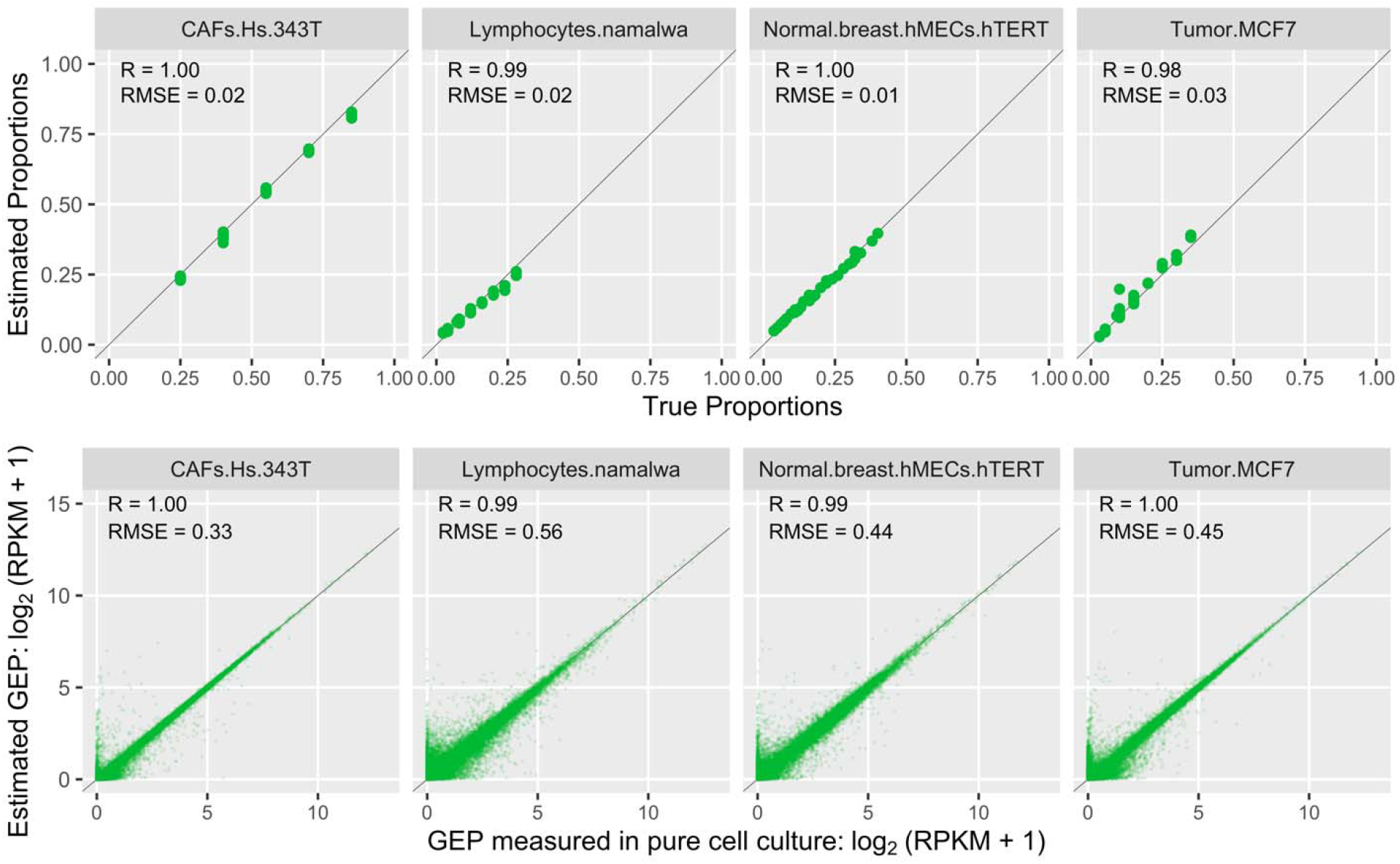
Results of CDSeqR analysis of the *in vitro* cell mixtures. The mixtures comprise 4 four cell types: cancer-associated fibroblast, lymphocytes, normal breast cells, tumor cells (see Supplementary Methods for details). We ran CDSeqR on 32 bulk RNA-seq samples with known cell proportions in each sample [18] using 7 randomly sampled gene subsets of size 500 (7 data blocks, each of 500 genes). In the first row, each point represents a sample (n=32); in the second row, each point represents a gene (n=~19,000). True and estimated GEPs are normalized as RPKM. The line in each panel is the x=y line.

**Figure 3.**
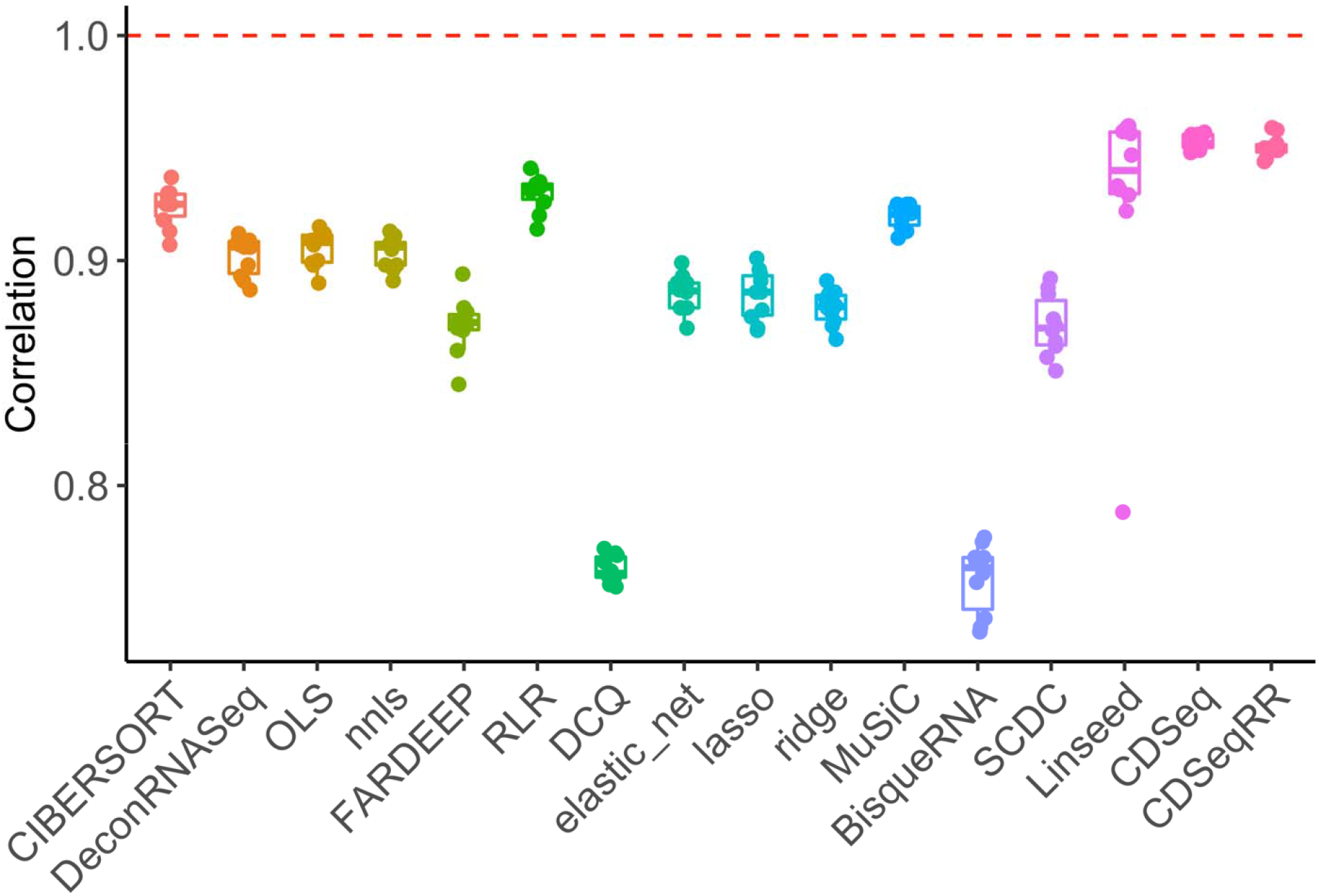
Comparisons with existing deconvolution methods using data generated from benchmark pipeline [17]. We compared CDSeqR with 14 other deconvolution methods using synthetic mixtures generated from three scRNA-seq datasets [20,21,41]. The results for the PBMC dataset [41] are shown here. The X-axis represents different deconvolution methods whereas the Y-axis denotes the correlation between the estimated cell type proportions by each method and the true cell type proportions. We generated 10 sets of 100 mixtures. Each dot in the box plots represents a deconvolution result from one of the 10 sets. CDSeqRR refers to deconvolution with the Reduce-Recover strategy whereas CDSeq denotes deconvolution without using Reduce-Recover option Additional results of the comparisons are available in Supplementary Fig. S6-S8.

**Figure 4.**
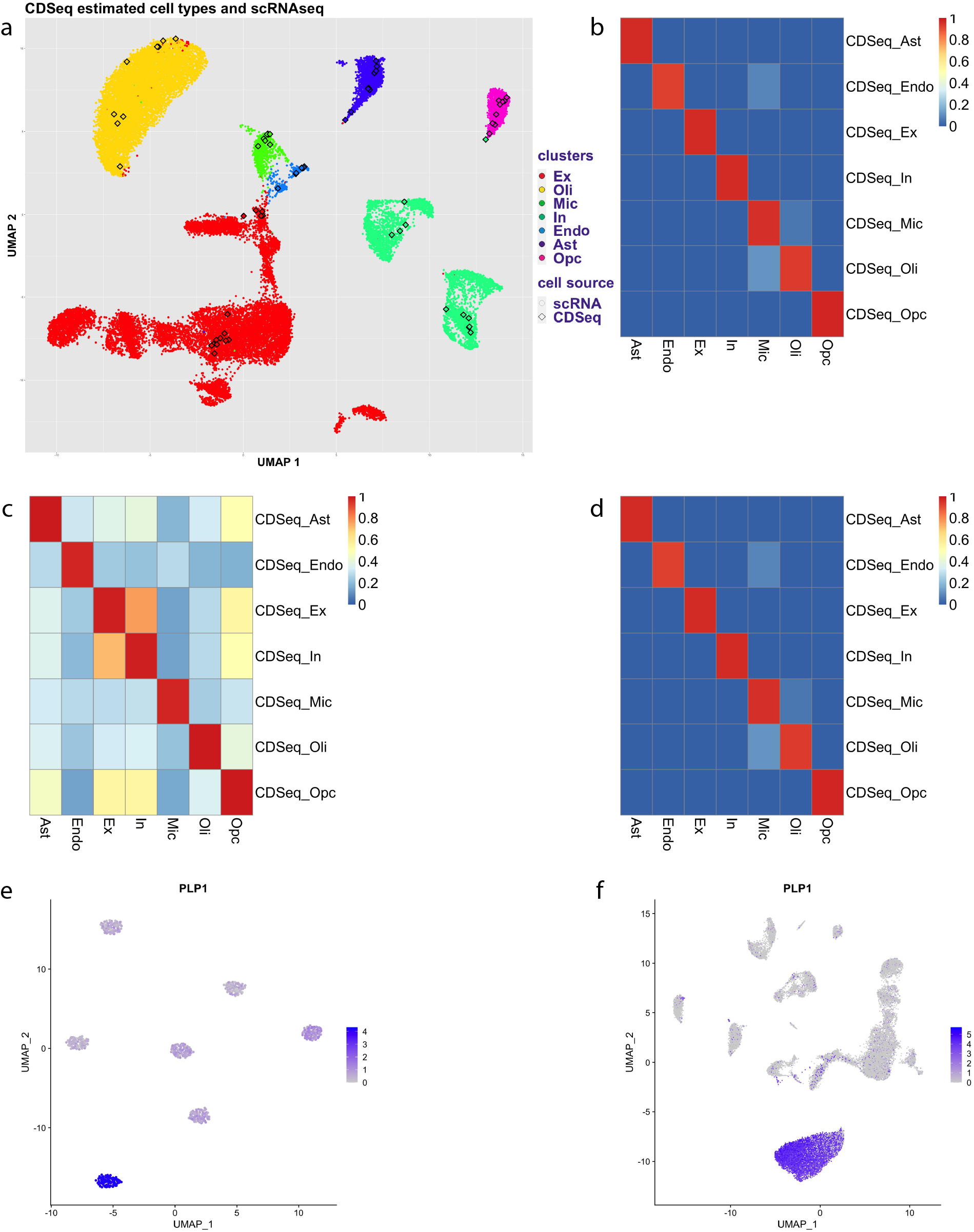
Annotation of CDSeq-identified cell types. CDSeqR was run with the Reduce-Recover option using 800 genes per block and 10 blocks. A reference scRNA-seq dataset was provided for cell type annotation using clustering analysis (a-b) and correlation analysis (c-d). (a) UMAP embeddings of reference scRNA-seq data (color dots) and pseudo single-cell data generated from the CDSeq-estimated cell-type specific gene expression profiles (color squares). Each distinct color in the UMAP plot denotes a cell type provided with the scRNA-seq annotation given in [21]. There are 7 major cell types provided with the scRNA-seq: excitatory neuron (Ex), inhibitory neuron (In), oligodendrocyte (Oli), microglia (Mic), endothelial (Endo), astrocyte (Ast) and oligodendrocyte progenitor cell (OPC). The deconvolution result used 7 as the number of cell types and for each CDSeq-estimated cell type, it generated 10 pseudo cells which correspond to a color square in the UMAP plot. The colors of the squares (i.e. CDSeq estimated cell types) were determined based on its annotation result. (b) correlations between the resulting CDSeqR estimated cell type proportions and true proportions; (c) correlations between the CDSeqR estimated cell-type-specific GEPs and scRNA-seq GEPs; (d) correlation of CDSeq-estimated cell type proportions with true proportions using correlation analysis (see Methods for details). When reference for cell annotation is not provided, CDSeq will perform clustering analysis on the pseudo single-cell data generated from the CDSeq-estimated cell-type specific gene expression profiles and perform differential gene expression analysis to identify the marker genes in each cluster using the Seurat software [25,26]. (e) Expression level of PLP1 (Oligodendrocyte marker [42]) in the clusters of CDSeq-estimated cell types (cell type number is 7); (f) Expression level of PLP1 in human brain prefrontal cortex scRNA-seq clusters [21]. See Supplementary Figure S10-S20 for more details.

## Results

We validated CDSeqR using both synthetic and cell mixtures [5] with known cell-type composition and cell-type-specific GEPs. For cell mixtures, the CDSeqR estimates using the Reduce and Recovery strategy matched the known cell-type-specific GEPs and sample-specific cell-type proportions (Fig. 2) while reducing computational time by about 96% (from about 120 min to 5 min) compared to the original implementation. In our previous work, we compared CDSeqR with seven other deconvolution methods [5]. Here, we employed a recently published benchmark pipeline [17] to simulate data for extensive comparisons with additional existing deconvolution methods. In total, we compared CDSeqR with 14 competing deconvolution algorithms [6,16,35,27–34] (Fig. 3 and Supplementary Fig. S6-S8) including linseed [6]. CDSeqR software and linseed both provided accurate cell-type proportion using similar amounts of computer time (Supplementary Fig. S1-S5). We further evaluated CDSeq and our new cell type annotation function by examining the results of using different number of cell types in the synthetic mixtures generated from scRNA-seq data (Supplementary Fig. S10-S16). In addition, we examined CDSeqR’s ability to detect immune cell subtypes in PBMC mixtures, a challenging problem referred to as deep deconvolution [18,28]. Specifically, we generated synthetic mixtures from 8 immune subtypes: B cells, Cytotoxic T cells, Helper T cells, Memory T cells, Monocytes, Naïve Cytotoxic T cells, Naïve T cells, and Natural killer T cells using scRNA-seq [19] and demonstrated that CDSeqR was able to identify all the immune cell subtypes and provided accurate estimates of cell type proportions (Supplementary Fig. S21 and S22). We further tested CDSeqR using GTEx [36] and TCGA [37] datasets and showed that the CDSeq identified cell types were consistent with the published single-cell RNA-Seq results [20,21] (Supplementary Fig. S23 and S24).

One input parameter for CDSeq function is the number of cell types to be deconvolved. The choice for the input is data dependent. For example, for a large dataset (e.g., >50), specifying a large number of cell type may help CDSeqR to detect all the major cell types due to the GEP heterogeneity even for a same cell type. Fortunately, CDSeqR allows the user to specify a range of possible cell type numbers as an input vector. For each choice in the vector, CDSeqR will perform computation in parallel and estimate the number of cell types based on the posterior distribution of the underlying model. In all *in silico* mixtures tested, CDSeqR provided accurate estimates for the number of cell types in the synthetic data.

In addition, one may use the cell type information in scRNA-seq data from the same tissue type as a guide. Here we showed that using synthetic mixture datasets generated from scRNA-seq data, the newly implemented cell type annotation function can successfully uncover the cell types (Fig. 4 and Supplementary fig. S10-24). Our simulation studies suggested that when the specified cell type number is larger than the true value, the cell type annotation function is able to identify and combine those cell types that are similar into one (Supplementary Fig. 10-16).

Another challenge for deconvolution is deep deconvolution [28], namely, to infer component subtypes from bulk tissues. For example, researchers are often interested in uncovering immune cell subtypes among major immune cell populations. In our previous work [18], we demonstrated that CDSeq was able to estimate immune cell subtypes from PBMC (peripheral blood mononuclear cell) data using microarray data. In this work, we further investigated CDSeq’s ability to infer immune cell subtypes using synthetic mixtures generated from scRNA-seq data [19]. Using a deconvolution benchmarking pipeline [17], we created 100 synthetic mixtures of various proportions of the 8 immune cell subtypes: B cells, Monocytes, Helper T cells, Natural killer cells, Cytotoxic T cells, Memory T cells, Naïve Cytotoxic T cells, and Naïve T cells. The highly-correlated GEPs of several of the cell types present major challenge for deconvolution methods. To test and validate the performance of CDSeqR, we ran CDSeqR by specifying a range of varying numbers of cell types from 2 to 40. CDSeqR can automatically identify similar cell types and merge them into one. CDSeqR was able to identify all 8 immune cell subtypes when the input number of the cell types was set to be greater than 10 (Fig 5; Supplementary Fig. S21 and S22).

**Figure 5.**
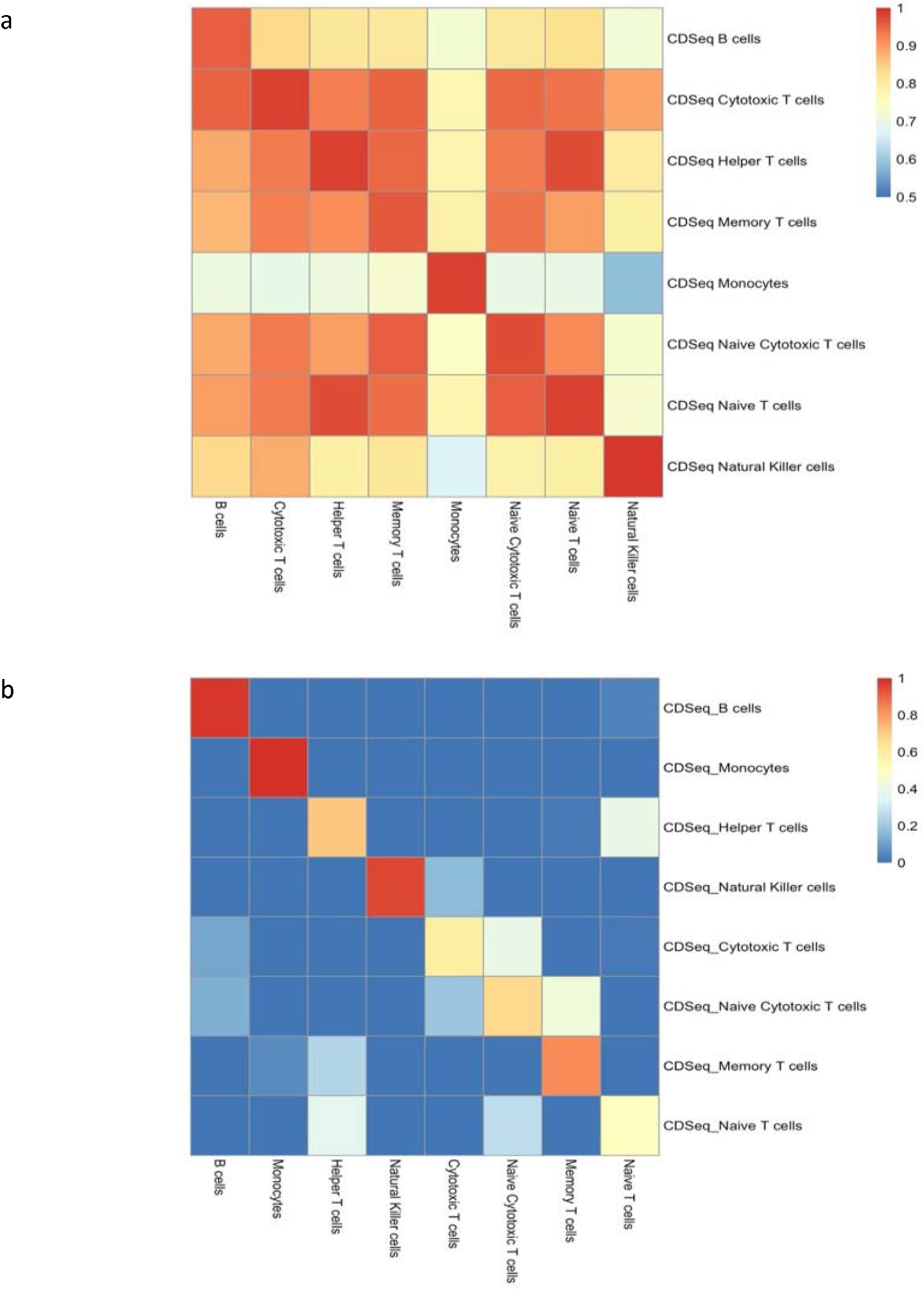
Results of deconvolution of PBMC immune cell subtype mixtures using Reduce-Recover strategy. The number of cell types was set to be 20. (a) Correlation between the CDSeq estimated cell-type-specific GEPs and true cell-type-specific GEPs; (b) Correlation between CDSeq estimated sample-specific cell-type proportions and true cell type proportions. Additional results for different specifications of the cell types are provided in Supplementary Fig. S21 and S22.

Besides the synthetic data, to further showcase the utility of CDSeqR and the newly implemented cell annotation function, we carried out deconvolution analysis of the RNA-seq data from human frontal cortex (209 samples) from the GTEx V8 [36]. We used publicly available brain single-cell RNA-seq data [21] as the reference for cell annotation. We set the number of cell types from 50 to 100 and ran CDSeqR with a dilution factor of 100 to facilitate computation. The result demonstrates that setting the number of cell types to 100 enabled detection of the major cell types in heterogenous brain samples whereas a smaller number (e.g., 50) failed to identify oligodendrocyte progenitor cell, a known major cell type (Supplementary Fig.S23). Setting the cell types to be 100 is reasonable as the brain is heterogenous and cell states are known to represent a continuum. This notion is consistent with the finding that, in brain prefrontal cortex [21,38], many of the major cell types consist of multiple subtypes. For example, excitatory neurons have more than ten subtypes [21,38]. In a second example, we deconvolved TCGA tumor samples without using any references. See Supplementary Fig.S24 for details.

## Discussion

CDSeq is a complete deconvolution method that uses gene expression data alone to estimate both sample-specific cell-type proportions and cell-type-specific gene expression profiles from RNA-seq data [5]. CDSeq was originally developed in MATLAB. In this work, we developed an R package, CDSeqR, for a broader group of users. Moreover, in the R implementation, we substantially improved CDSeq’s computational efficiency in both speed and memory usage. Specifically, we proposed a novel strategy – Reduce and Recover – to allow CDSeq to run on multiple processors simultaneously (parallelization). We further validated our CDSeqR package using data from both *in silico* and *in vitro* cell mixtures. We compared the performance of CDSeqR with that of 14 competing deconvolution method and showed that cell-type proportions estimated by CDSeqR are as or more accurate than those estimated by the competing methods. However, unlike many competing methods CDSeqR does not require reference GEPs for deconvolution and also provides cell-type-specific GEP estimates. We provided recommendations for parameter settings in Supplementary Table S1 and the parameters we used in our results in Supplementary Table S2.

In the R package, we also implemented a new cell-type assignment function. This new function takes advantage of the publicly available scRNA-seq datasets and use them as a reference to aid the annotation and visualization of the CDSeq estimated cell types. Specifically, CDSeqR transforms its estimated cell-type-specific GEPs into pseudo single-cell read counts (see Methods). Next, the two single-cell datasets (i.e. the external experimental scRNA-seq and the pseudo-scRNA-seq generated from CDSeq estimates) are combined and clustered using existing software such as Seurat [25,26]. Lastly, CDSeqR annotates the CDSeq cell types using either clustering analysis or correlation analysis (See Methods). In the case where no reference scRNA-seq dataset is available, CDSeqR identifies marker genes and uses them to aid cell type annotation.

The ongoing discovery of new cell subtypes [21,23,39] suggest that cell types may be better described as a continuum rather than as a sequence of discrete states. This recognition, consequently, brings challenges to deconvolution algorithms, especially for those that rely reference gene expression profiles [4,16,28,40]. The references will not fully capture the heterogeneity and the continuum of states exhibited by cell types in bulk tissues. Compared to existing deconvolution methods, CDSeq employs an unsupervised framework (does not require a reference panel) and can discover cell types in the data. Nonetheless, CDSeqR can only predict discrete cell types, not cell continuum. However, we showed that CDSeqR can do more than detect only major cell types but also can help users identify immune subtypes.

Since cell types represent a continuum, it is difficult to assign a value to the number of cell types present. This issue is analogous to determining the number of clusters from a clustering analysis. Nonetheless, we suggest that, in practice, one may set the number of cell types to be a vector that covers a range of plausible values. For example, set parameter cell_type_number = seq(20,100,10)) will allow CDSeq to try the number of cell types from 20 to 100 with an increment of 10. One can then visualize the clustering results with experimental scRNA-seq data using the cell-type annotation function and also consider the values of log posterior reported by CDSeq as a reference to determine the appropriate number of cell types. Importantly, the underlying biology would need to be considered when determining the number cell types in the data.

## Conclusion

We developed an R package for our complete deconvolution method [5] and implemented new functions to improve computational efficiency. Our method employs an unsupervised learning framework and requires no reference profile for the deconvolution procedure. We provided multiple ways to help identify CDSeq-estimated cell types after deconvolution. When a reliable single cell reference dataset is available, one can annotate deconvolved cell types using correlation analysis or clustering analysis. In the case where a single cell reference is not available, CDSeqR will identify marker genes for each deconvolved cell type to help users annotate the identities of those cell types. Our method is also able to infer the number of cell types from bulk RNA-seq measurements. In summary, we believe that our CDSeqR package will be a valuable tool for computationally deciphering sample heterogeneity using bulk RNA-Seq data.

## Abbreviations

TCGA: The Cancer Genome Atlas
GTEx: The Genotype-Tissue Expression (GTEx) project

## Availability and requirements

Project name: CDSeq R package

Project home page: https://github.com/kkang7/CDSeq_R_Package

Operating system: Platform independent

Programming language: R, C++, C.

Other requirements: None

License: GPL-3

Any restrictions to use by non-academics: None

## Declarations

Ethics approval and consent to participate

Not Applicable.

## Consent for publication

Not Applicable.

## Availability of data and materials

Source code are available on Github (https://github.com/kkang7/CDSeq_R_Package) and CRAN (https://cran.r-project.org/web/packages/CDSeq/index.html).

## Authors’ contributions

LL, KK and DU conceived the project. KK designed the new strategy for parallelization and developed the new function for annotating CDSeq-estimated cell types. KK and DH implemented the R package. KK, DH and YL performed numerical testing and validations. All authors contributed to the writing of the manuscript. LL guided the research. All authors read and approved the final manuscript.

## Acknowledgements

The authors thank the NIEHS Office of Scientific Computing for computing support and the NIEHS Epigenomics and DNA Sequencing Core Facility for RNA-seq analysis of the cell mixture samples.

## Additional files

File name: Additional file 1

Title: Supplementary Note for CDSeqR

Description: Supplementary methods, tables and figures

## Reference

1. Demaria O, Cornen S, Daëron M, Morel Y, Medzhitov R, Vivier E. Harnessing innate immunity in cancer therapy. Nature. 2019.

2. Zheng C, Zheng L, Yoo JK, Guo H, Zhang Y, Guo X, et al. Landscape of Infiltrating T Cells in Liver Cancer Revealed by Single-Cell Sequencing. Cell. 2017;

3. Shen-Orr SS, Tibshirani R, Khatri P, Bodian DL, Staedtler F, Perry NM, et al. Cell type--specific gene expression differences in complex tissues. Nat Methods. 2010;7(4):287–9.

4. Newman AM, Steen CB, Liu CL, Gentles AJ, Chaudhuri AA, Scherer F, et al. Determining cell type abundance and expression from bulk tissues with digital cytometry. Nat Biotechnol. 2019;

5. Kang K, Przytycka TM, Meng Q, Shats I, Umbach DM, Li M, et al. CDSeq: A novel complete deconvolution method for dissecting heterogeneous samples using gene expression data. PLOS Comput Biol [Internet]. 2019 Dec 2; Available from: https://doi.org/10.1371/journal.pcbi.1007510

6. Zaitsev K, Bambouskova M, Swain A, Artyomov MN. Complete deconvolution of cellular mixtures based on linearity of transcriptional signatures. Nat Commun. 2019;

7. Shen-Orr SS, Gaujoux R. Computational deconvolution: extracting cell type-specific information from heterogeneous samples. Curr Opin Immunol. 2013;25(5):571–8.

8. Erkkilä T, Lehmusvaara S, Ruusuvuori P, Visakorpi T, Shmulevich I, Lähdesmäki H. Probabilistic analysis of gene expression measurements from heterogeneous tissues. Bioinformatics. 2010;

9. Qiao W, Quon G, Csaszar E, Yu M, Morris Q, Zandstra PW. PERT: a method for expression deconvolution of human blood samples from varied microenvironmental and developmental conditions. PLoS Comput Biol. 2012;8(12):e1002838.

10. Gong T, Szustakowski JD. DeconRNASeq: a statistical framework for deconvolution of heterogeneous tissue samples based on mRNA-Seq data. Bioinformatics. 2013;29(8):1083–5.

11. Zhong Y, Wan YW, Pang K, Chow LML, Liu Z. Digital sorting of complex tissues for cell type-specific gene expression profiles. BMC Bioinformatics. 2013;

12. Gaujoux R, Seoighe C. Semi-supervised Nonnegative Matrix Factorization for gene expression deconvolution: a case study. Infect Genet Evol. 2012;12(5):913–21.

13. Li Y, Xie X. A mixture model for expression deconvolution from RNA-seq in heterogeneous tissues. BMC Bioinformatics. 2013;14(5):S11.

14. Ahn J, Yuan Y, Parmigiani G, Suraokar MB, Diao L, Wistuba II, et al. DeMix: Deconvolution for mixed cancer transcriptomes using raw measured data. Bioinformatics. 2013;

15. Newman AM, Liu CL, Green MR, Gentles AJ, Feng W, Xu Y, et al. Robust enumeration of cell subsets from tissue expression profiles. Nat Methods. 2015;

16. Wang X, Park J, Susztak K, Zhang NR, Li M. Bulk tissue cell type deconvolution with multi-subject single-cell expression reference. Nat Commun. 2019;

17. Avila Cobos F, Alquicira-Hernandez J, Powell JE, Mestdagh P, De Preter K. Benchmarking of cell type deconvolution pipelines for transcriptomics data. Nat Commun. 2020;

18. Kang K, Meng Q, Shats I, Umbach DM, Li M, Li Y, et al. CDSeq: A novel complete deconvolution method for dissecting heterogeneous samples using gene expression data. PLoS Comput Biol. 2019;

19. Zheng GXY, Terry JM, Belgrader P, Ryvkin P, Bent ZW, Wilson R, et al. Massively parallel digital transcriptional profiling of single cells. Nat Commun. 2017;

20. Puram S V., Tirosh I, Parikh AS, Patel AP, Yizhak K, Gillespie S, et al. Single-Cell Transcriptomic Analysis of Primary and Metastatic Tumor Ecosystems in Head and Neck Cancer. Cell. 2017;

21. Mathys H, Davila-Velderrain J, Peng Z, Gao F, Mohammadi S, Young JZ, et al. Single-cell transcriptomic analysis of Alzheimer’s disease. Nature. 2019;

22. Griffiths TL, Steyvers M. Finding scientific topics. Proc Natl Acad Sci. 2004;

23. Pastushenko I, Brisebarre A, Sifrim A, Fioramonti M, Revenco T, Boumahdi S, et al. Identification of the tumour transition states occurring during EMT. Nature. 2018;

24. Han X, Zhou Z, Fei L, Sun H, Wang R, Chen Y, et al. Construction of a human cell landscape at single-cell level. Nature. 2020;

25. Butler A, Hoffman P, Smibert P, Papalexi E, Satija R. Integrating single-cell transcriptomic data across different conditions, technologies, and species. Nat Biotechnol. 2018;

26. Stuart T, Butler A, Hoffman P, Hafemeister C, Papalexi E, Mauck WM, et al. Comprehensive Integration of Single-Cell Data. Cell. 2019;

27. Chambers JM, Hastie TJ. Statistical models in S. Statistical Models in S. 2017.

28. Newman AM, Liu CL, Green MR, Gentles AJ, Feng W, Xu Y, et al. Robust enumeration of cell subsets from tissue expression profiles. Nat Methods. 2015;12(5):453.

29. Du R, Carey V, Weiss ST. DeconvSeq: Deconvolution of cell mixture distribution in sequencing data. Bioinformatics. 2019;

30. Hao Y, Yan M, Heath BR, Lei YL, Xie Y. Fast and robust deconvolution of tumor infiltrating lymphocyte from expression profiles using least trimmed squares. PLoS Comput Biol. 2019;

31. Riplley B, Venables B, Bates DM, Firth D, Hornik K, Gebhardt A. Package “MASS”. Support Functions and Datasets for Venables and Ripley’s MASS. Document freely available on the internet at: http://www.r-project.org. 2018.

32. Altboum Z, Steuerman Y, David E, Barnett-Itzhaki Z, Valadarsky L, Keren-Shaul H, et al. Digital cell quantification identifies global immune cell dynamics during influenza infection. Mol Syst Biol. 2014;

33. Friedman J, Hastie T, Tibshirani R. Regularization paths for generalized linear models via coordinate descent. J Stat Softw. 2010;

34. Jew B, Alvarez M, Rahmani E, Miao Z, Ko A, Garske KM, et al. Accurate estimation of cell composition in bulk expression through robust integration of single-cell information. Nat Commun. 2020;

35. Dong M, Thennavan A, Urrutia E, Li Y, Perou CM, Zou F, et al. SCDC: bulk gene expression deconvolution by multiple single-cell RNA sequencing references. Brief Bioinform. 2021;

36. Aguet F, Barbeira AN, Bonazzola R, Brown A, Castel SE, Jo B, et al. The GTEx Consortium atlas of genetic regulatory effects across human tissues. Science (80-). 2020;

37. Weinstein JN, Collisson EA, Mills GB, Shaw KRM, Ozenberger BA, Ellrott K, et al. The cancer genome atlas pan-cancer analysis project. Nature Genetics. 2013.

38. Gandal MJ, Zhang P, Hadjimichael E, Walker RL, Chen C, Liu S, et al. Transcriptome-wide isoform-level dysregulation in ASD, schizophrenia, and bipolar disorder. Science (80-). 2018;

39. Trapnell C. Defining cell types and states with single-cell genomics. Genome Res. 2015;25(10):1491–8.

40. Hunt GJ, Freytag S, Bahlo M, Gagnon-Bartsch JA. Dtangle: Accurate and robust cell type deconvolution. Bioinformatics. 2019;

41. Ding J, Adiconis X, Simmons SK, Kowalczyk MS, Hession CC, Marjanovic ND, et al. Systematic comparison of single-cell and single-nucleus RNA-sequencing methods. Nat Biotechnol. 2020;

42. McKenzie AT, Wang M, Hauberg ME, Fullard JF, Kozlenkov A, Keenan A, et al. Brain Cell Type Specific Gene Expression and Co-expression Network Architectures. Sci Rep. 2018;

