## Supplemental materials for "CDSeqR: fast complete deconvolution for gene expression data from bulk tissues"

**Supplementary Methods**

### 1 A dimension *Reduce-Recover* strategy using parallel computing

A major practical challenge of CDSeq [1] is the computational time. The Gibbs sampler we employed for parameter estimation requires storage of several vectors with very large sizes; their lengths equal to the total number of reads from all samples. Depending on the datasets, the lengths of such vectors typically range from millions to billions. As a result, a large amount of memory is required for parameter estimation, dramatically diminishing the utility of CDSeq. In this work, we proposed a novel strategy for alleviating the issue. We call it a dimension *Reduce-Recover* strategy. The schematic approach is presented in Fig. 1. We provide the details of this approach in the following.

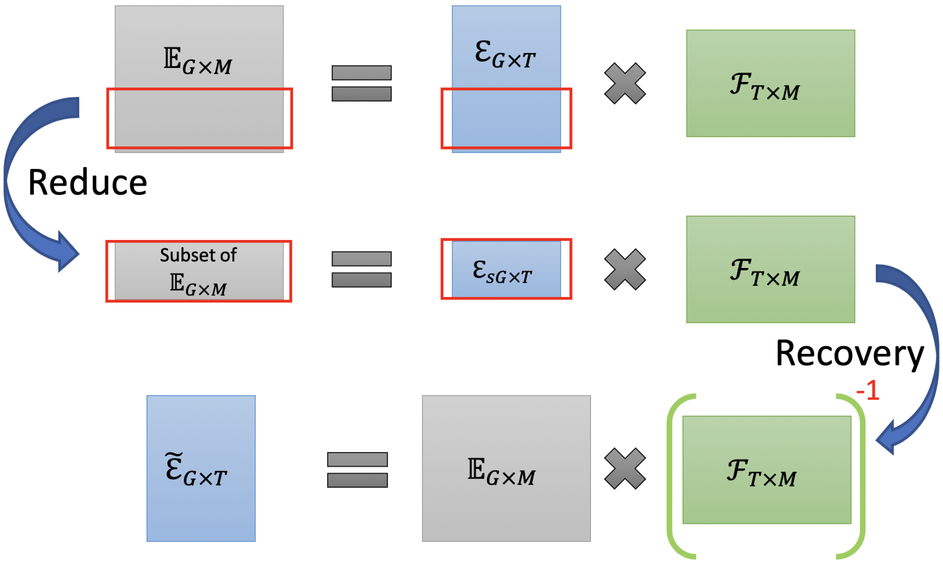


Figure 1: Dimension *Reduce-Recover* strategy.

Let $\mathcal{E}_{G\times T}$ denote the matrix of cell-type-specific gene expression profiles, where $G$ is the total number of genes and $T$ is the number of cell types. Each column in $\mathcal{E}_{G\times T}$ represents a Dirichlet random variable, i.e., $\sum_{i=1}^{G} \mathcal{E}_{i,t} = 1, \forall t \in \{1, \cdots, T\}$. We denote the cell-type fractions of all samples using $\mathcal{F}_{T\times M}$, where $M$ is the number of tissue samples. Each column in $\mathcal{F}_{T\times M}$ also represents a Dirichlet random variable, i.e., $\sum_{t=1}^{T} \mathcal{F}_{t,j} = 1, j = 1, \cdots, M$. Next, let $\mathbb{E}_{G\times M}$ denote the gene expression profile of bulk samples and its column is again a Dirichlet random variable. Then, based on our model, we have that

$$\mathcal{E}_{G\times T} \cdot \mathcal{F}_{T\times M} = \mathbb{E}_{G\times M}. \tag{1}$$

In a bulk tissue sample, for any gene, the mapped reads are a mixture of reads from its component cell types. We assume that the same cell type will always have the same GEP profile in all samples. Therefore, for each gene, the cell-type-specific contribution to the expression remains the same.

Specifically, based on (1), it is clear that, in sample $j$,

$$\mathbb{E}_{i,j} = \sum_{t=1}^{T} \mathcal{E}_{i,t}\mathcal{F}_{t,j},$$

where $\mathcal{F}_{t,j}$ is the contribution to the expression of gene $i$ from cell type $t$, and $\mathcal{F}_{t,j}$ does not vary with respect to different genes in a sample. This observation leads to the dimension *reduce-recover* strategy. It contains two steps: first, we can randomly pick a subset of genes, instead of using all the genes, to perform the parameter estimation. The cell-type fraction $\mathcal{F}_{T\times M}$ should be invariant to the dimension reduction on genes.

One output will then be an estimate of $\mathcal{F}_{T\times M}$, based on a subset of genes. That estimate should approximate the one based on the full set of genes. Another output will be an estimate of $\mathcal{E}_{\tilde{G},T}$ where $\tilde{G} \leq G$. Then, the second step is to recover $\mathcal{E}_{G,T}$. We propose to perform the following: first, estimate $\mathbb{E}_{G\times M}$ by normalizing the read count bulk RNA-seq data; second, perform

$$\tilde{\mathcal{E}}_{G,T} = \mathbb{E}_{G\times M} \cdot \mathcal{F}^{-1}_{T\times M},$$

where $\mathcal{F}^{-1}_{T\times M} = (\mathcal{F}^{T}_{T\times M}\mathcal{F}_{T\times M})^{-1}\mathcal{F}^{T}_{T\times M}$ is the left inverse of $\mathcal{F}_{T\times M}$; third, estimate $\mathcal{E}_{G\times T}$ by normalizing the columns of $\tilde{\mathcal{E}}_{G,T}$, namely,

$$\mathcal{E}_{i,j} = \frac{|\tilde{\mathcal{E}}_{i,j}|}{\sum\limits_{j=1}^{G} |\tilde{\mathcal{E}}_{i,j}|},$$

where $|\cdot|$ denotes the absolute value. Of note, since it is unlikely in practice that any two heterogeneous samples will have the same fraction of component cell types, the cell-type fraction matrix $\mathcal{F}_{T\times M}$ will have full column rank, thereby, guaranteeing the existence of its left inverse. We used

$ginv()$ function in MASS package for computing the matrix inverse. The $ginv()$ procedure calculates the Moore-Penrose inverse by using a singular value decomposition approach.

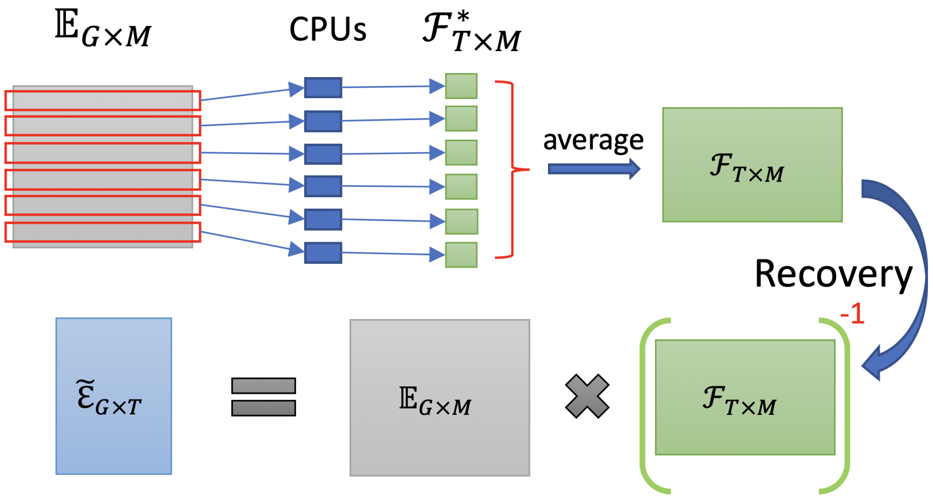


Figure 2: Dimension *Reduce-Recover* strategy using parallel computing.

### 2 *In vitro* mixtures

The *in vitro* mixtures we used for validations in main text Fig.2 were the same data we used in [1]. In brief, total mRNA was prepared from four cell types: Namalwa (Burkitt's lymphoma), Hs343T (fibroblast line derived from a mammary gland adenocarcinoma), hTERT-HME1 (normal mammary epithelial cells immortalized with hTERT), and MCF7 (estrogen receptor positive breast cancer cell line). A detailed description of the data generation can be found in Supplementary material in [1].

### 3 Benchmarking

We employed benchmarking pipeline developed in [2] to carry out comparisons with 14 existing deconvolution methods [3–14]. The benchmarking pipeline takes scRNA-seq and divide it into testing data and training data. Then, it constructs synthetic mixtures using training data and use testing data as the reference for deconvolution methods. In the constructed mixtures, the cell-type-specific GEPs and sample-specific cell-type proportions are known and are used for accessing the performance of different deconvolution methods. We used three different scRNA-seq datasets for benchmarking the performance: human prefrontal cortex [15], PBMC [16] and head and neck cancer [17]. Note that most of the methods are reference-based methods and rely on reference profile, i.e. cell-type-specific GEPs, to provide estimations of sample-specific cell-type proportions. We emphasis that CDSeqR is a complete deconvolution method and does not require any reference GEPs and outputs estimates for cell-type-specific GEPs. We show comparisons of cell type proportion estimations with other methods.

### Supplementary Figures and Tables

| Parameters | Recommended setting |
|---|---|
| *beta* | $0.5$ |
| *alpha* | $5$ |
| *mcmc_iteration* | $700 \sim 2000$ |
| *dilution_factor* | $2 \sim 10$ |
| *gene_subset_size* | $> 200$ |
| *block_number* | $> 5$ |

Table S1: Parameter setting recommendation.

| Settings in results | | | | | | | |
|---|---|---|---|---|---|---|---|
| Data | Mode | Cell type number | MCMC iteration | Dilution factor | Block number | Block size | Figure |
| *in vitro* | CDSeqRR | 4 | 1000 | 10 | 50 | 1000 | S1, S2 |
| | CDSeq | 4 | 1000 | 10 | NA | NA | S1, S2 |
| *in silico* | CDSeqRR | 6 | 1000 | 1 | 50 | 1000 | S3, S4 |
| | CDSeq | 6 | 1000 | 1 | NA | NA | S3, S4 |
| PBMC | CDSeqRR | 5 | 2000 | 10 | 10 | 800 | S6 |
| | CDSeq | 5 | 700/1000/2000/3000 | 10 | 1 | NA | S6 |
| | CDSeqRR | 2:20 | 1000 | 10 | 10 | 800 | S6 |
| PBMC immune | CDSeqRR | 2:40 | 1000 | 1 | 10 | 800 | S21, S22 |
| | CDSeq | 2:40 | 1000 | 1 | NA | NA | S21, S22 |
| Brain | CDSeqRR | 7 | 1000 | 1 | 10 | 800 | S7 |
| | CDSeq | 7 | 700/1000/2000 | 1 | NA | NA | S7 |
| | CDSeqRR | 2:20 | 1000 | 1 | 10 | 800 | S7 |
| Tumor | CDSeqRR | 7 | 2000 | 1 | 10 | 400 | S8 |
| | CDSeq | 7 | 700/1000/2000/3000 | 1 | NA | NA | S8 |
| GTEx | CDSeq | seq(5,100,5) | 1000 | 100 | NA | NA | S23 |
| TCGA | CDSeq | 10:40 | 1000 | 10 | NA | NA | S24 |

Table S2: Parameter settings in the results.

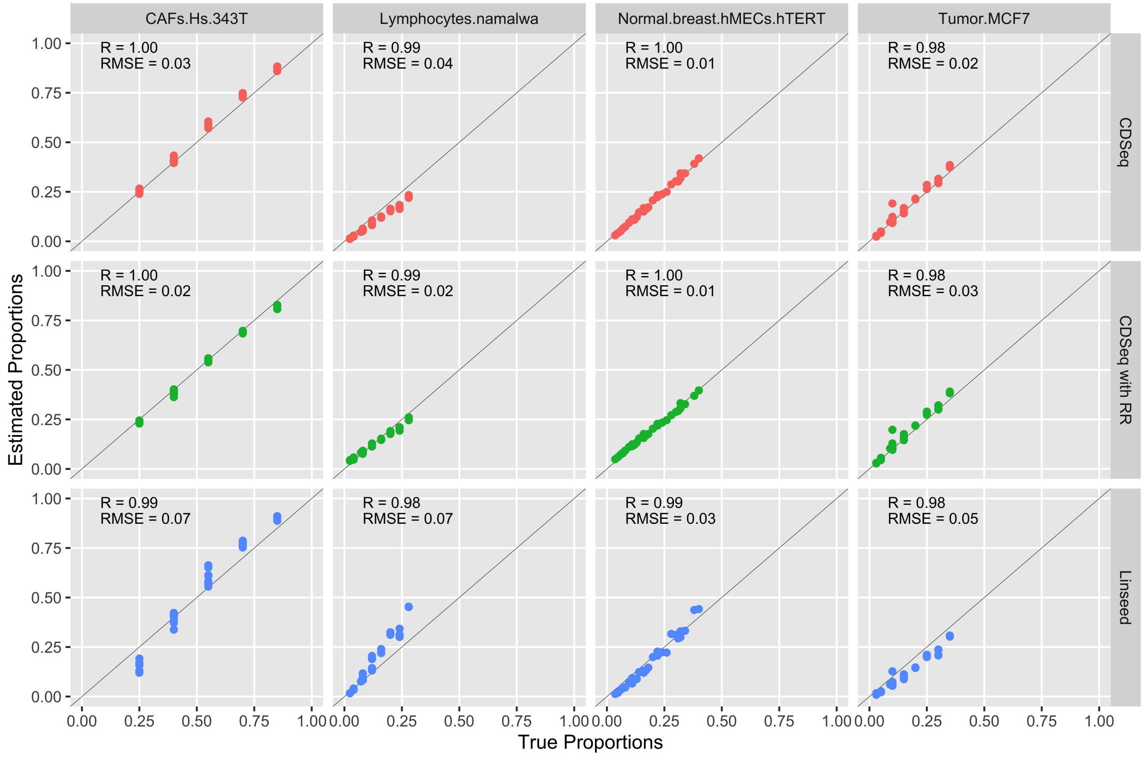


Figure S1: Sample-specific cell-type proportion estimation on 32 *in vitro* mixtures of four cell types. CDSeq used 50 blocks with 1,000 genes randomly sampled in each block. 1000 MCMC iterations were performed for both CDSeq and CDSeq with reduce-recover (RR) (first two rows). For Linseed [18], top 10,000 genes were used. We compared estimated proportions with true proportions and reported correlation (R) and root-mean-square-error (RMSE). Each point represents a sample. The black diagonal line is the $x = y$ line.

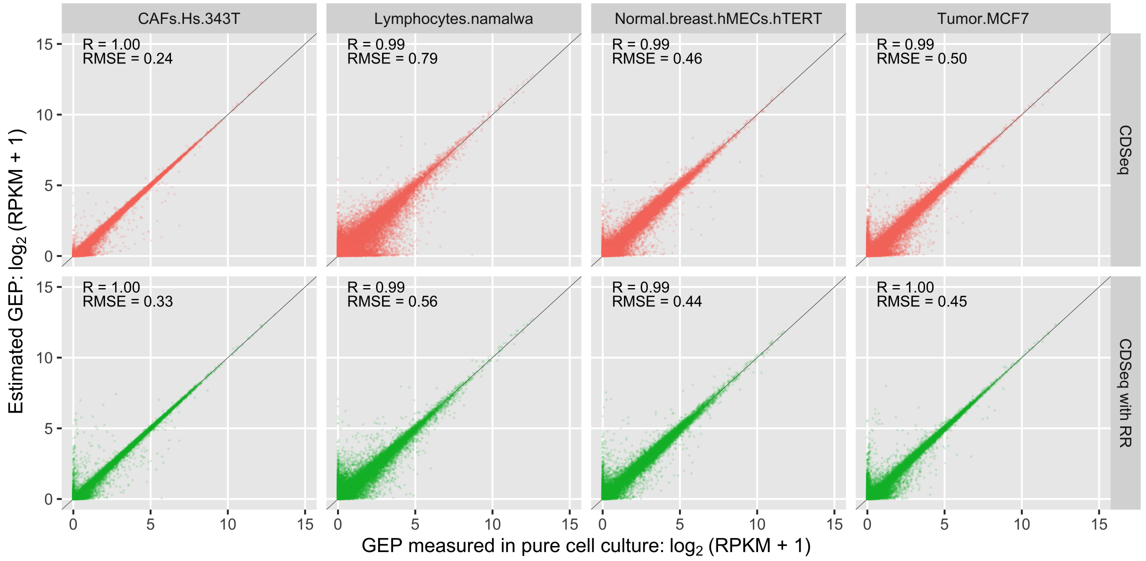


Figure S2: Cell-type-specific GEP (RPKM) estimation on 32 in vitro mixtures of four cell types. We compared the estimated GEPs with the true GEPs and reported correlation (R) and root-mean-square-error (RMSE). Each point represents a gene. Total number of genes is 19653. True and estimated GEP are normalized as RPKM. Linseed does not provide cell-type-specific GEPs' estimates. The black diagonal line is the $x = y$ line.

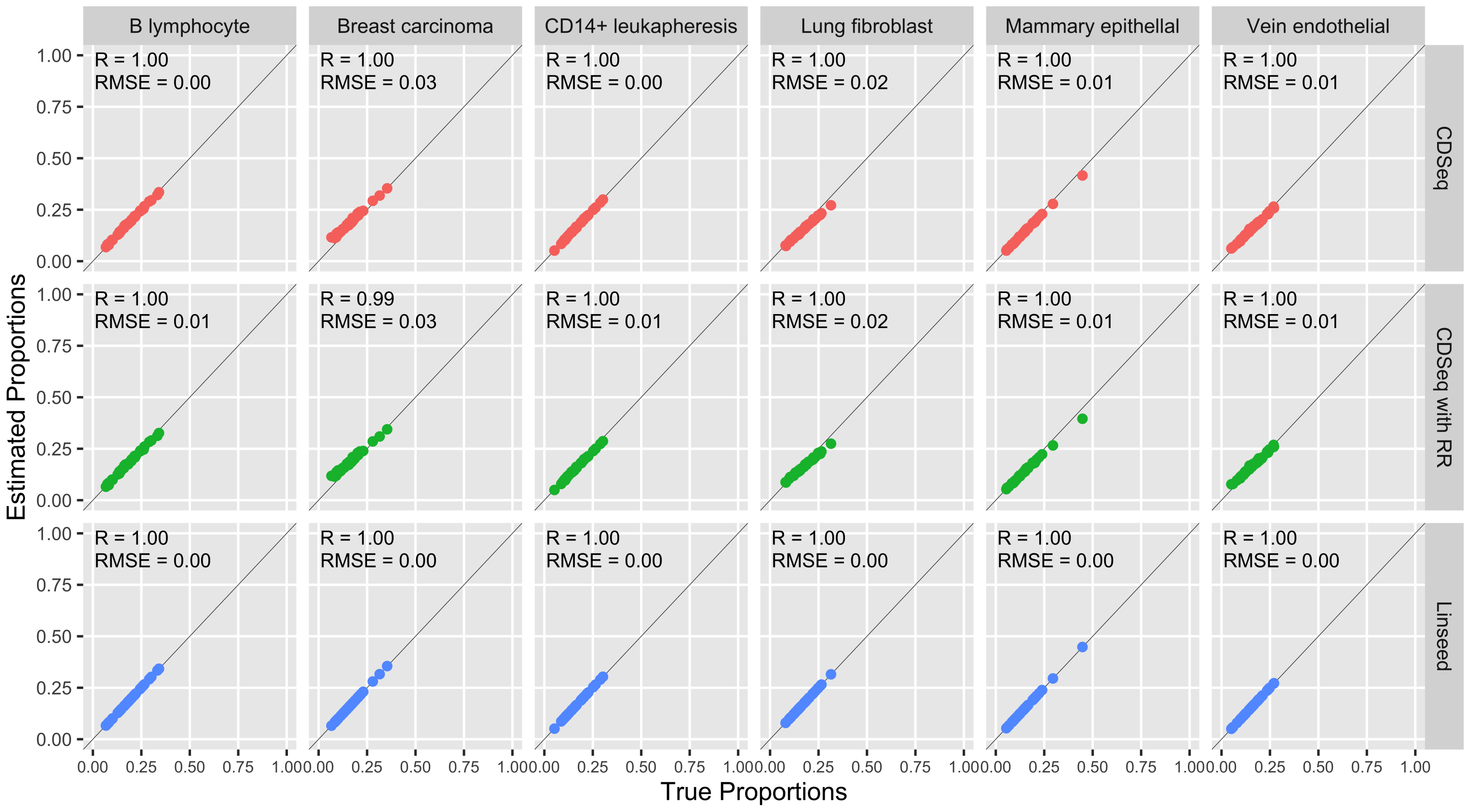


Figure S3: Sample-specific cell-type proportion estimates based on 40 *in silico* mixtures of six cell types. See Figure S1 legend for additional details. Each point represents a sample. The black diagonal line is the $x = y$ line.

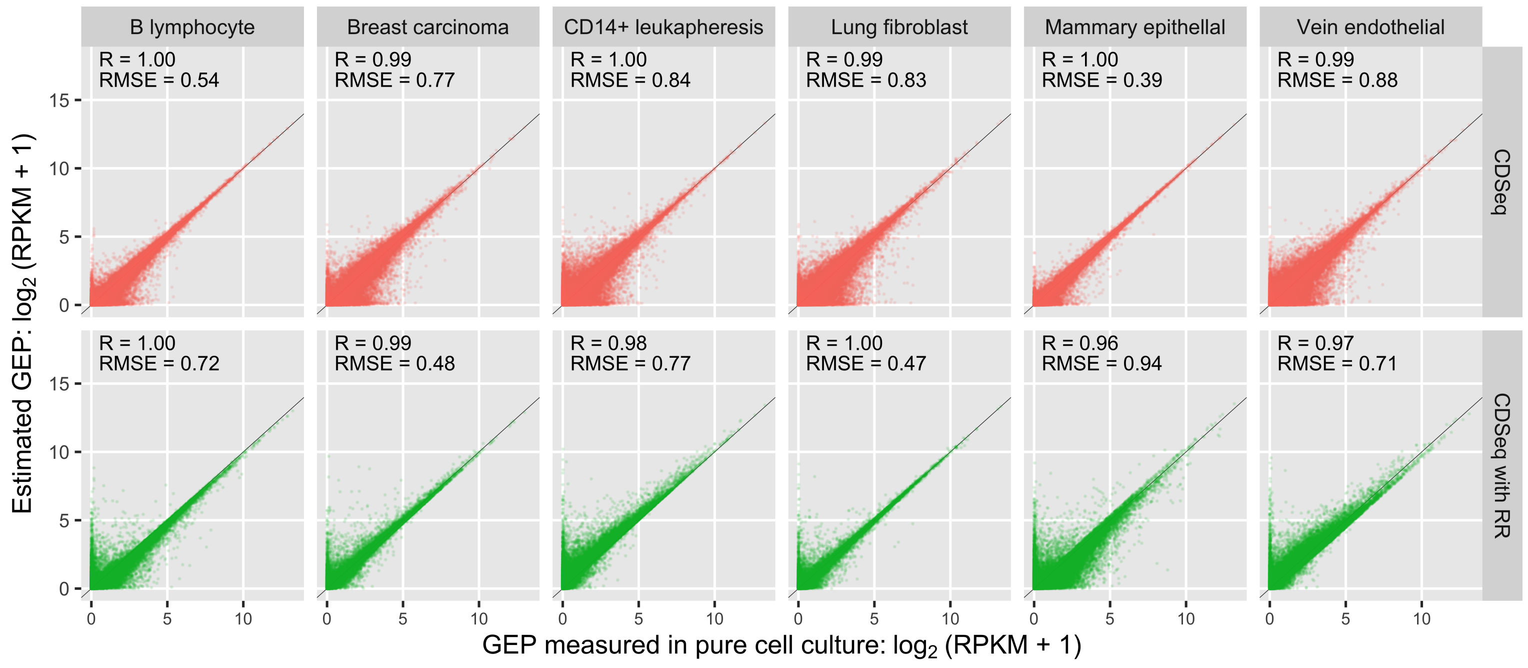


Figure S4: Cell-type-specific GEP (RPKM) estimates based on 40 *in silico* mixtures of six cell types. We compared the estimated GEPs with the true GEPs and reported correlation (R) and root-mean-square-error (RMSE). Each point represents a gene. Total number of genes is 22498. True and estimated GEP are normalized as RPKM. Linseed does not provide cell-type-specific GEPs' estimates. The black diagonal line is the $x = y$ line.

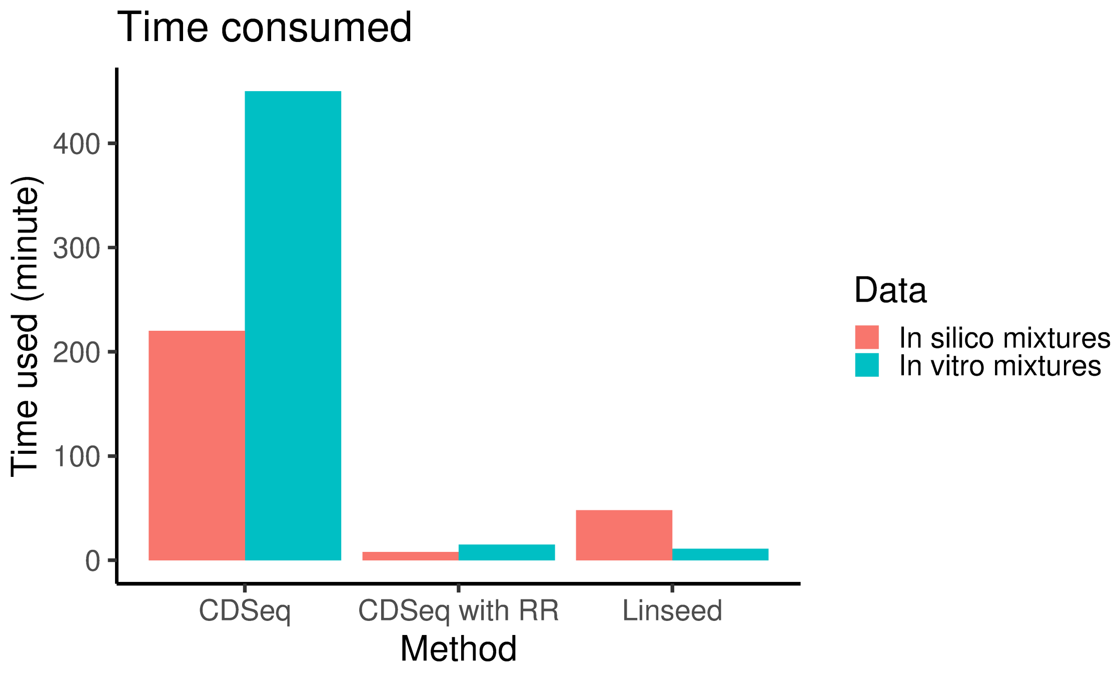


Figure S5: Running time comparisons. CDSeq with RR (Reduce-Recover) took about 8 minutes and 15 minutes using 50 cores (Intel(R) Xeon(R) Platinum 8160 CPU @ 2.10GHz) for *in silico* and *in vitro* mixtures, respectively; Linseed took about 45 minutes and 11 minutes respectively.

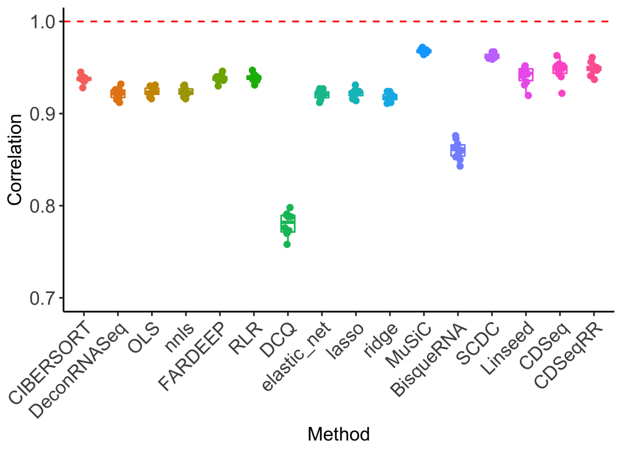


(a) Comparison of correlations between true cell type proportions and cell type proportions estimated by different deconvolution methods.

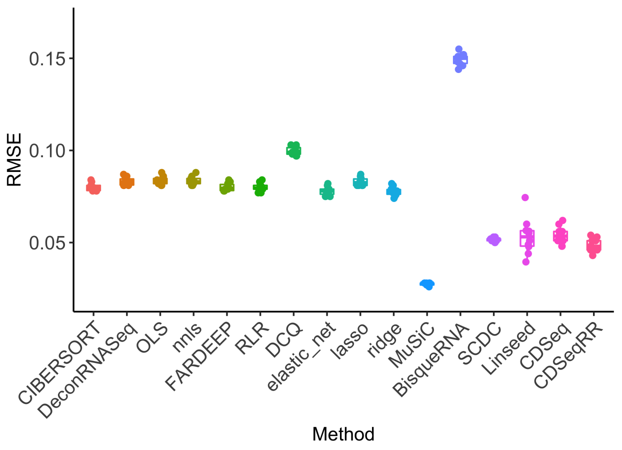


(b) Comparison of RMSE between true cell type proportions and cell type proportions estimated by different deconvolution methods.

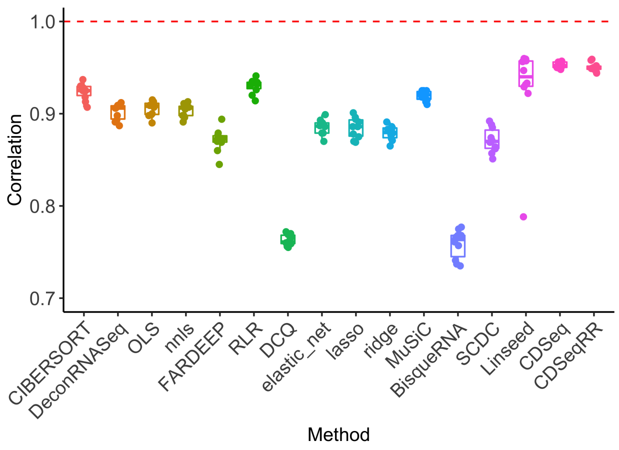


(c) Comparison of correlations between true cell type proportions and cell type proportions estimated by different deconvolution methods. Training and testing data are from two datasets.

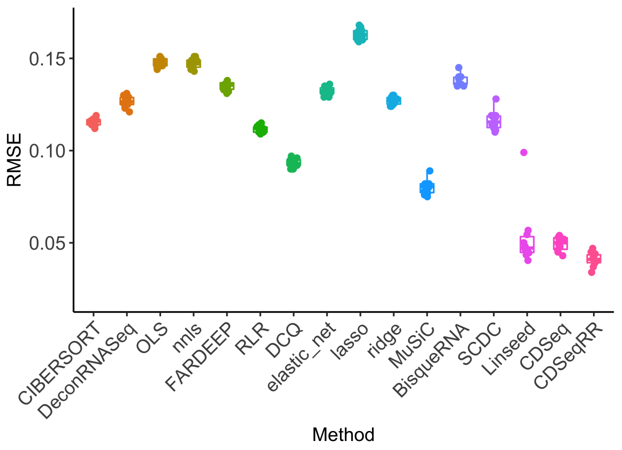


(d) Comparison of RMSE between true cell type proportions and cell type proportions estimated by different deconvolution methods. Training and testing data are from two datasets.

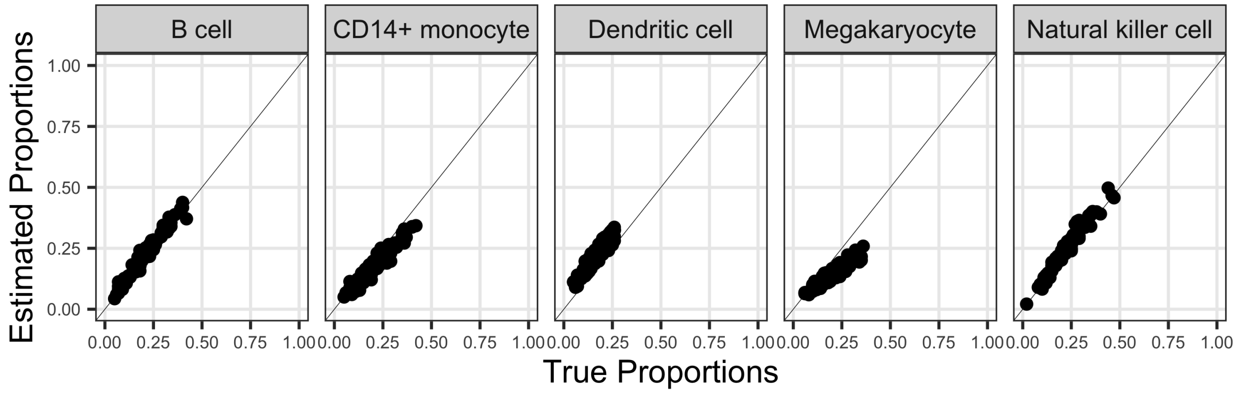


(e) CDSeq estimated proportions versus true proportions.

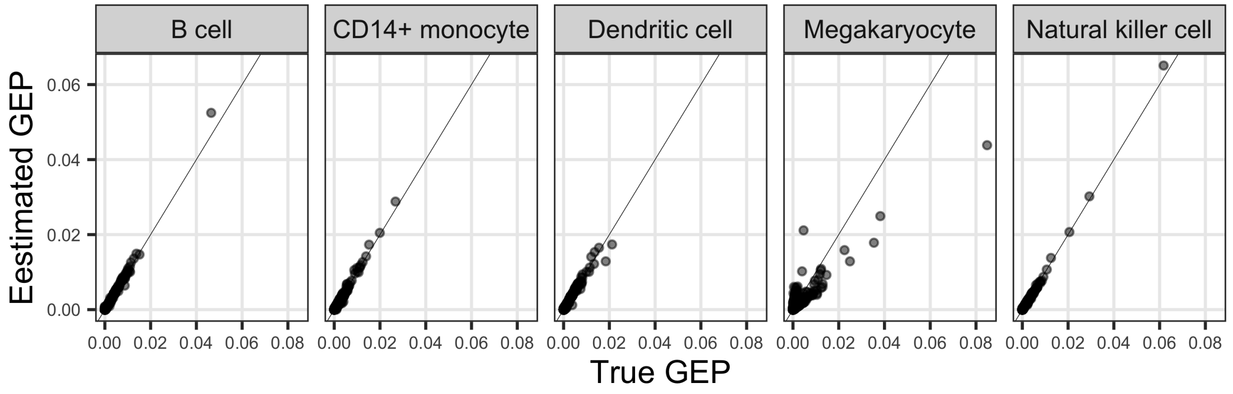


(f) CDSeq estimated GEPs versus true GEPs..

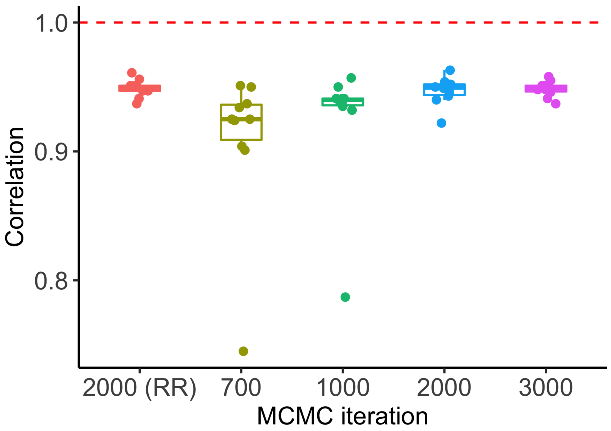


(g) Comparison of correlations using different MCMC runs.

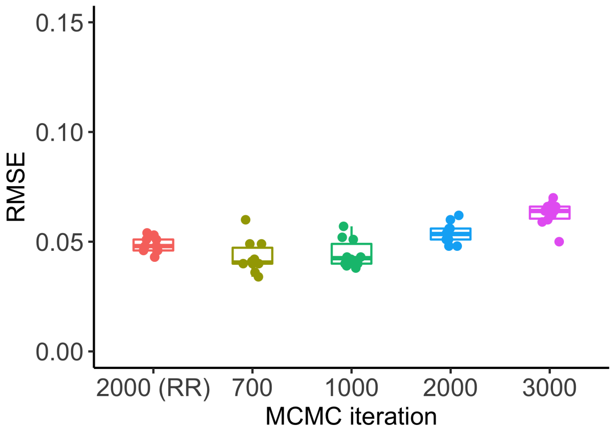


(h) Comparison of RMSEs using different MCMC runs.

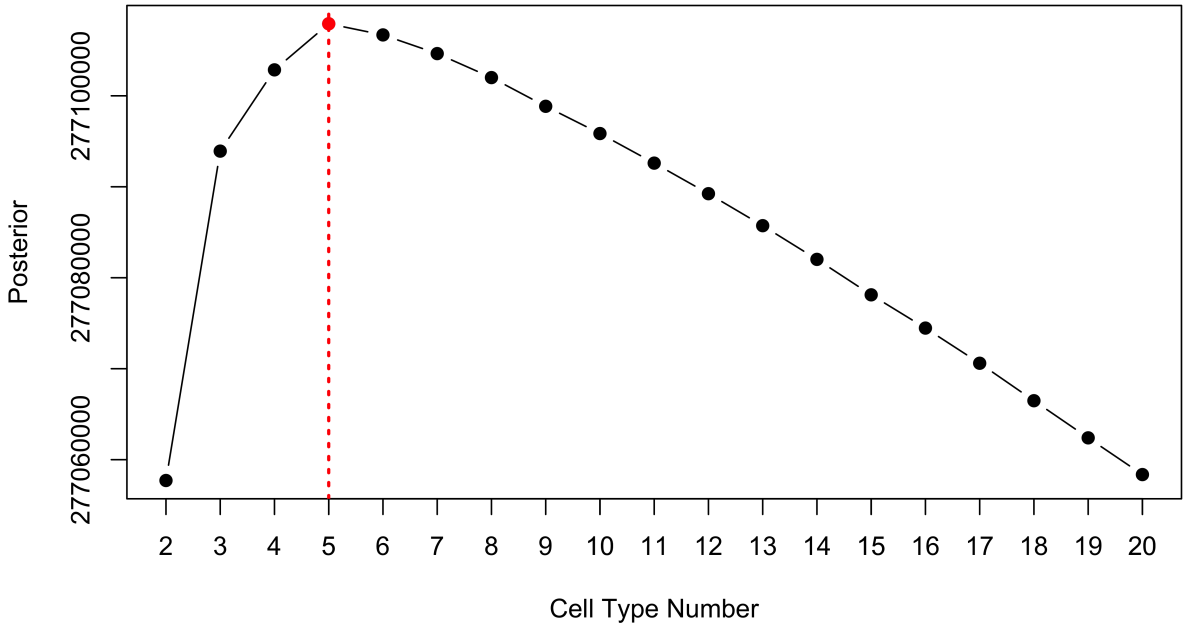


(i) CDSeq is able to accurately estimate the number of cell types in the synthetic mixtures.

Figure S6: Comparison of different deconvolution methods using synthetic mixtures generated by PBMC single cell data [16]. We employed the benchmarking pipeline developed in [2] to carry out comparisons using PBMC scRNA-seq data [16] and PBMC data from Seurat tutorial page. We can see that most methods performed reasonably well and CDSeq is among the top performers. It is worth noting that reference based methods is affected by the reference profile being used. As an illustration, we performed deconvolution using the reference data from a same scRNA-seq dataset as shown in Fig.S6a and Fig.S6b and using the reference data from a different dataset (Fig. S6cand Fig.S6d). It is noted that most reference based methods resulted in a lower correlation and higher RMSE whereas CDSeq was not affected. We further tested the effect of different MCMC runs and show that more MCMC runs will provide more accurate results as shown in Fig.S6g and Fig.S6h. In addition, CDSeq is able to estimate the number of cell types in the bulk sample as indicated in Fig.S6i.

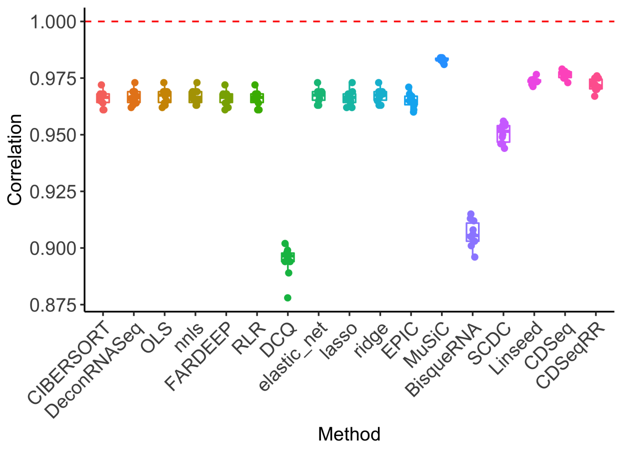


(a) Correlation comparisons between different deconvolution methods.

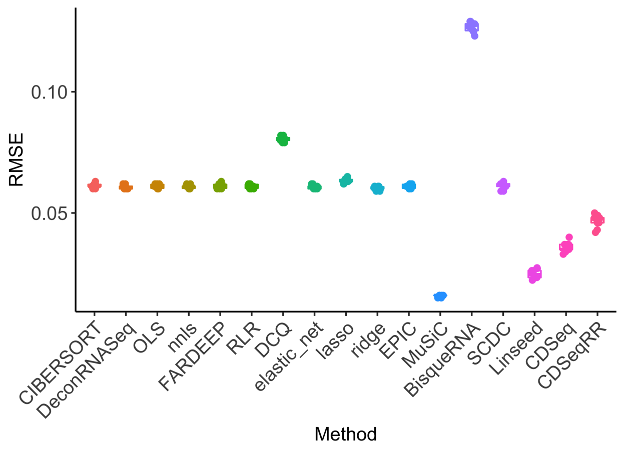


(b) RMSE comparisons between different deconvolution methods.

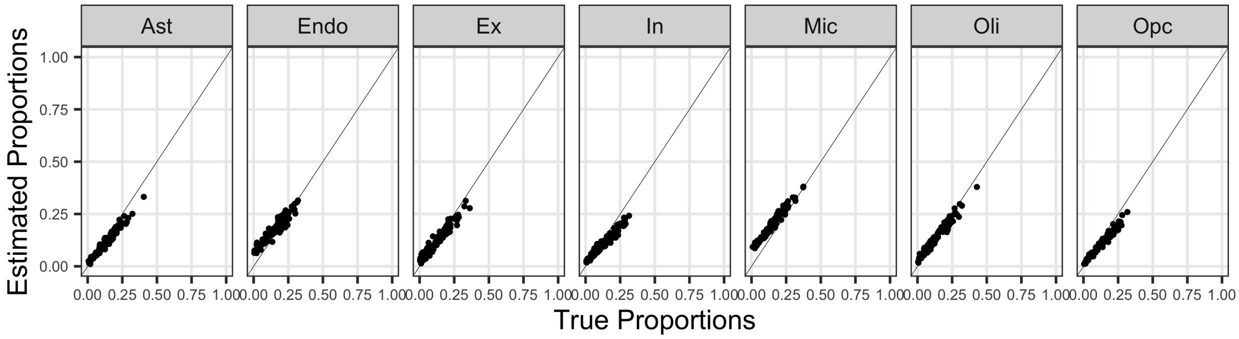


(c) CDSeq-estimated cell type proportions versus true proportions.

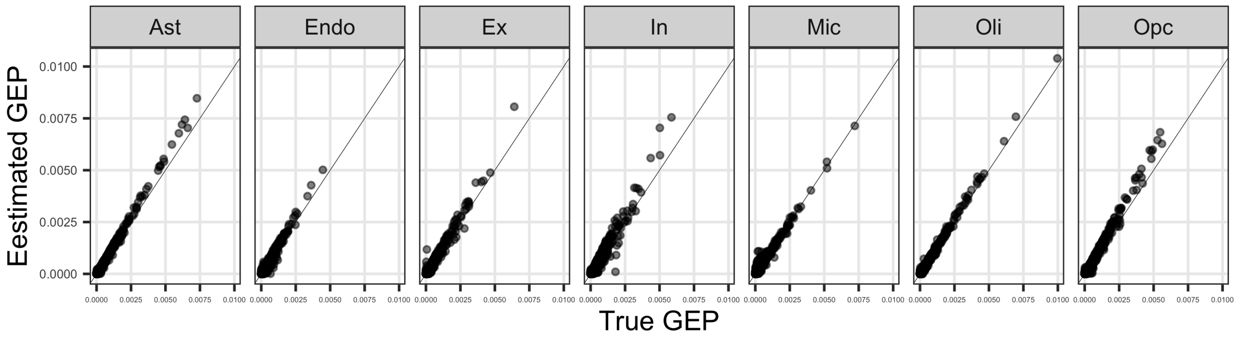


(d) CDSeq-estimated cell type GEPs versus true GEPs.

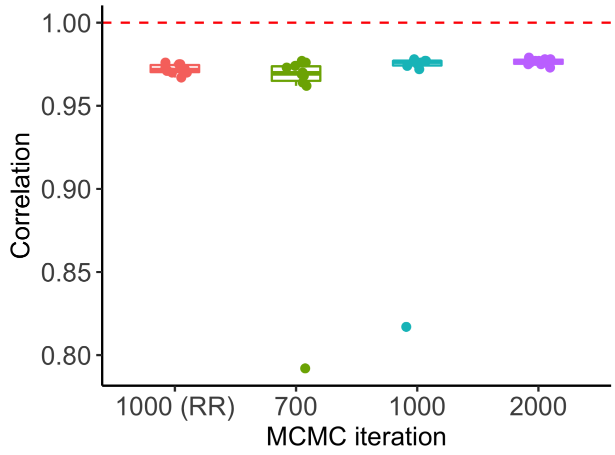


(e) Comparison of results using different MCMC runs.

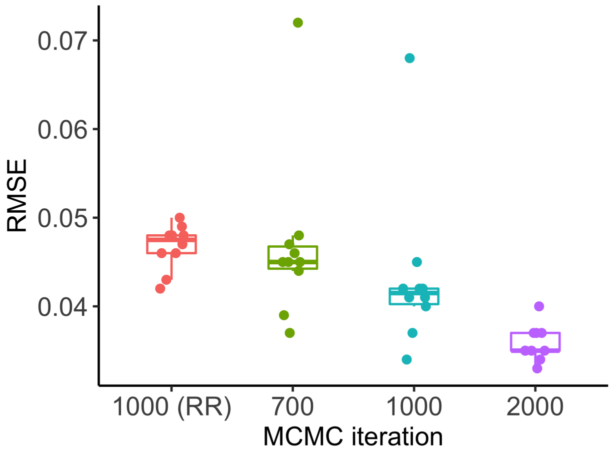


(f) Comparison of results using different MCMC runs.

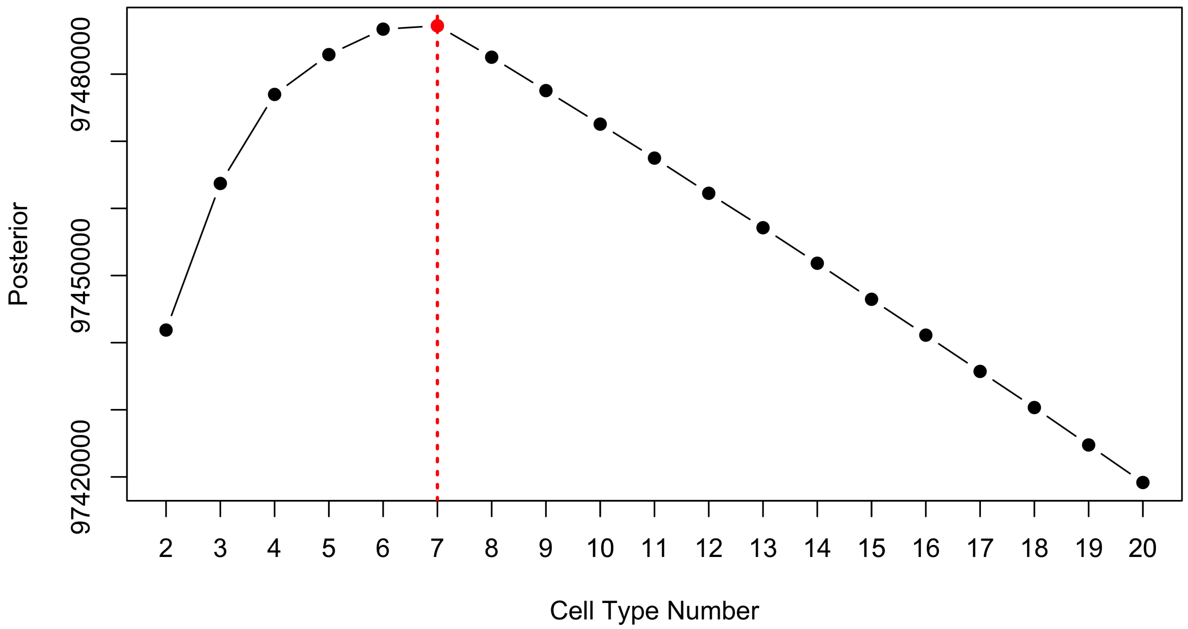


(g) CDSeq is able to accurately estimate the number of cell types in the mixtures.

Figure S7: Comparison of different deconvolution methods using synthetic mixtures generating by human brain frontal cortex single cell [15]. As shown in Fig. S7a and Fig. S7b, most methods performed well with correlations above 0.95. In particular, CDSeq provided accurate estimations for both cell type proportions (Fig.S7c) and cell-type-specific GEPs (Fig.S7d). We further tested the effect of different MCMC runs and show that more MCMC runs will provide more accurate results as shown in Fig.S7e and Fig.S7f. In addition, CDSeq is able to estimate the number of cell types in the bulk sample as indicated in Fig.S7g.

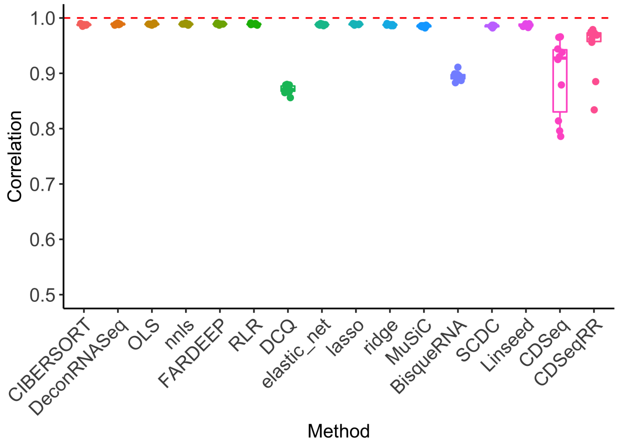


(a) Correlation comparisons of different deconvolution methods.

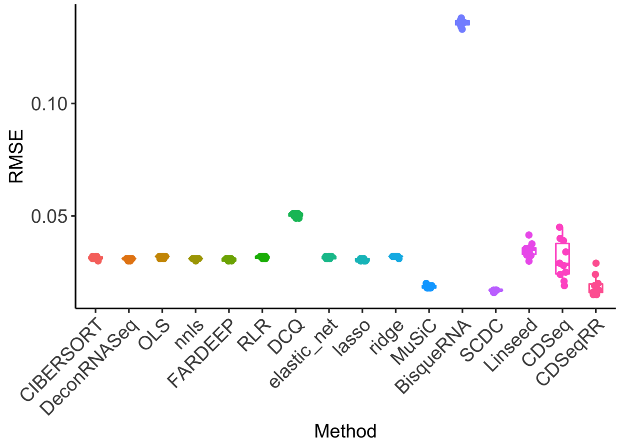


(b) RMSE comparisons of different deconvolution methods.

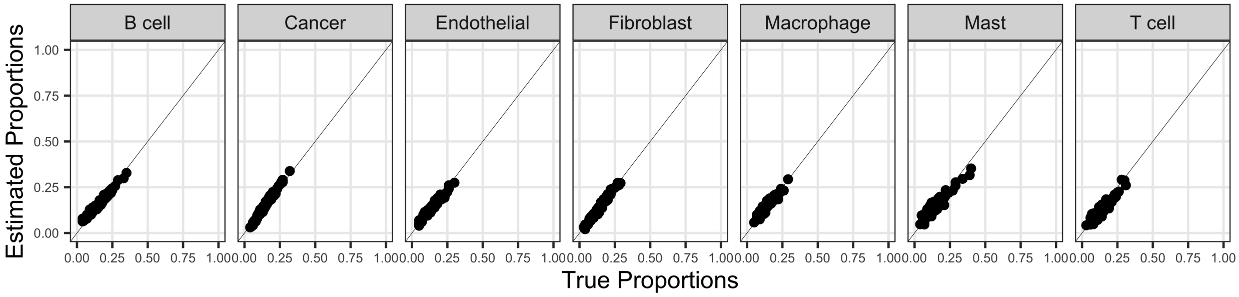


(c) CDSeq-estimated cell type proportions versus true proportions.

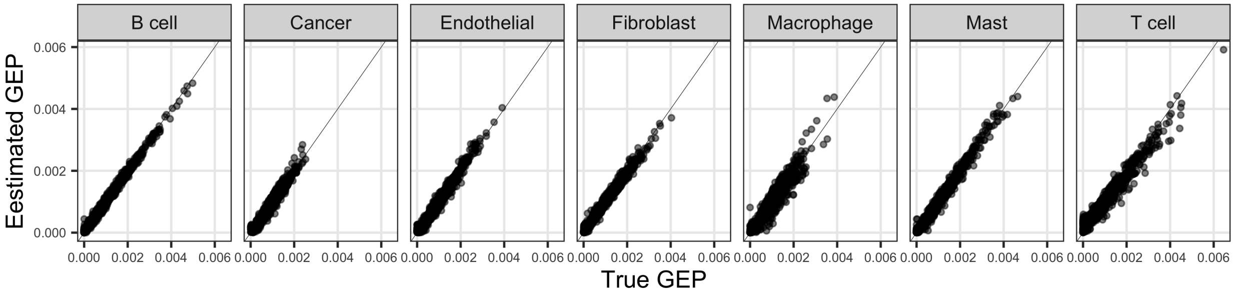


(d) CDSeq-estimated cell type GEPs versus true GEPs.

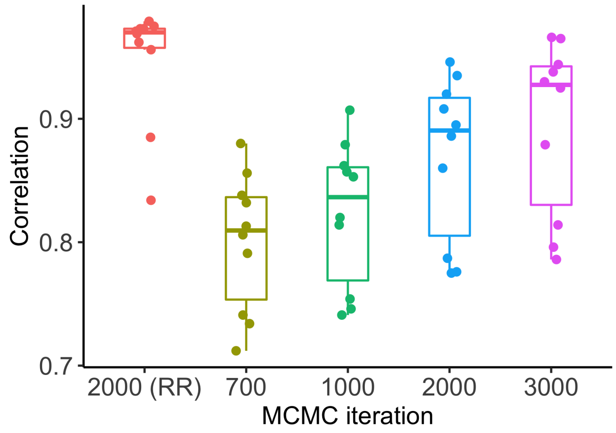


(e) Comparison of correlations using different MCMC runs.

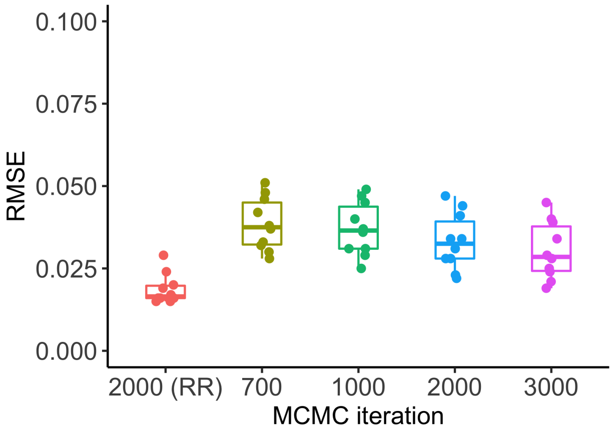


(f) Comparison of RMSEs using different MCMC runs.

Figure S8: Comparison of different deconvolution methods using synthetic mixtures generated by tumor single cell data [17]. It is noted that in this comparison CDSeq performed a bit worse than most deconvolution methods whose correlations are almost equal to 1. This is due to the fact that the scRNA-seq we downloaded are normalized data and the raw read counts data are not available. Therefore, we treated the normalized data as the raw counts by rounding off the normalized expression values to integers. Nevertheless, we showed that CDSeq-estimated cell type proportion and cell-type-specific GEPs are very accurate as shown in Fig.S8c and Fig.S8d.

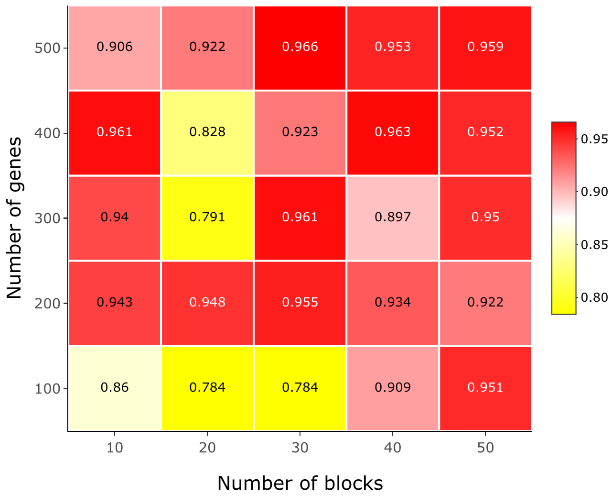


(a) Comparison of results using different number of genes and different number of blocks in Reduce-Recover setting.

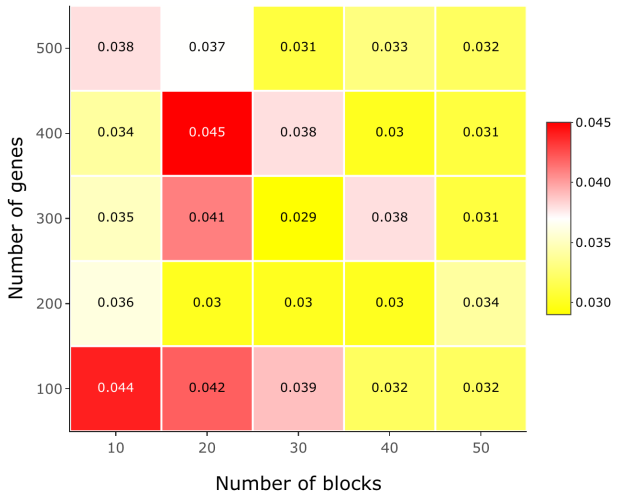


(b) Comparison of results using different number of genes and different number of blocks in Reduce-Recover setting.

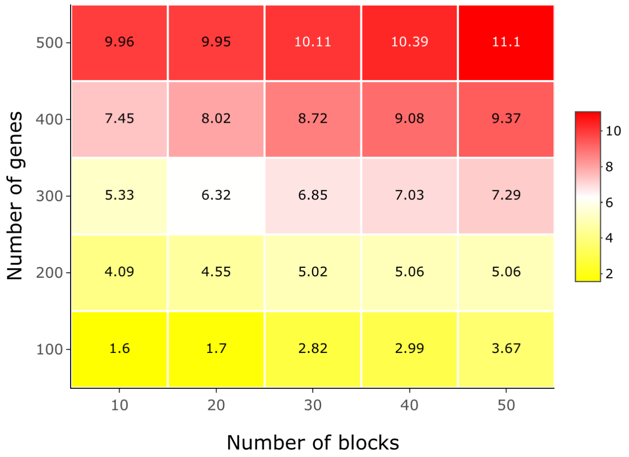


(c) Comparison of running times using different number of genes and different number of blocks in Reduce-Recover setting.

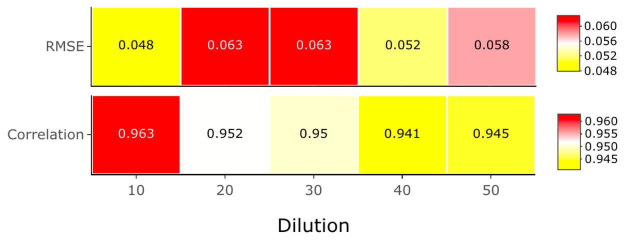


(d) Comparison of results using different parameter settings.

Figure S9: Comparison of CDSeq results using different parameter settings using synthetic PBMC mixtures [16]. We demonstrate the effect of using different number of genes and blocks in the Reduce-Recover mode in CDSeq. Overall, it is shown that more number of genes and blocks will result in better accuracy as shown in Fig.S9a and Fig.S9b. However, even using less genes and blocks, CDSeq is still able to provide reasonably good estimations. Using different dilution factors, we can see that the correlations are decreasing as the dilution factors increase.

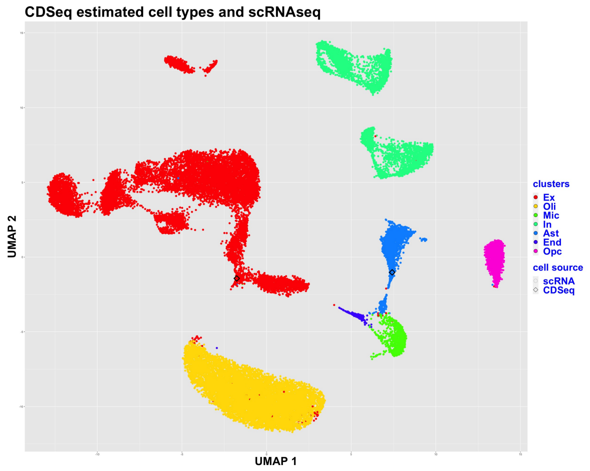


(a) Clustering of scRNA-seq (color dots) and CDSeq-estimated cell types (color squares). Generate 1 pseudo cell for each CDSeq-estimated cell type.

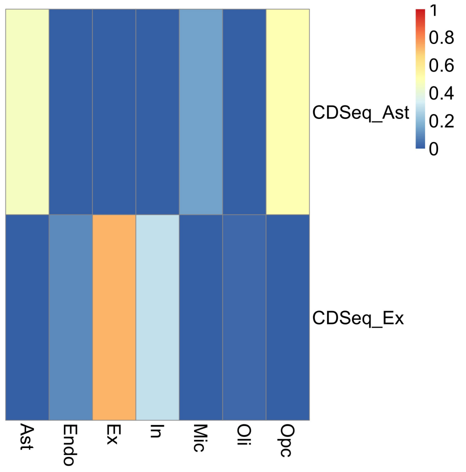


(b) Correlation between CDSeq-estimated cell type proportions and true proportions.

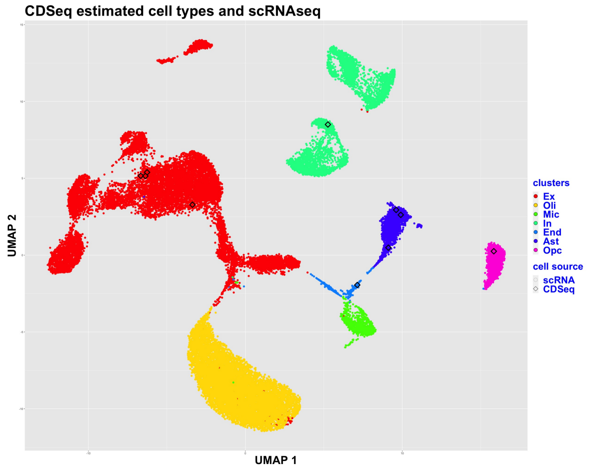


(c) Clustering of scRNA-seq (color dots) and CDSeq-estimated cell types (color squares). Generate 5 pseudo cell for each CDSeq-estimated cell type.

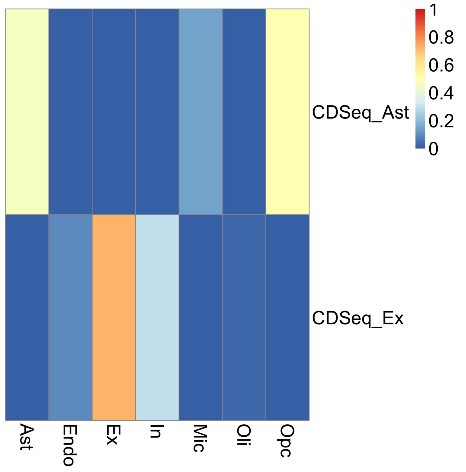


(d) Correlation between CDSeq-estimated cell type proportions and true proportions.

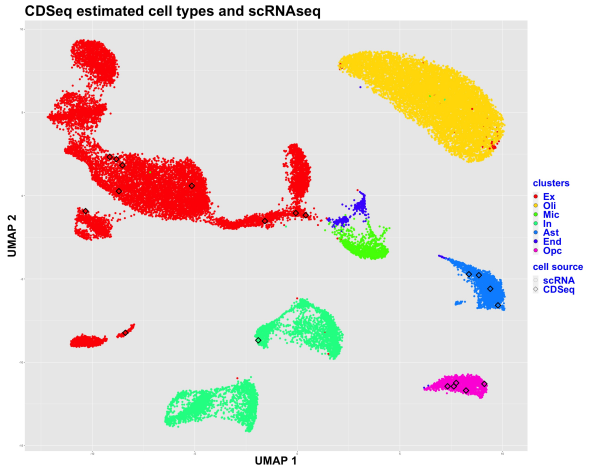


(e) Clustering of scRNA-seq (color dots) and CDSeq-estimated cell types (color squares). Generate 10 pseudo cell for each CDSeq-estimated cell type.

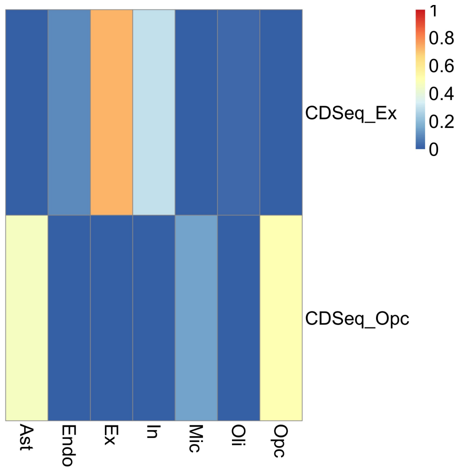


(f) Correlation between CDSeq-estimated cell type proportions and true proportions.

Figure S10: Annotating CDSeq-estimated cell types using scRNA-seq. Cell type number is 2. We show the clustering results of using different number of pseudo cells for each CDSeq-estimated cell types and the correlation between the corresponding cell type proportion and true proportion after annotation. Different number of pseudo cells produced the same annotation results.

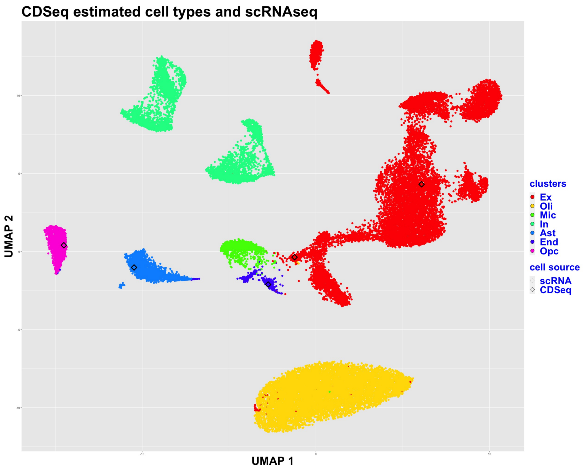


(a) Clustering of scRNA-seq (color dots) and CDSeq-estimated cell types (color squares). Pseudo cell number is 1.

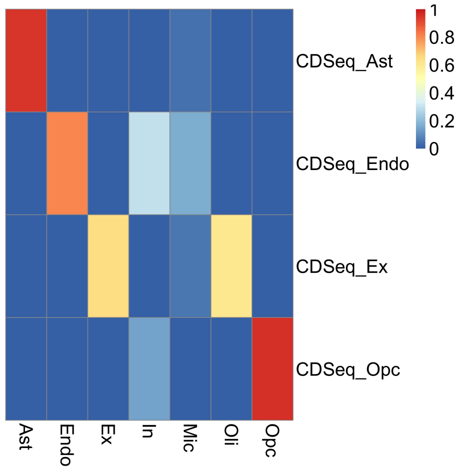


(b) Correlation between CDSeq-estimated cell type proportions and true proportions.

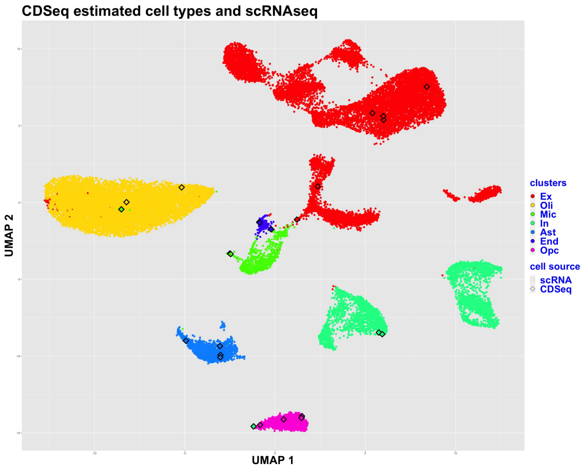


(c) Clustering of scRNA-seq (color dots) and CDSeq-estimated cell types (color squares). Pseudo cell number is 5.

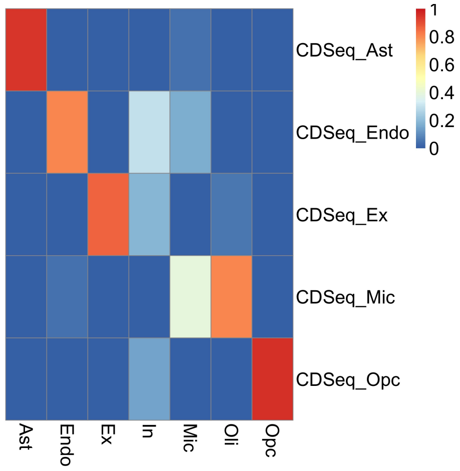


(d) Correlation between CDSeq-estimated cell type proportions and true proportions.

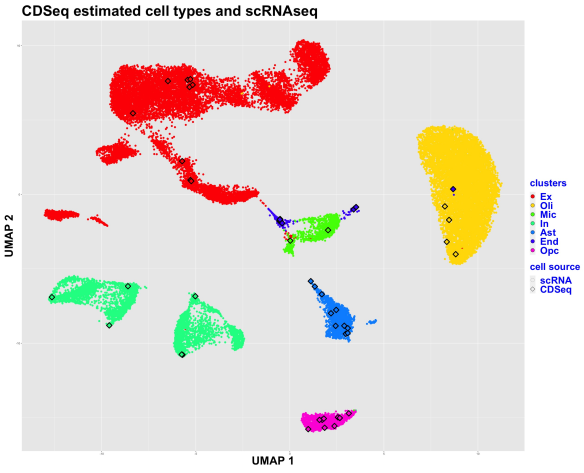


(e) Clustering of scRNA-seq (color dots) and CDSeq-estimated cell types (color squares). Pseudo cell number is 10.

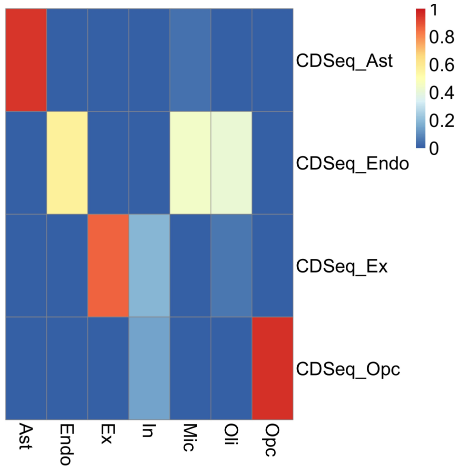


(f) Correlation between CDSeq-estimated cell type proportions and true proportions.

Figure S11: Annotating CDSeq-estimated cell types using scRNA-seq. Cell type number is 5. We show the clustering results of using different number of pseudo cells for each CDSeq-estimated cell types and the correlation between the corresponding cell type proportion and true proportion after annotation. Different number of pseudo cells produced the same annotation results.

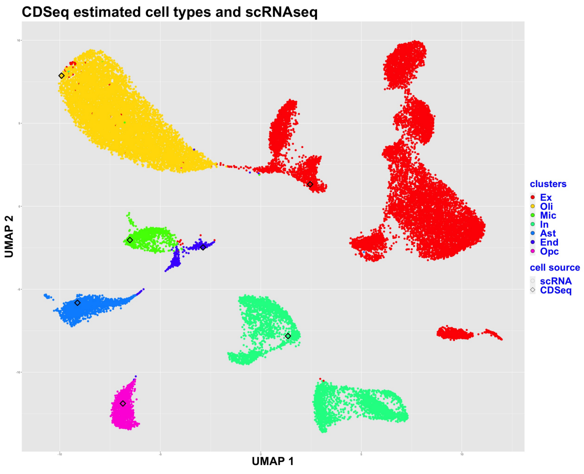


(a) Clustering of scRNA-seq (color dots) and CDSeq-estimated cell types (color squares). Pseudo cell number is 1.

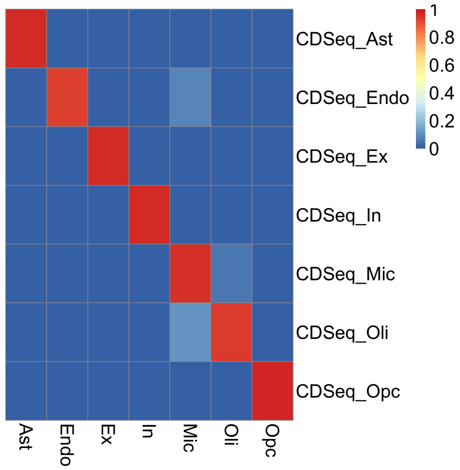


(b) Correlation between CDSeq-estimated cell type proportions and true proportions.

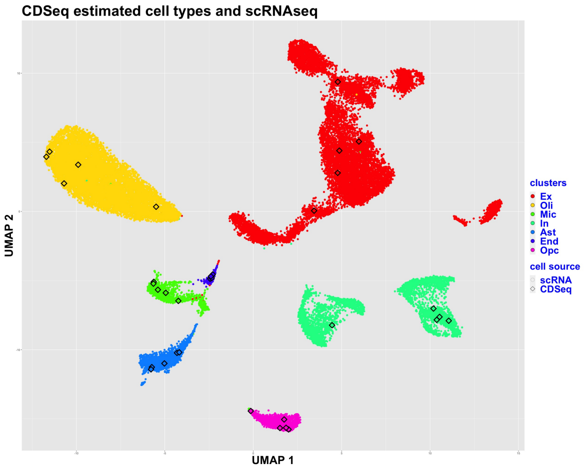


(c) Clustering of scRNA-seq (color dots) and CDSeq-estimated cell types (color squares). Pseudo cell number is 5.

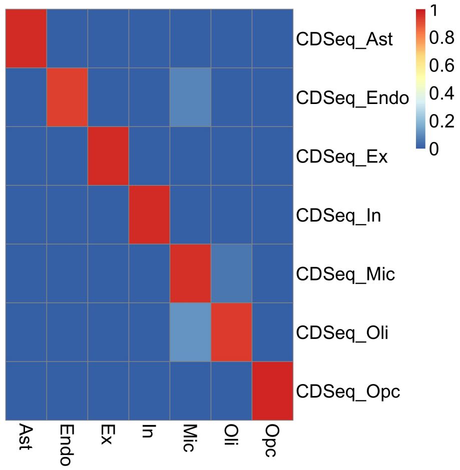


(d) Correlation between CDSeq-estimated cell type proportions and true proportions.

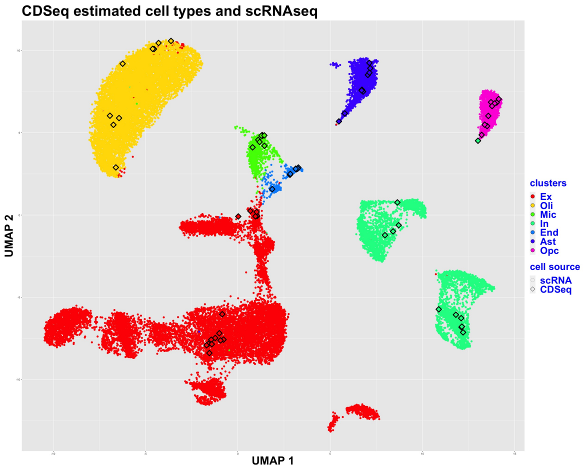


(e) Clustering of scRNA-seq (color dots) and CDSeq-estimated cell types (color squares). Pseudo cell number is 10.

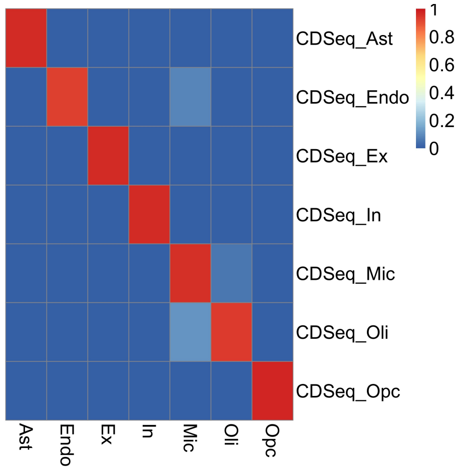


(f) Correlation between CDSeq-estimated cell type proportions and true proportions.

Figure S12: Annotating CDSeq-estimated cell types using scRNA-seq. Cell type number is 7. We show the clustering results of using different number of pseudo cells for each CDSeq-estimated cell types and the correlation between the corresponding cell type proportion and true proportion after annotation. Different number of pseudo cells produced the same annotation results.

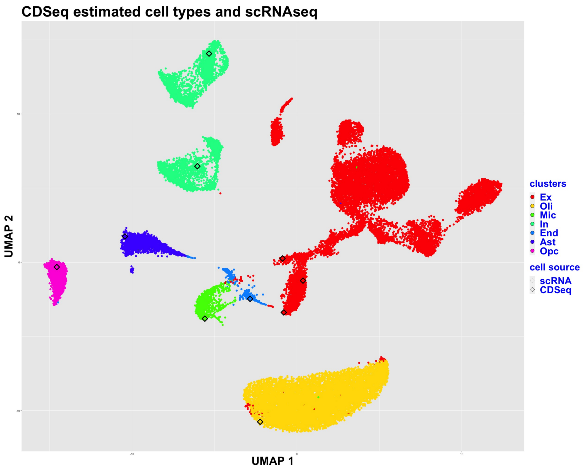


(a) Clustering of scRNA-seq (color dots) and CDSeq-estimated cell types (color squares). Pseudo cell number is 1.

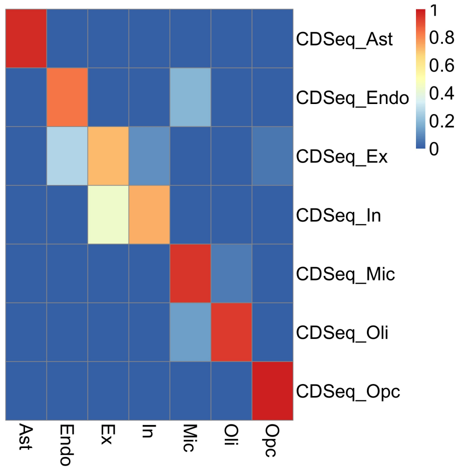


(b) Correlation between CDSeq-estimated cell type proportions and true proportions.

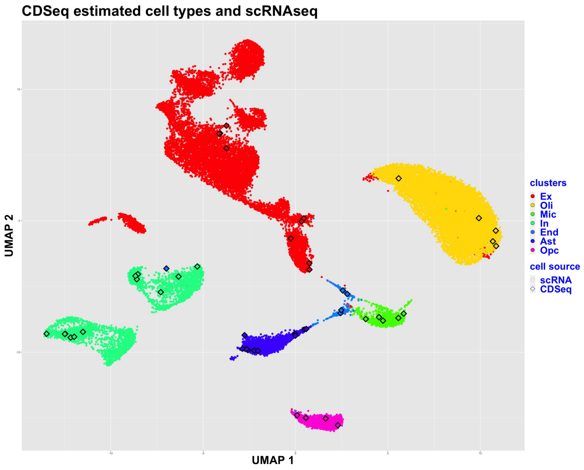


(c) Clustering of scRNA-seq (color dots) and CDSeq-estimated cell types (color squares). Pseudo cell number is 5.

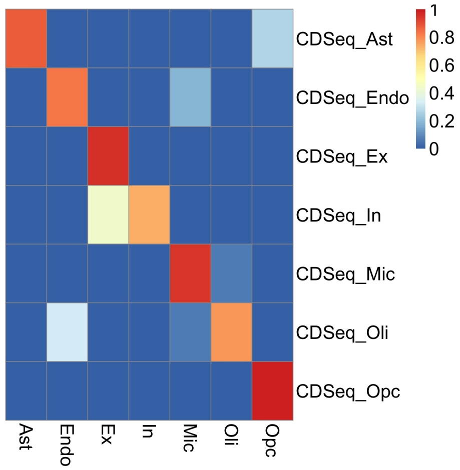


(d) Correlation between CDSeq-estimated cell type proportions and true proportions.

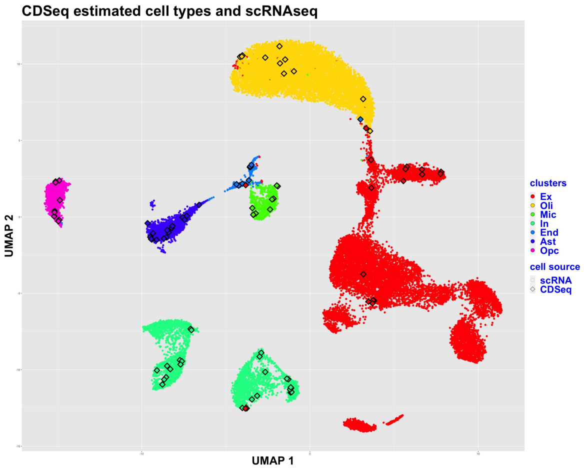


(e) Clustering of scRNA-seq (color dots) and CDSeq-estimated cell types (color squares). Pseudo cell number is 10.

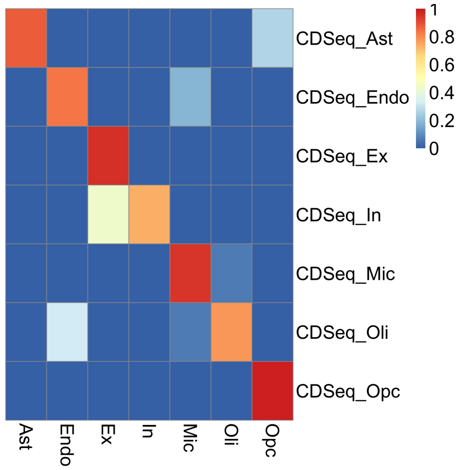


(f) Correlation between CDSeq-estimated cell type proportions and true proportions.

Figure S13: Annotating CDSeq-estimated cell types using scRNA-seq. Cell type number is 10. We show the clustering results of using different number of pseudo cells for each CDSeq-estimated cell types and the correlation between the corresponding cell type proportion and true proportion after annotation. Different number of pseudo cells produced the same annotation results.

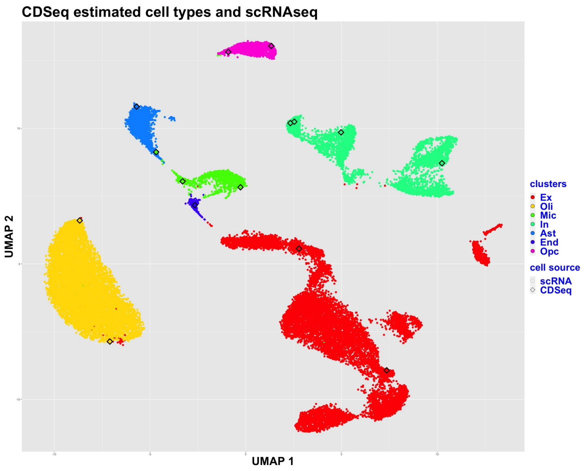


(a) Clustering of scRNA-seq (color dots) and CDSeq-estimated cell types (color squares). Pseudo cell number is 1.

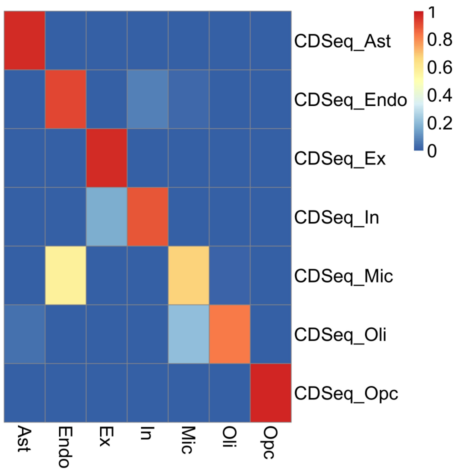


(b) Correlation between CDSeq-estimated cell type proportions and true proportions.

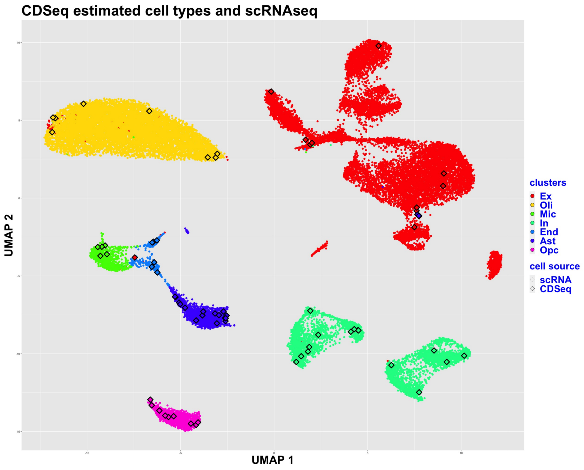


(c) Clustering of scRNA-seq (color dots) and CDSeq-estimated cell types (color squares). Pseudo cell number is 5.

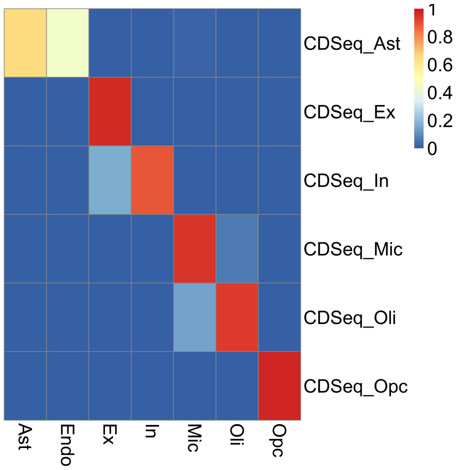


(d) Correlation between CDSeq-estimated cell type proportions and true proportions.

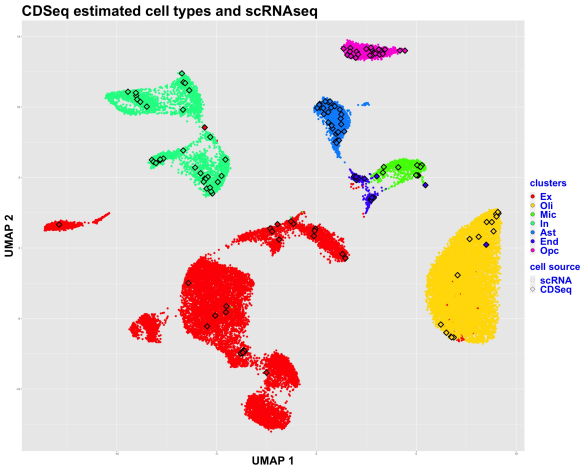


(e) Clustering of scRNA-seq (color dots) and CDSeq-estimated cell types (color squares). Pseudo cell number is 10.

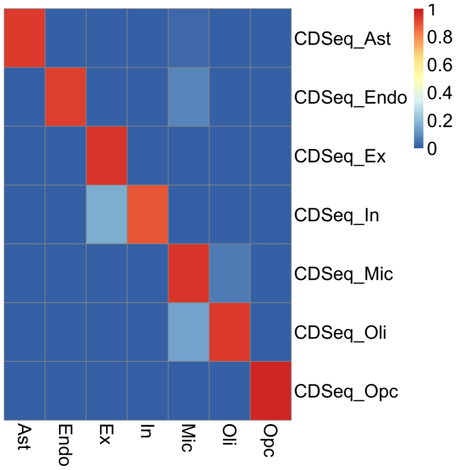


(f) Correlation between CDSeq-estimated cell type proportions and true proportions.

Figure S14: Annotating CDSeq-estimated cell types using scRNA-seq. Cell type number is 15. We show the clustering results of using different number of pseudo cells for each CDSeq-estimated cell types and the correlation between the corresponding cell type proportion and true proportion after annotation. Different number of pseudo cells produced the same annotation results.

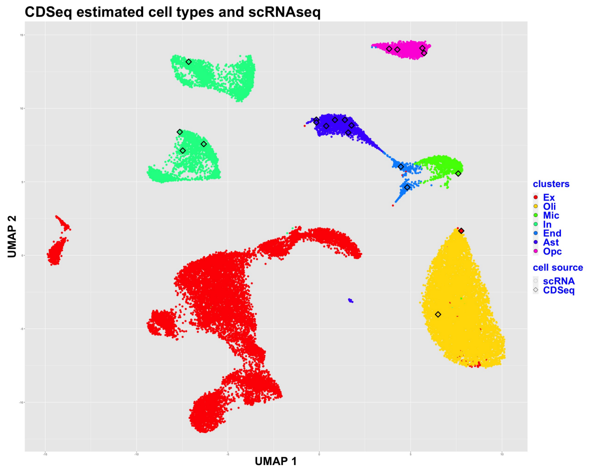


(a) Clustering of scRNA-seq (color dots) and CDSeq-estimated cell types (color squares). Pseudo cell number is 1.

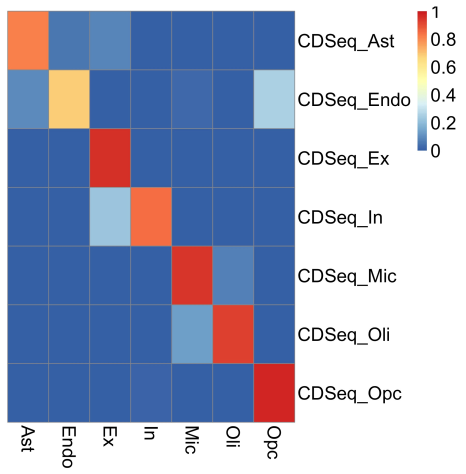


(b) Correlation between CDSeq-estimated cell type proportions and true proportions.

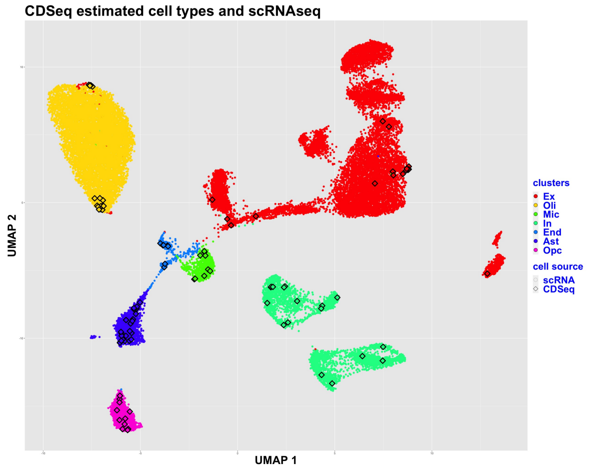


(c) Clustering of scRNA-seq (color dots) and CDSeq-estimated cell types (color squares). Pseudo cell number is 5.

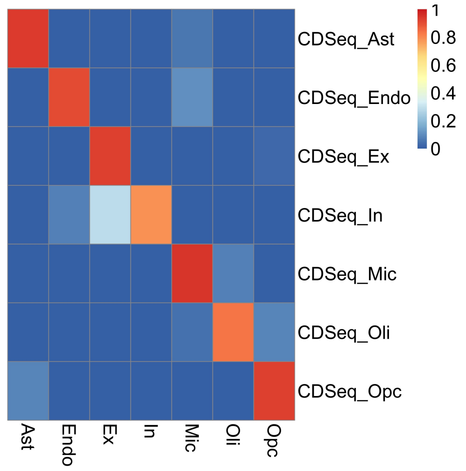


(d) Correlation between CDSeq-estimated cell type proportions and true proportions.

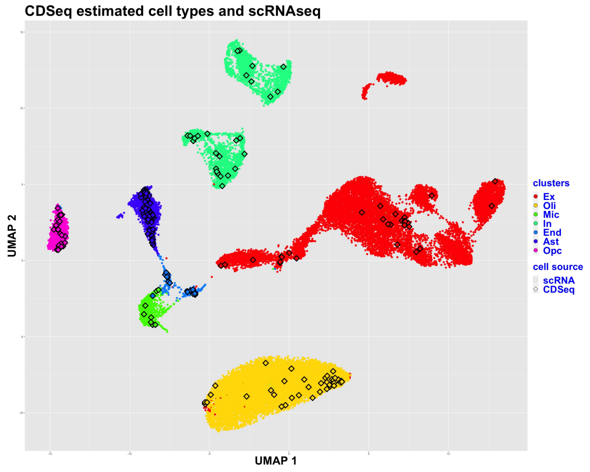


(e) Clustering of scRNA-seq (color dots) and CDSeq-estimated cell types (color squares). Pseudo cell number is 10.

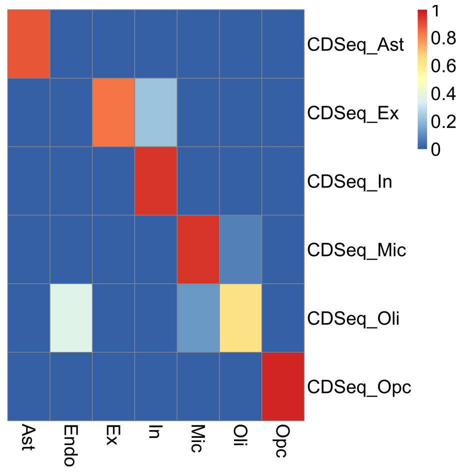


(f) Correlation between CDSeq-estimated cell type proportions and true proportions.

Figure S15: Annotating CDSeq-estimated cell types using scRNA-seq. Cell type number is 20. We show the clustering results of using different number of pseudo cells for each CDSeq-estimated cell types and the correlation between the corresponding cell type proportion and true proportion after annotation. Different number of pseudo cells produced the same annotation results.

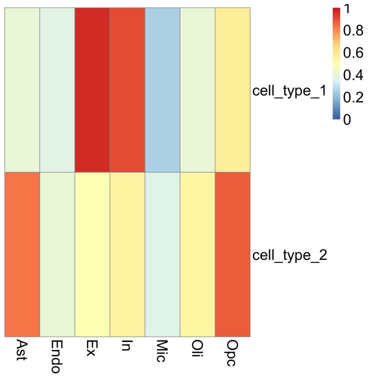


(a) Correlation bwteen CDSeq-estimates and scRNA-seq pseudo-bulk.

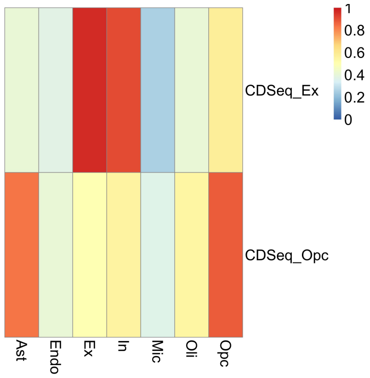


(b) Correlation bwteen annotated CDSeq-estimates and scRNA-seq pseudo-bulk.

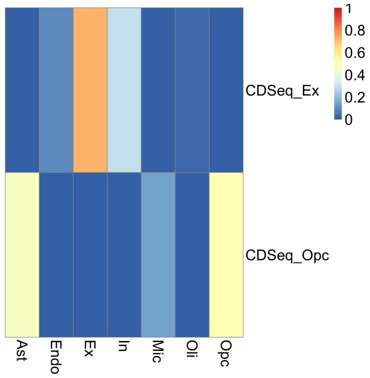


(c) Correlation between CDSeq-estimated cell type proportion with true proportion.

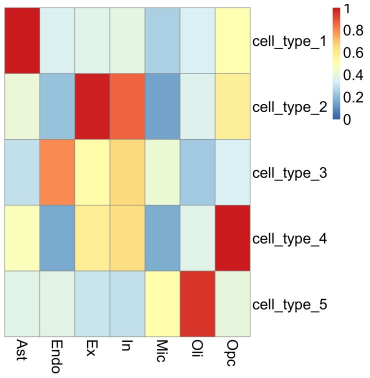


(d) Correlation bwteen CDSeq-estimates and scRNA-seq pseudo-bulk.

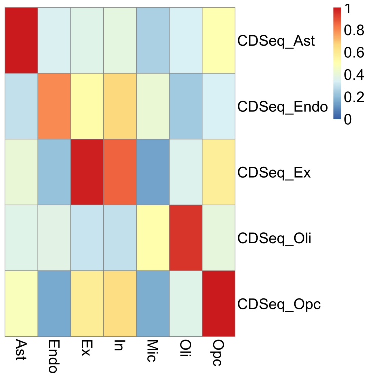


(e) Correlation bwteen annotated CDSeq-estimates and scRNA-seq pseudo-bulk.

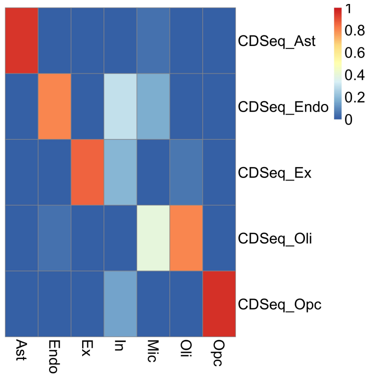


(f) Correlation between CDSeq-estimated cell type proportion with true proportion.

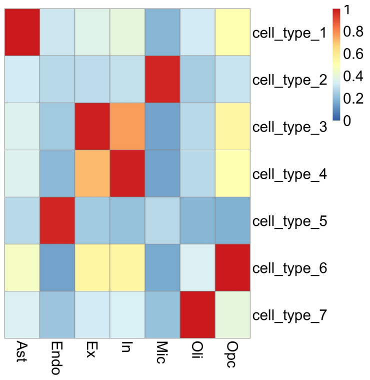


(g) Correlation bwteen CDSeq-estimates and scRNA-seq pseudo-bulk.

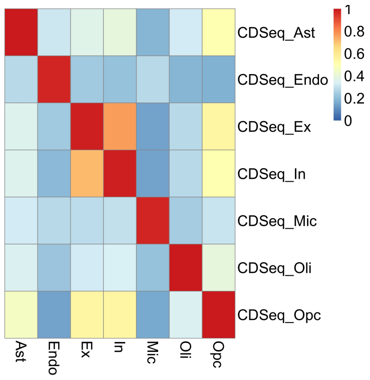


(h) Correlation bwteen annotated CDSeq-estimates and scRNA-seq pseudo-bulk.

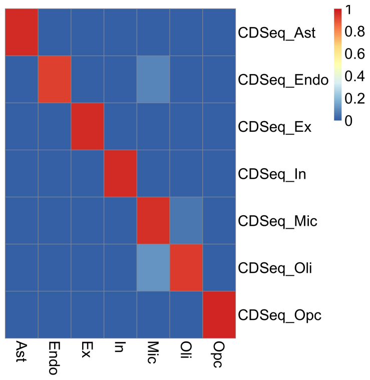


(i) Correlation between CDSeq-estimated cell type proportion with true proportion.

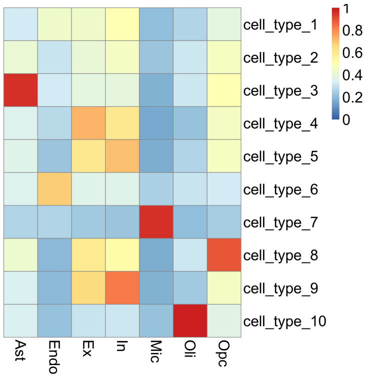


(j) Correlation bwteen CDSeq-estimates and scRNA-seq pseudo-bulk.

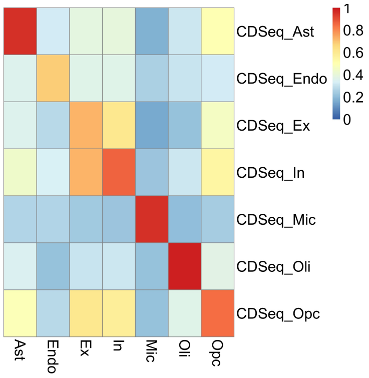


(k) Correlation bwteen annotated CDSeq-estimates and scRNA-seq pseudo-bulk.

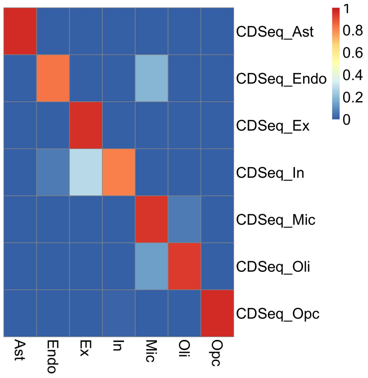


(l) Correlation between CDSeq-estimated cell type proportion with true proportion.

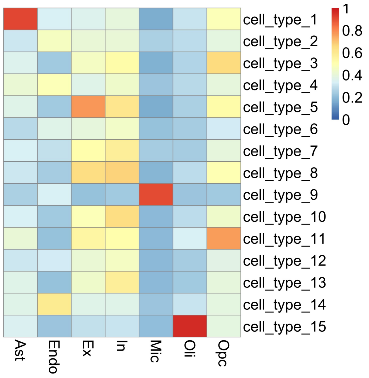


(m) Correlation bwteen CDSeq-estimates and scRNA-seq pseudo-bulk.

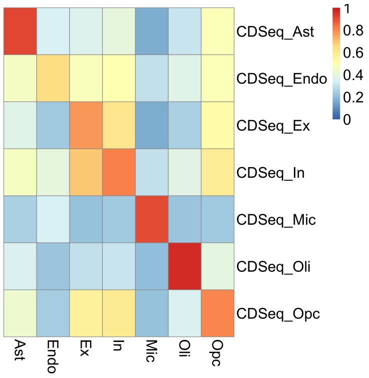


(n) Correlation bwteen annotated CDSeq-estimates and scRNA-seq pseudo-bulk.

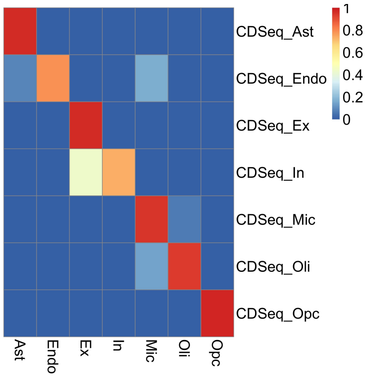


(o) Correlation between CDSeq-estimated cell type proportion with true proportion.

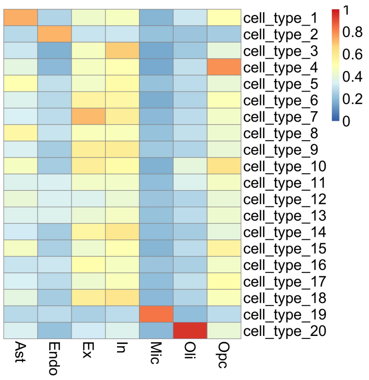


(p) Correlation bwteen CDSeq-estimates and scRNA-seq pseudo-bulk.

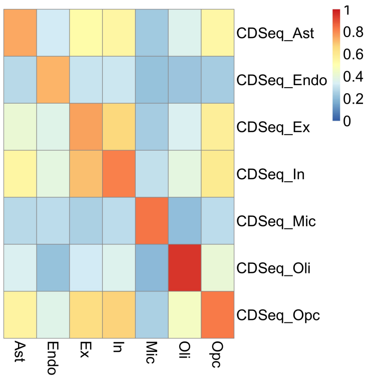


(q) Correlation bwteen annotated CDSeq-estimates and scRNA-seq pseudo-bulk.

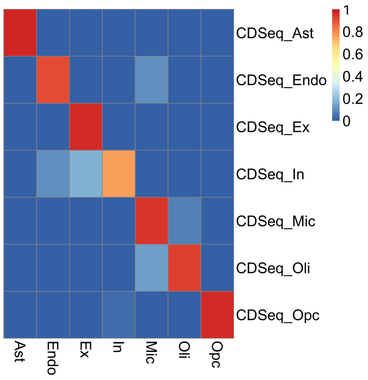


(r) Correlation between CDSeq-estimated cell type proportion with true proportion.

Figure S16: Annotating CDSeq-estimated cell types using scRNA-seq reference and correlation analysis.

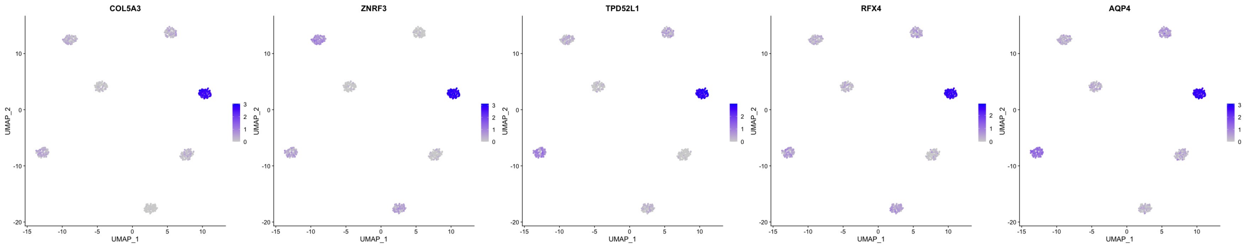


(a) Expression level of the top 5 marker genes of CDSeq-estimated cell type 1.

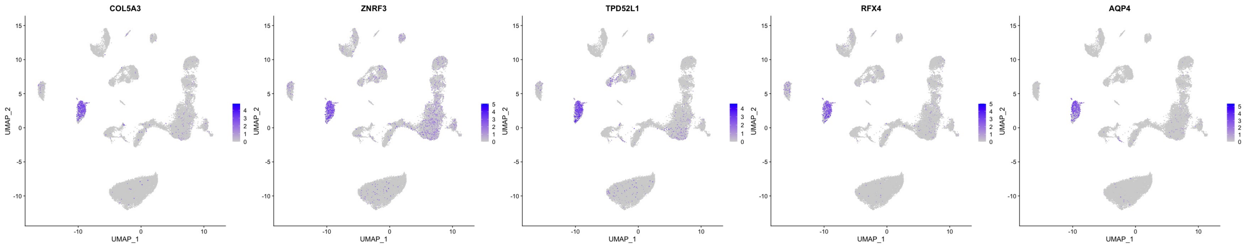


(b) Expression level of top markers in CDSeq-estimated cell type 1 in scRNA-seq.

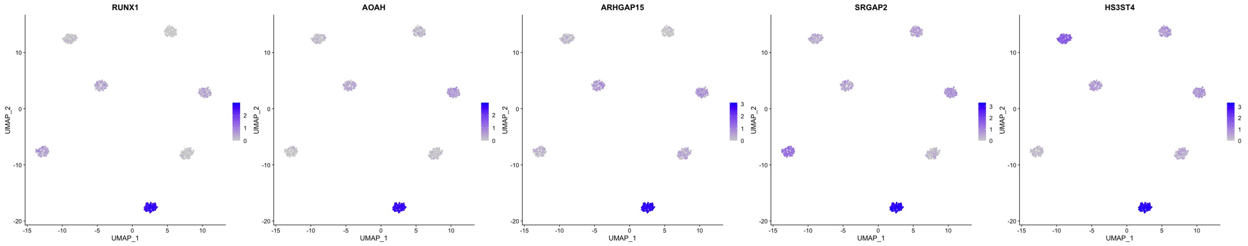


(c) Expression level of the top 5 marker genes of CDSeq-estimated cell type 2.

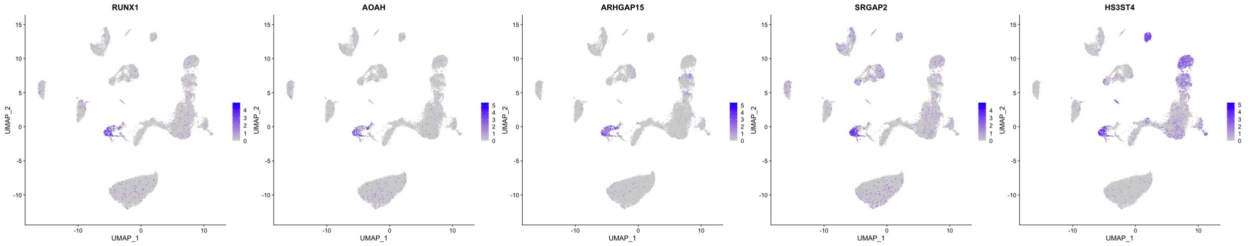


(d) Expression level of top markers in CDSeq-estimated cell type 2 in scRNA-seq.

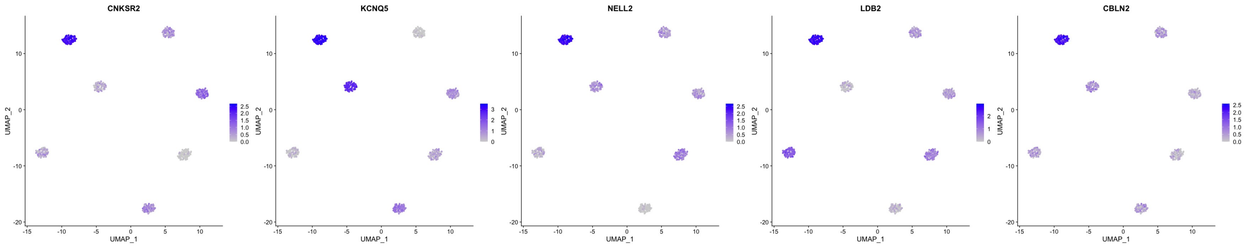


(e) Expression level of the top 5 marker genes of CDSeq-estimated cell type 3.

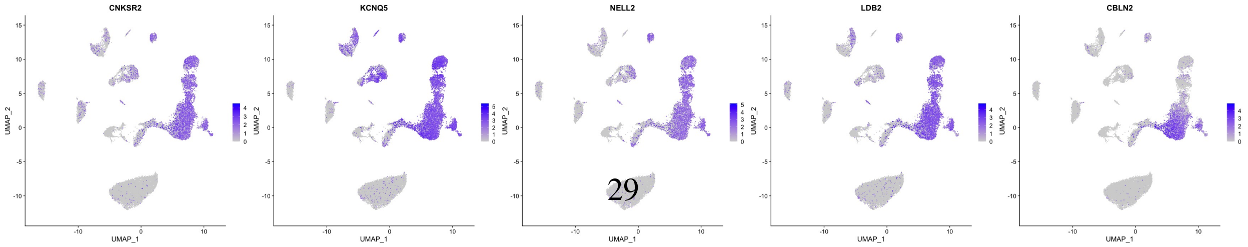


(f) Expression level of top markers in CDSeq-estimated cell type 3 in scRNA-seq.

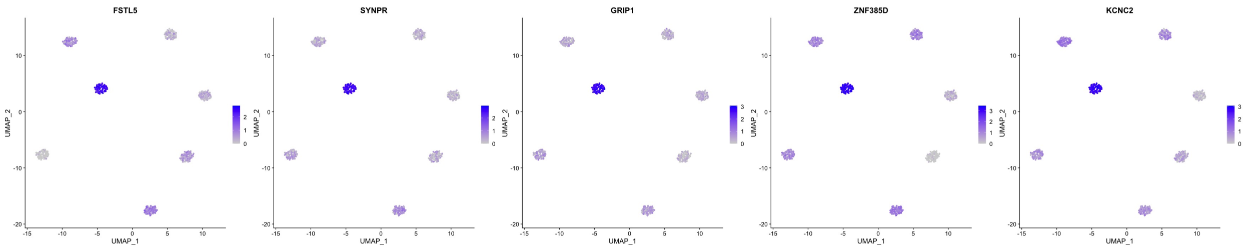


(g) Expression level of the top 5 marker genes of CDSeq-estimated cell type 4.

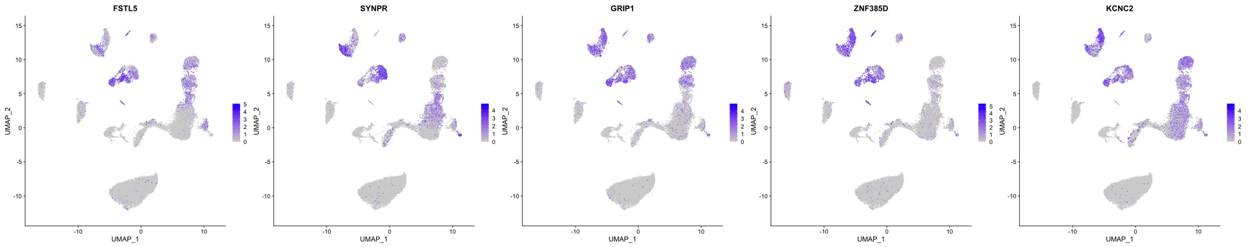


(h) Expression level of top markers in CDSeq-estimated cell type 4 in scRNA-seq.

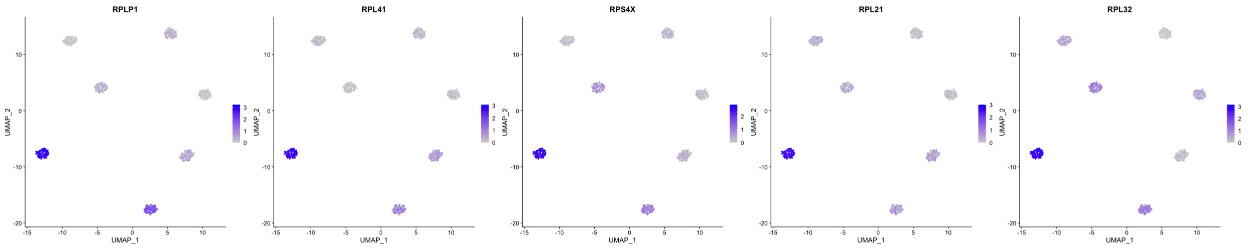


(i) Expression level of the top 5 marker genes of CDSeq-estimated cell type 5.

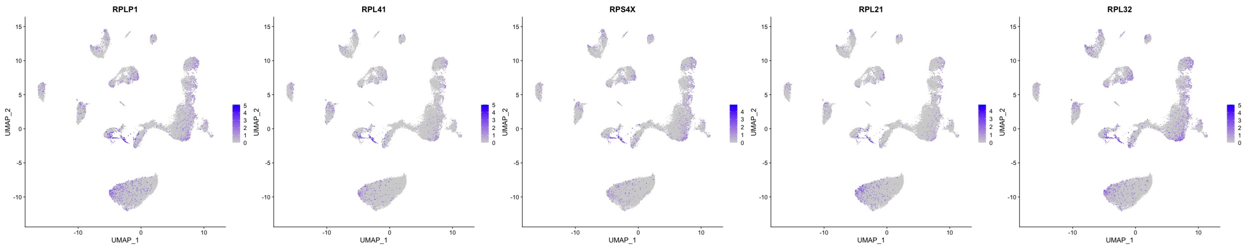


(j) Expression level of top markers in CDSeq-estimated cell type 5 in scRNA-seq.

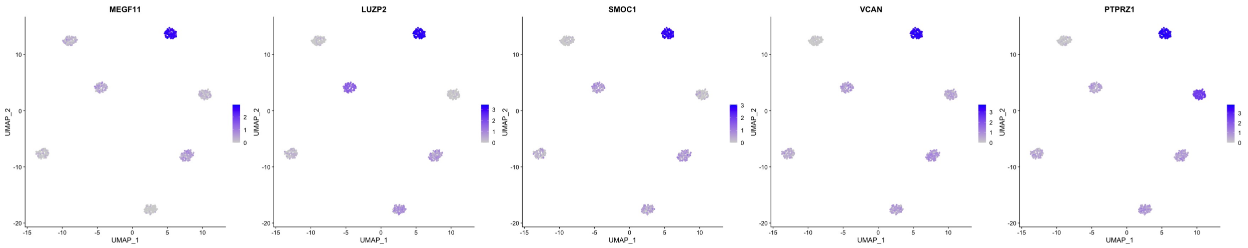


(k) Expression level of the top 5 marker genes of CDSeq-estimated cell type 6.

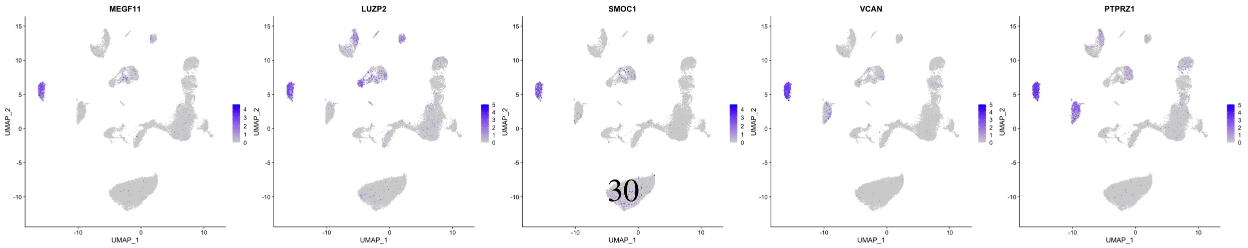


(l) Expression level of top markers in CDSeq-estimated cell type 6 in scRNA-seq.

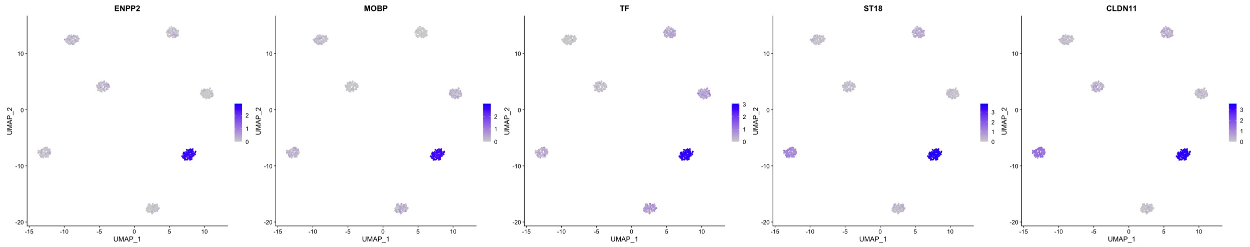


(m) Expression level of the top 5 marker genes of CDSeq-estimated cell type 7.

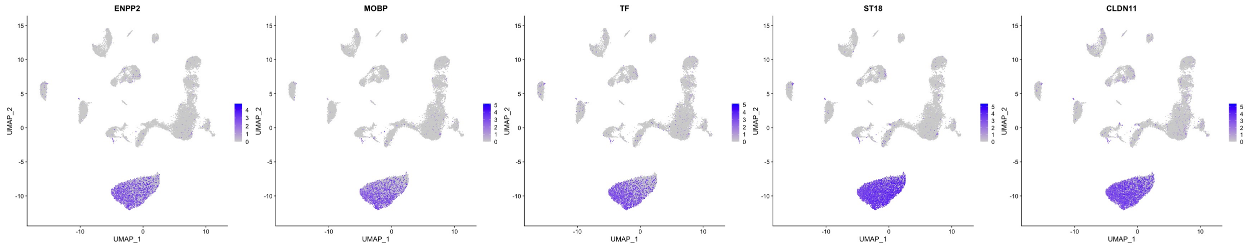


(n) Expression level of top markers in CDSeq-estimated cell type 7 in scRNA-seq.

Figure S17: Annotating CDSeq-estimated cell types using scRNA-seq without reference. We show that the all the top 5 marker genes of each cluster of CDSeq-estimated cell types are also highly expressed in very specific clusters in the scRNA-seq data suggesting the potential of annotating CDSe-estimated cell types without any reference data.

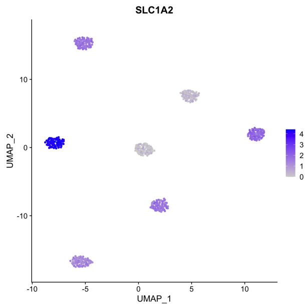


(a) Expression level of Astrocyte marker gene SLC1A2.

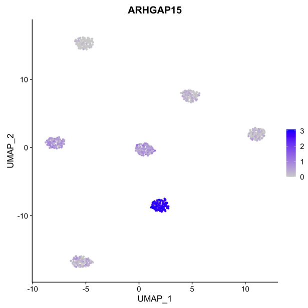


(b) Expression level of Microglia marker gene ARHGAP15.

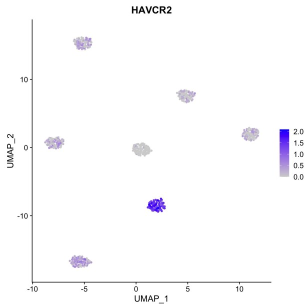


(c) Expression level of Microglia marker gene HAVCR2.

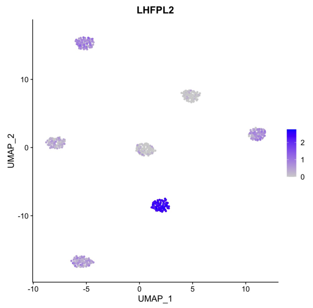


(d) Expression level of Microglia marker gene LHFPL2.

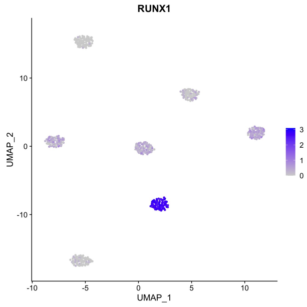


(e) Expression level of Microglia marker gene RUNX1.

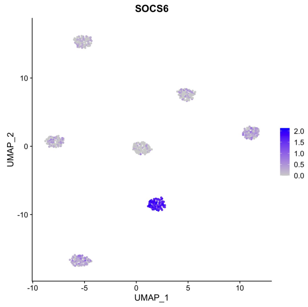


(f) Expression level of Microglia marker gene SOCS6.

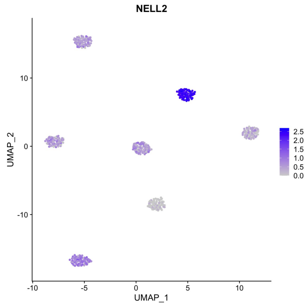


(g) Expression level of Neuron marker gene NELL2.

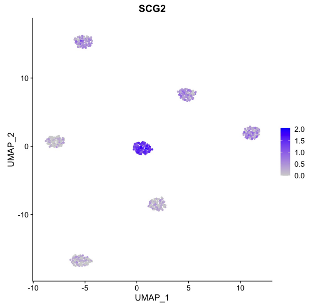


(h) Expression level of Neuron marker gene SCG2.

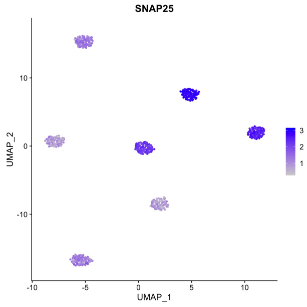


(i) Expression level of Neuron marker gene SNAP25.

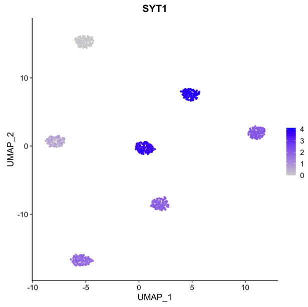


(j) Expression level of Neuron marker gene SYT1.

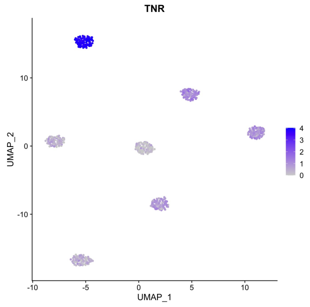


(k) Expression level of oligodendrocyte precursor cell (OPC) marker gene TNR.

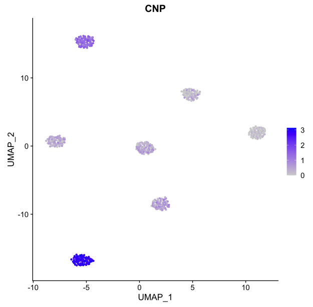


(l) Expression level of Oligodendrocyte marker gene CNP.

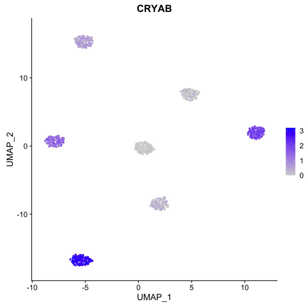


(m) Expression level of Oligodendrocyte marker gene CRYAB.

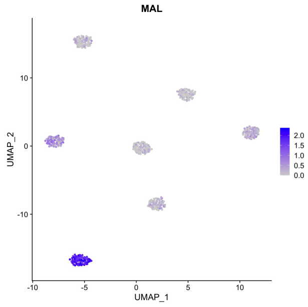


(n) Expression level of Oligodendrocyte marker gene MAL.

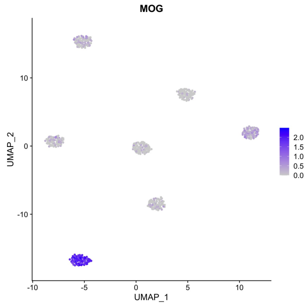


(o) Expression level of Oligodendrocyte marker gene MOG.

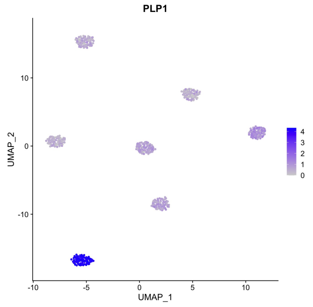


(p) Expression level of Oligodendrocyte marker gene PLP1.

Figure S18: Annotating CDSeq-estimated cell types using scRNA-seq without reference. We show that cell-type-specific marker genes [19, 20] are highly expressed in specific CDSeq-estimated cell type clusters suggesting CDSeq identifies cell-type-specific GEPs and it is possible to use marker genes to annotate CDSeq-estimated cell types. We further verify that, in Supplementary Figure S20, these marker genes are highly expressed in specific clusters.

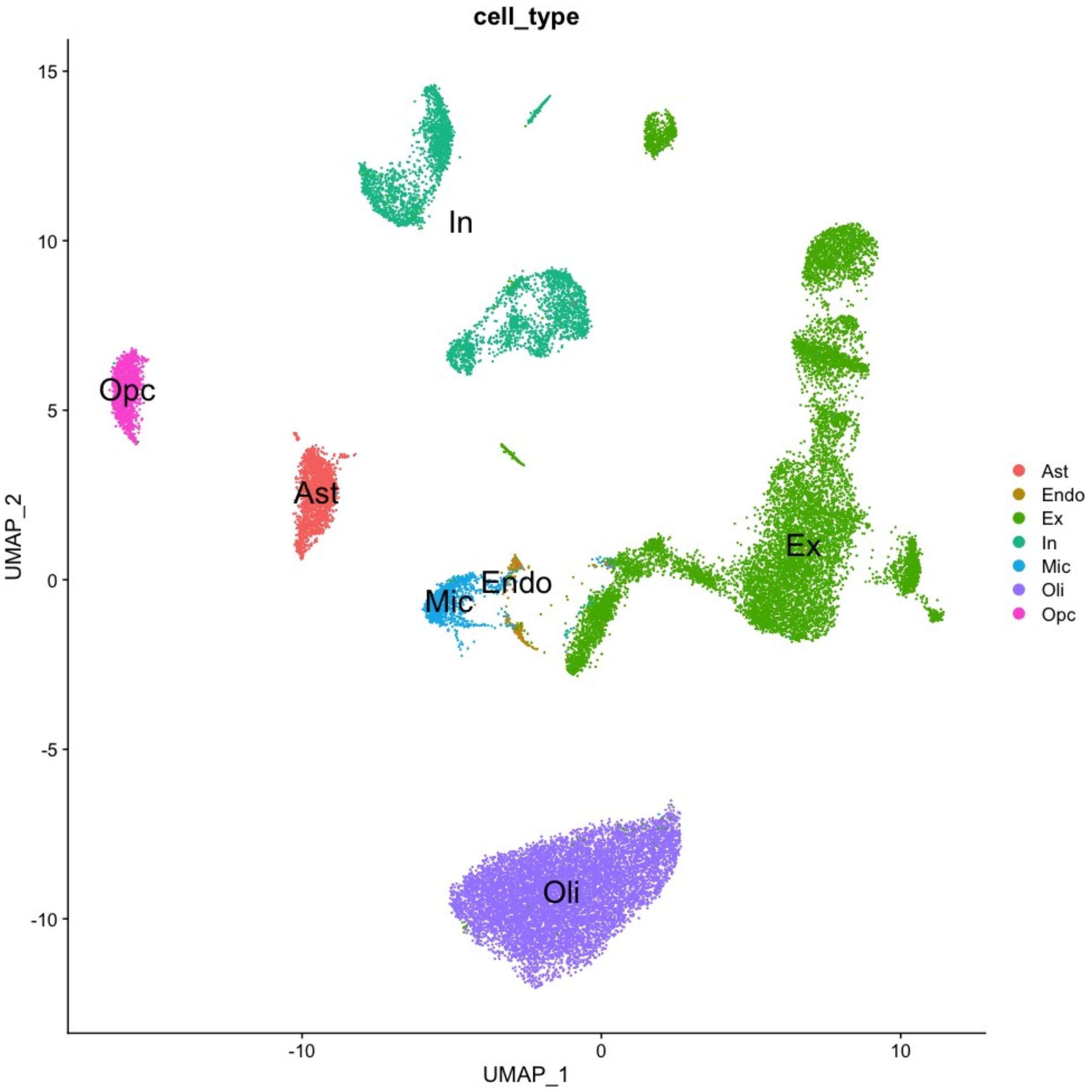


Figure S19: Brain frontal cortex single cell data with annotations.

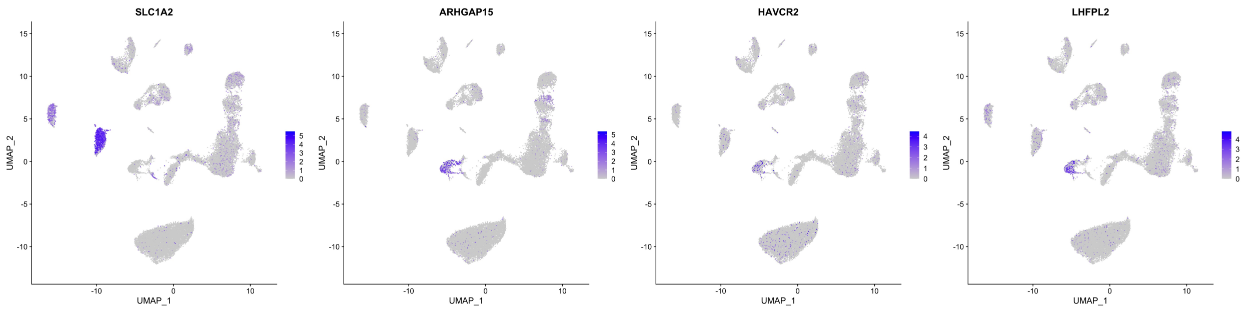


(a) Expression level of Astrocyte marker gene SLC1A2.

(b) Expression level of Microglia marker gene ARHGAP15.

(c) Expression level of Microglia marker gene HAVCR2.

(d) Expression level of Microglia marker gene LHFPL2.

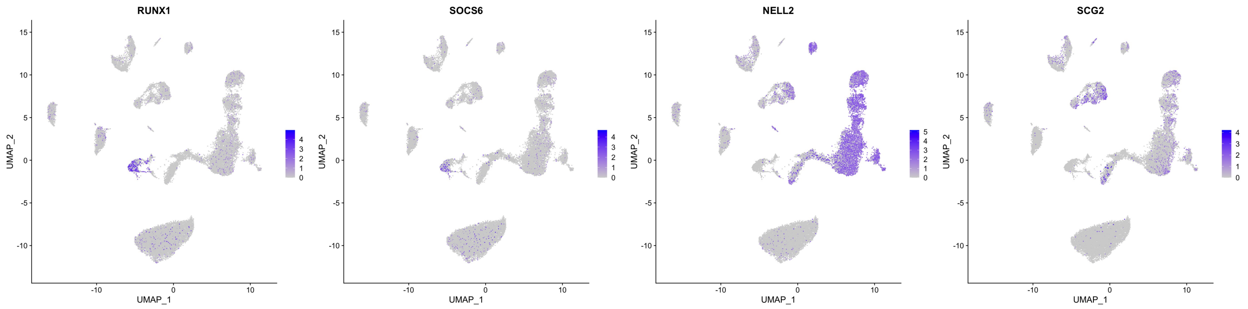


(e) Expression level of Microglia marker gene RUNX1.

(f) Expression level of Microglia marker gene SOCS6.

(g) Expression level of Neuron marker gene NELL2.

(h) Expression level of Neuron marker gene SCG2.

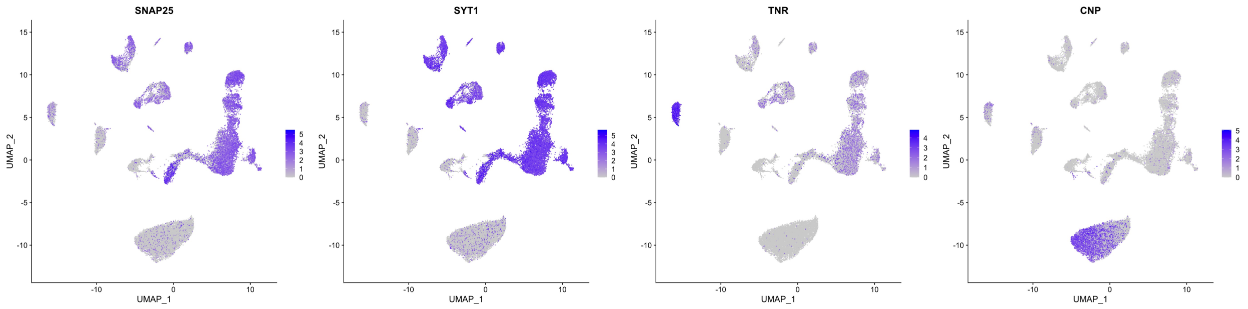


(i) Expression level of Neuron marker gene SNAP25.

(j) Expression level of Neuron marker gene SYT1.

(k) Expression level of oligodendrocyte precursor cell (OPC) marker gene TNR.

(l) Expression level of Oligodendrocyte marker gene CNP.

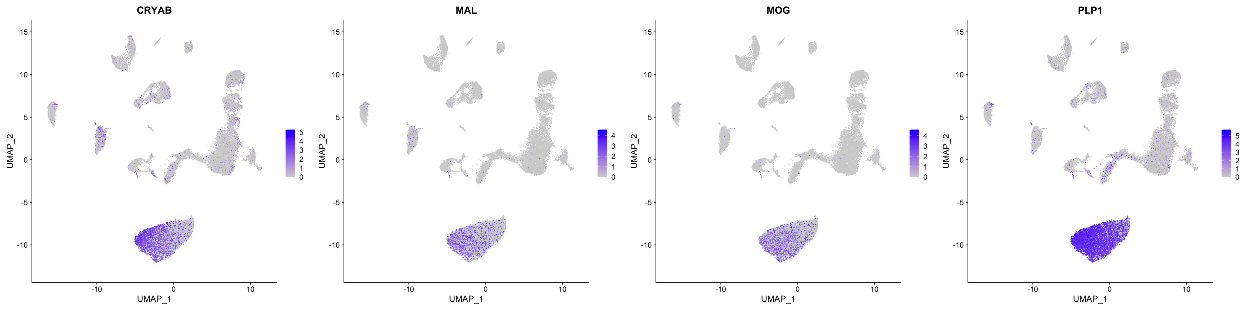


(m) Expression level of Oligodendrocyte marker gene CRYAB.

(n) Expression level of Oligodendrocyte marker gene MAL.

(o) Expression level of Oligodendrocyte marker gene MOG.

(p) Expression level of Oligodendrocyte marker gene PLP1.

Figure S20: Annotating CDSeq-estimated cell types using scRNA-seq without reference. We show that the cell-type-specific markers are highly expressed in specific clusters in the scRNA-seq reference data.

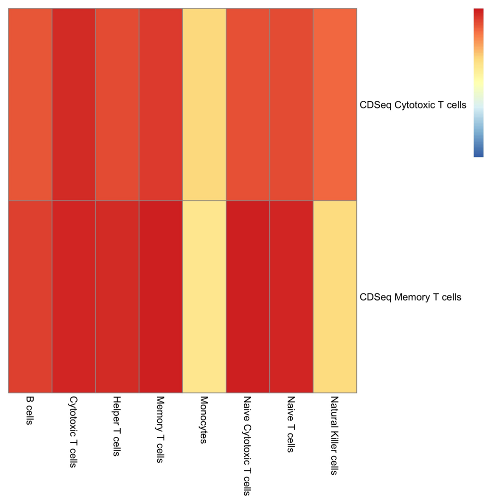


(a) Correlation between CDSeq-estimated cell-type-specific GEPs and true GEPs. Cell type number is 2.

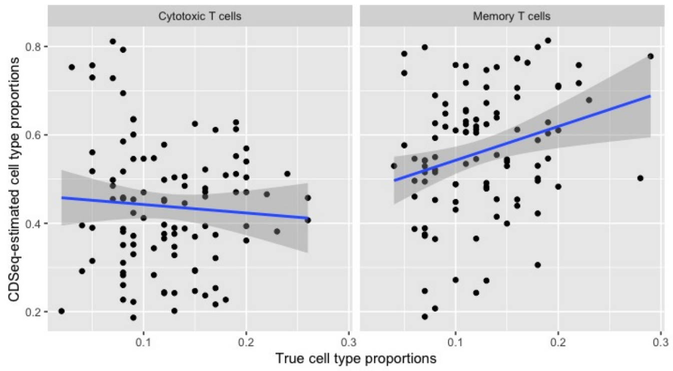


(b) Comparison between CDSeq-estimated cell type proportions and true proportion. Cell type number is 2

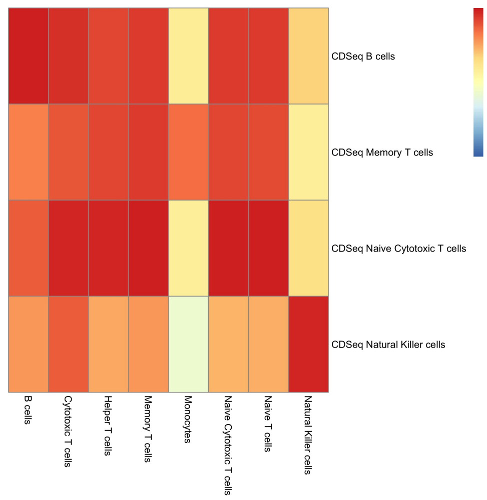


(c) Correlation between CDSeq-estimated cell-type-specific GEPs and true GEPs. Cell type number is 4.

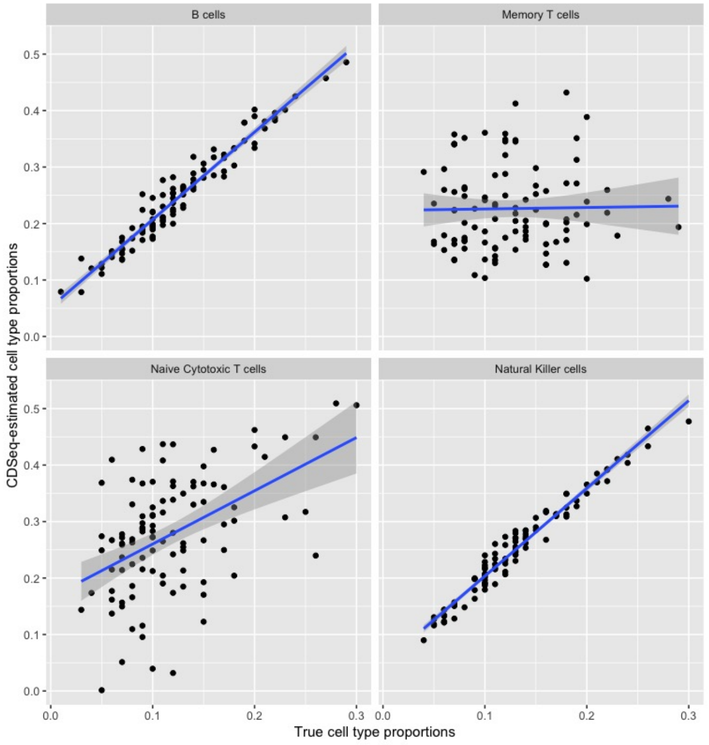


(d) Comparison between CDSeq-estimated cell type proportions and true proportion. Cell type number is 4.

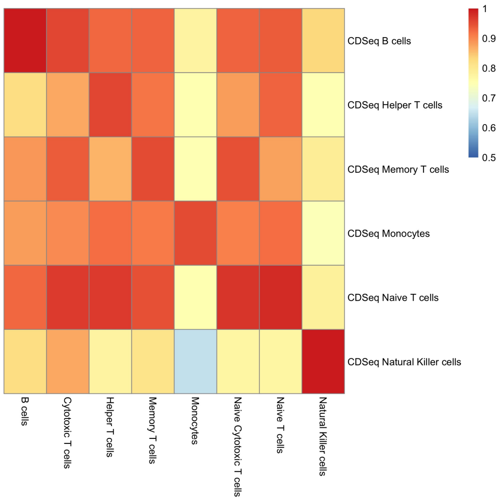


(e) Correlation between CDSeq-estimated cell-type-specific GEPs and true GEPs. Cell type number is 6.

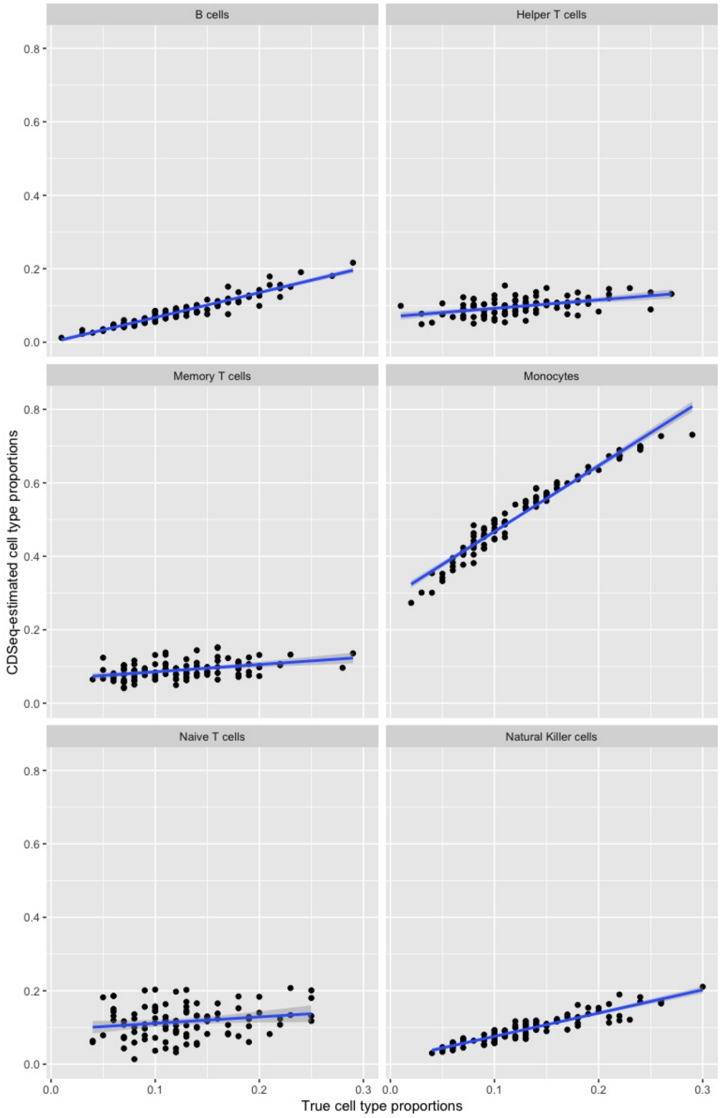


(f) Comparison between CDSeq-estimated cell type proportions and true proportion. Cell type number is 6.

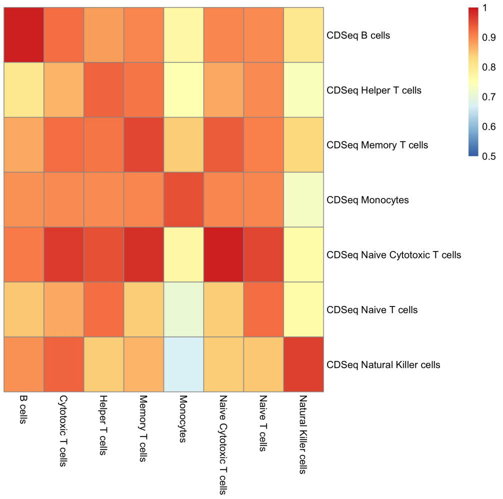


(g) Correlation between CDSeq-estimated cell-type-specific GEPs and true GEPs. Cell type number is 10.

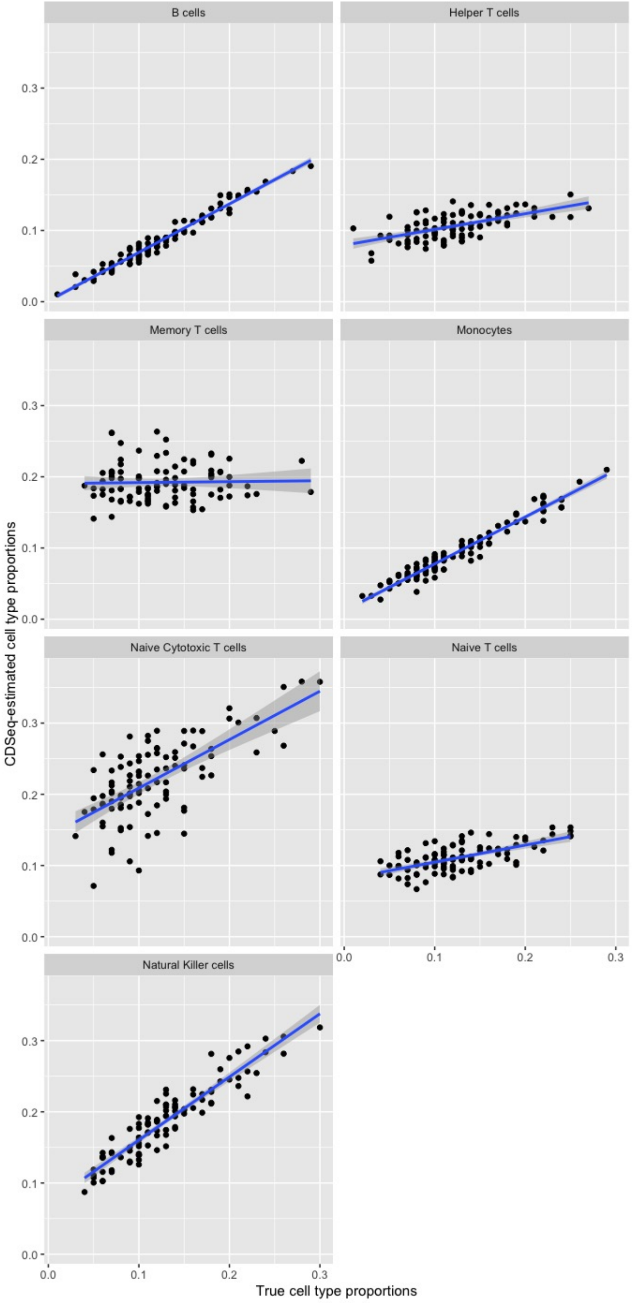


(h) Comparison between CDSeq-estimated cell type proportions and true proportion. Cell type number is 10.

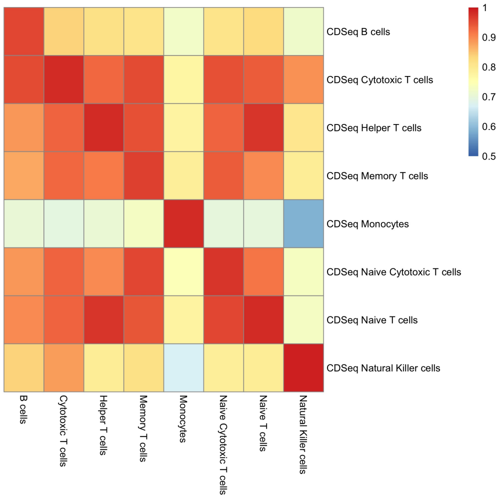


(i) Correlation between CDSeq-estimated cell-type-specific GEPs and true GEPs. Cell type number is 20.

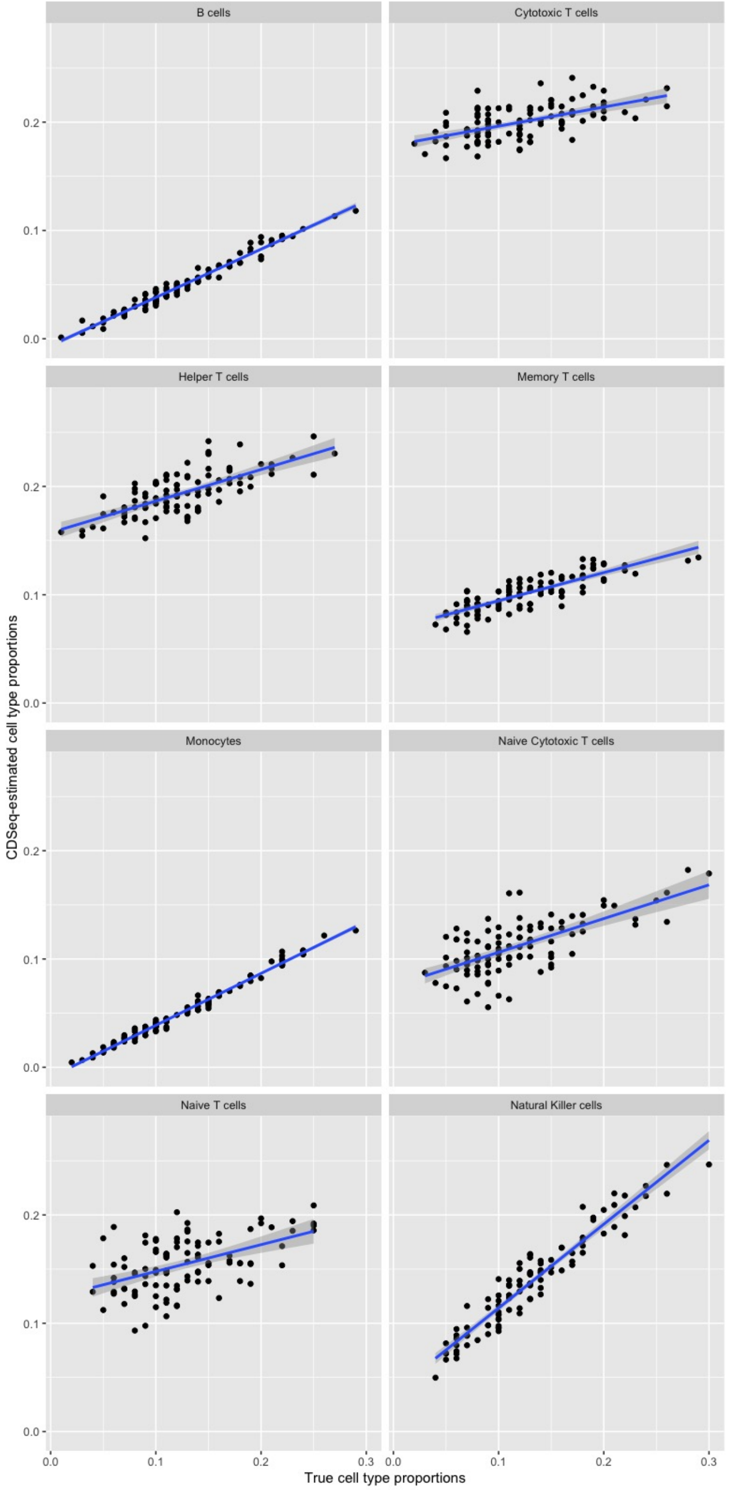


(j) Comparison between CDSeq-estimated cell type proportions and true proportion. Cell type number is 20.

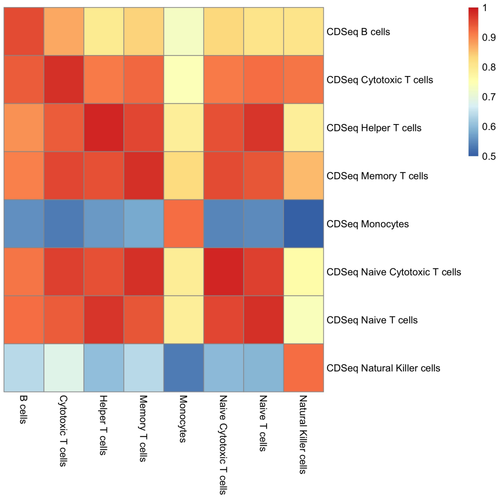


(k) Correlation between CDSeq-estimated cell-type-specific GEPs and true GEPs. Cell type number is 40.

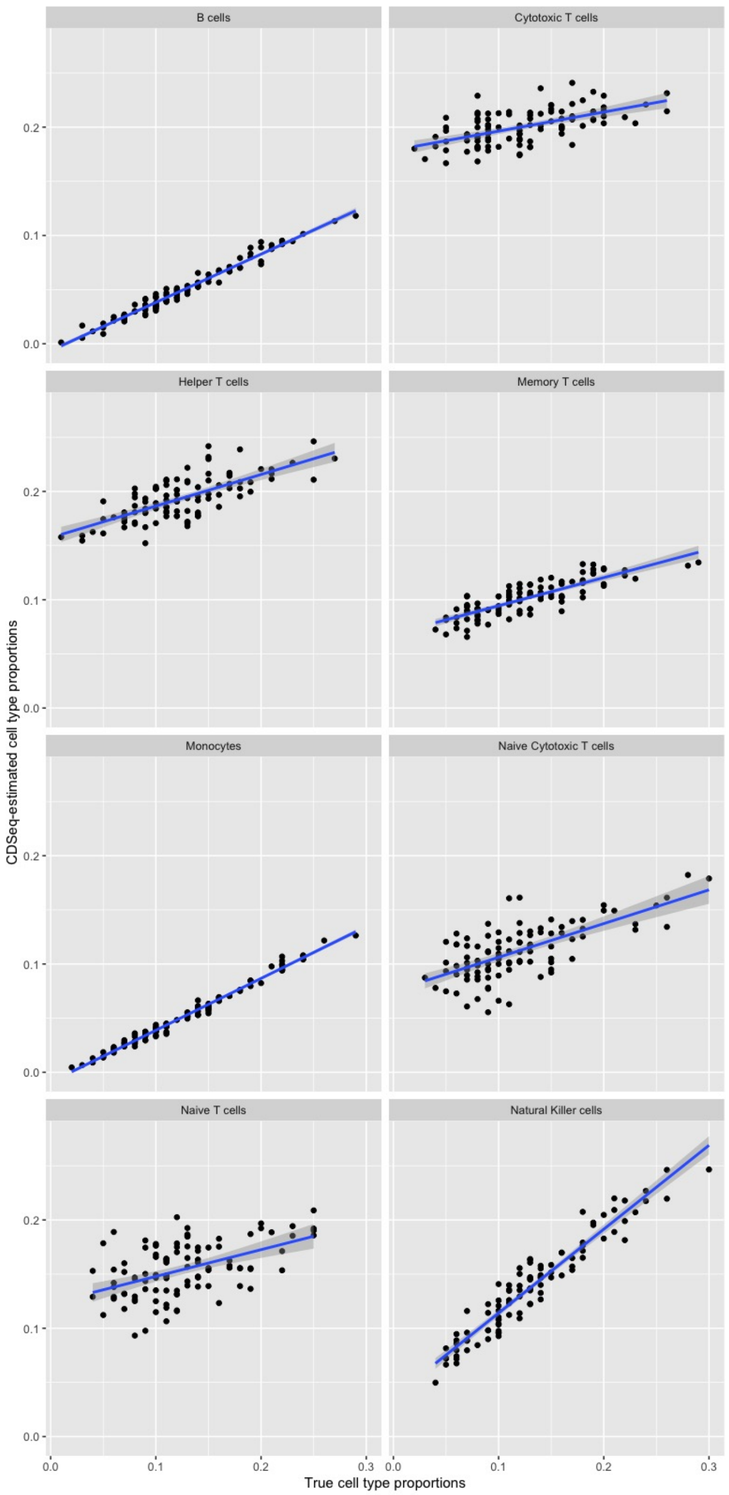


(l) Comparison between CDSeq-estimated cell type proportions and true proportion. Cell type number is 40.

Figure S21: Results of PBMC immune cell subtypes deconvolution. We created synthetic mixtures using 8 PBMC immune subtypes: B cells, Monocytes, Helper T cells, Natural Killer cells, Cytotoxic T cells, Memory T cells, Naive Cytotoxic T cells,Naive T cells [21]. We show results of using cell type number 2 (Fig.S21a and Fig.S21b), 4 (Fig.S21c and Fig.S21d), 6 (Fig.S21e and Fig.S21f), 10 (Fig.S21g and Fig.S21h), 20 (Fig.S21i and Fig.S21j), and 40 (Fig.S21k and Fig.S21l). For each case, we show the heatmap of correlation between CDSeq-estimated cell-type-specific GEPs and scatter plot comparing the CDSeq-estimated cell type proportions and true cell proportions. The blue lines are regression line. The results indicates CDSeq is able to detect the PBMC immune subtypes accurately. In this example, the number of cell types CDSeq estimated was 4, and the result suggests that for celll subtype detection, it may help to provide a larger number of cell types and then annotate the CDSeq-estimated cell types using a reference dataset. Note that CDSeq provides RNA proportion estimation whereas the true proportion is the cell proportion.

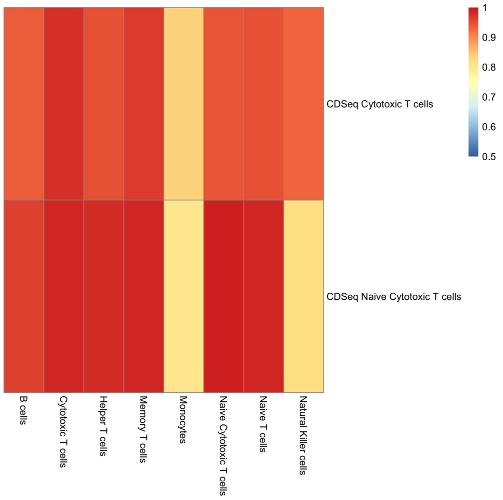


(a) Correlation between CDSeq-estimated cell-type-specific GEPs and true GEPs. Cell type number is 2.

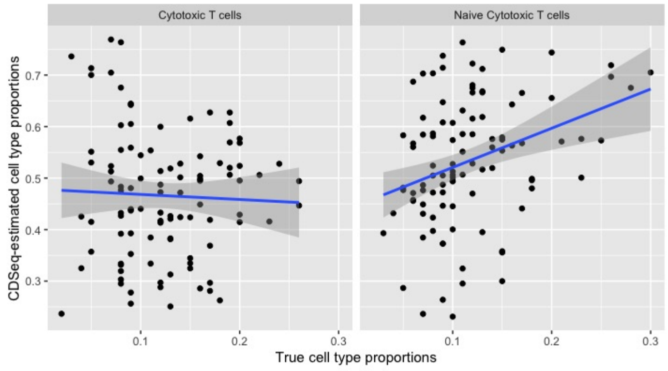


(b) Comparison between CDSeq-estimated cell type proportions and true proportion. Cell type number is 2

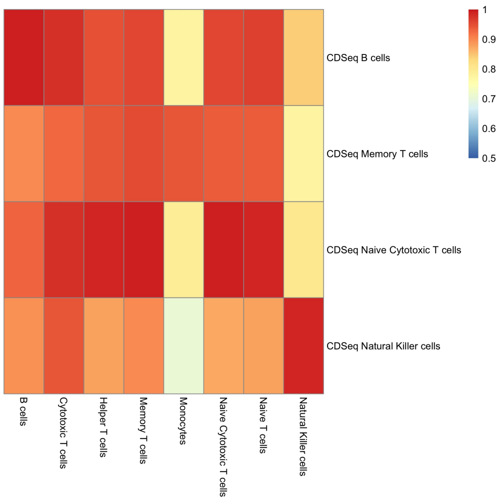


(c) Correlation between CDSeq-estimated cell-type-specific GEPs and true GEPs. Cell type number is 4.

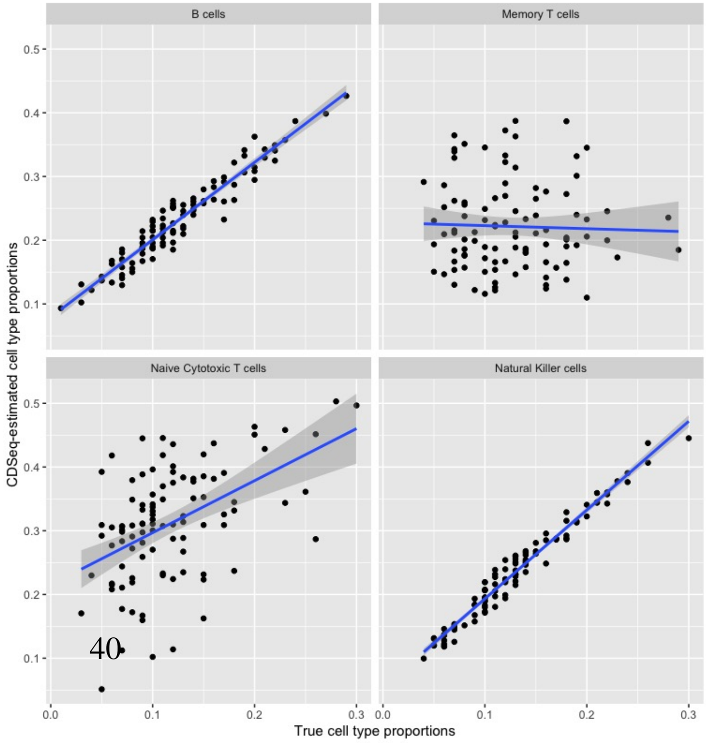


(d) Comparison between CDSeq-estimated cell type proportions and true proportion. Cell type number is 4.

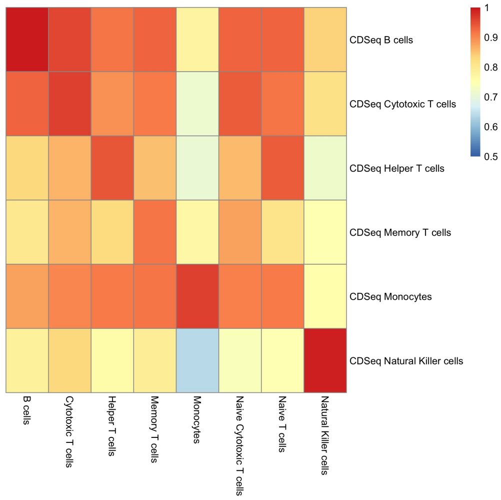


(e) Correlation between CDSeq-estimated cell-type-specific GEPs and true GEPs. Cell type number is 6.

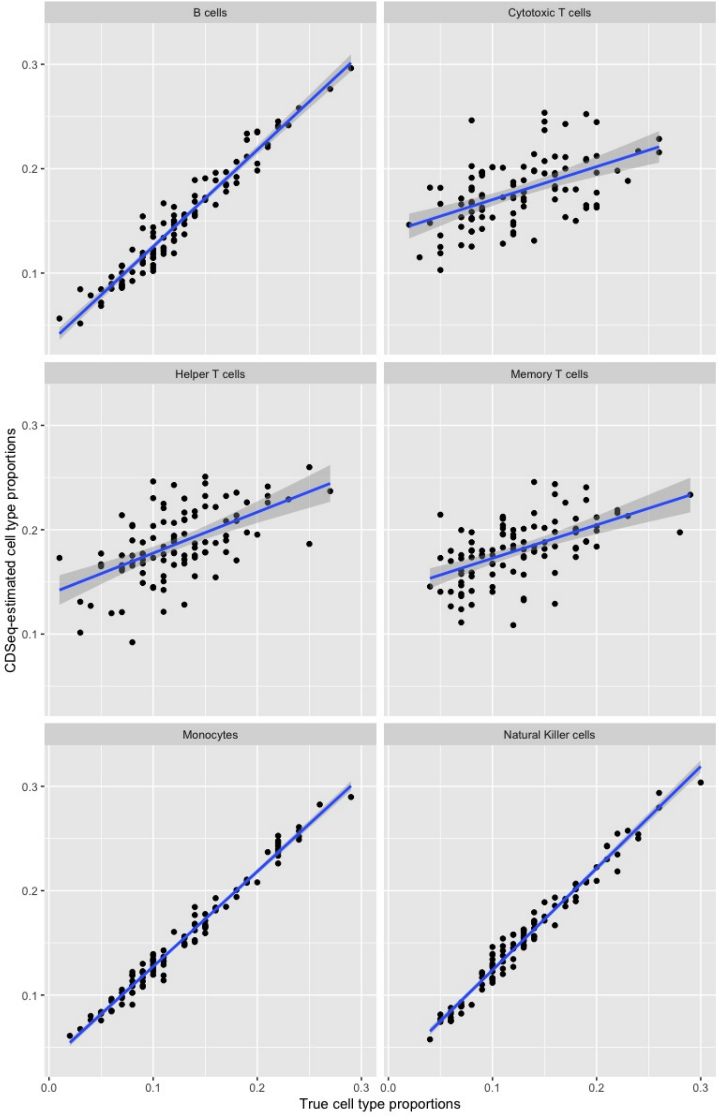


(f) Comparison between CDSeq-estimated cell type proportions and true proportion. Cell type number is 6.

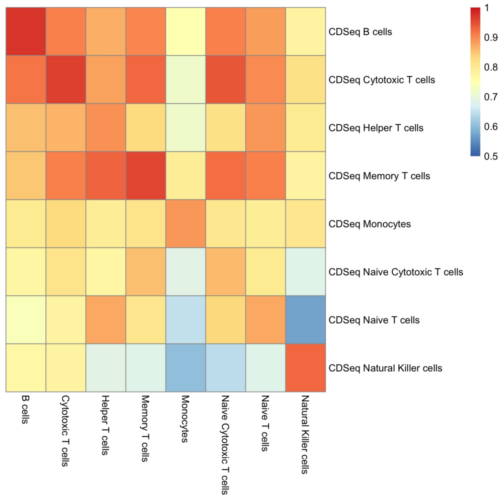


(g) Correlation between CDSeq-estimated cell-type-specific GEPs and true GEPs. Cell type number is 10.

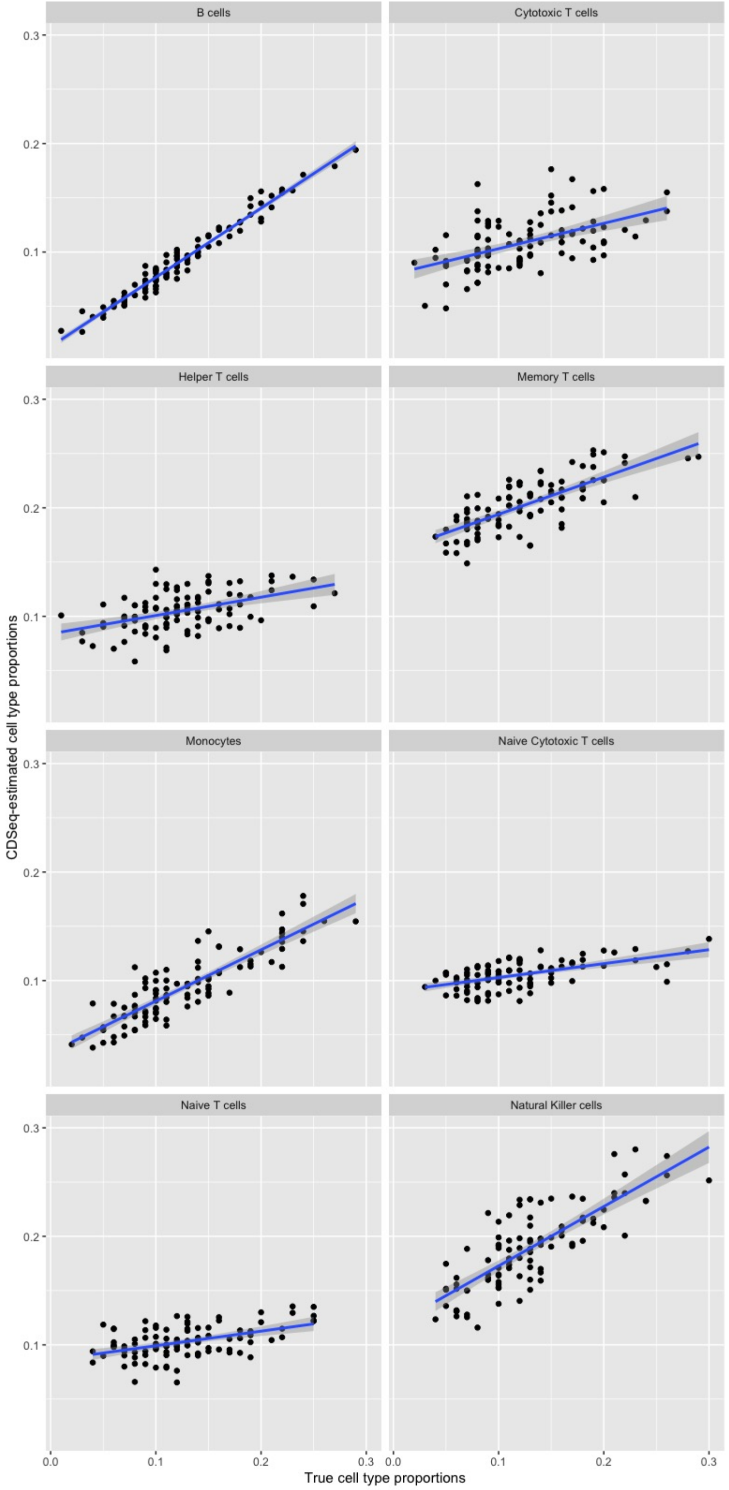


(h) Comparison between CDSeq-estimated cell type proportions and true proportion. Cell type number is 10.

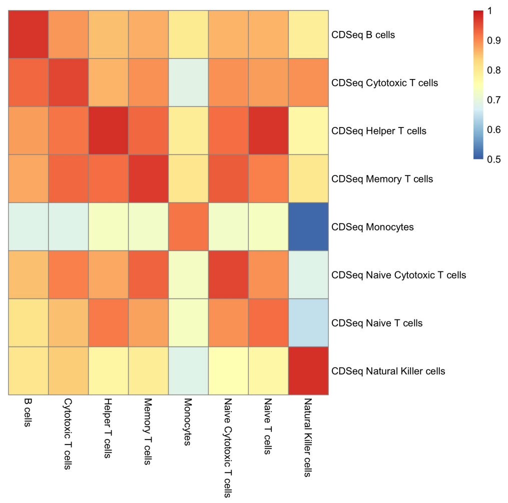


(i) Correlation between CDSeq-estimated cell-type-specific GEPs and true GEPs. Cell type number is 20.

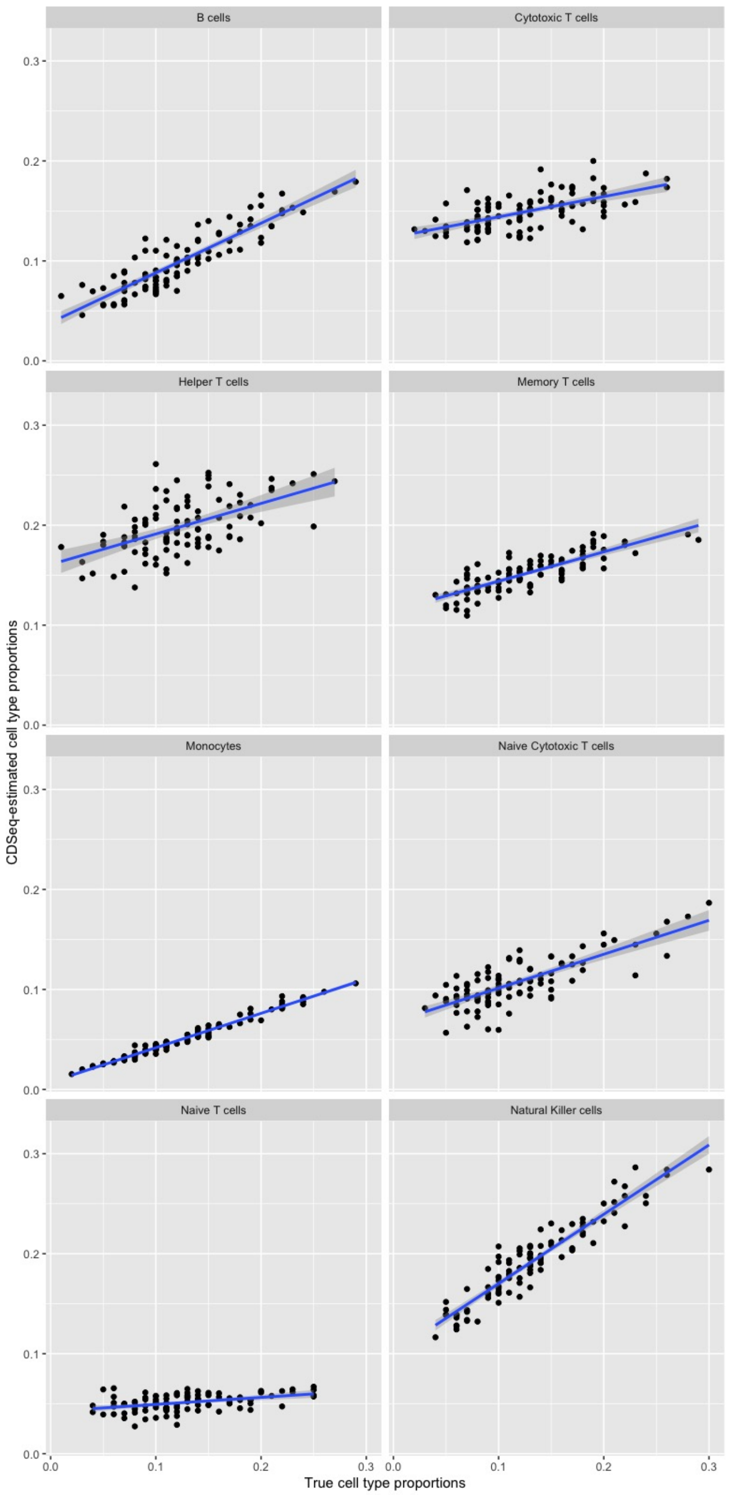


(j) Comparison between CDSeq-estimated cell type proportions and true proportion. Cell type number is 20.

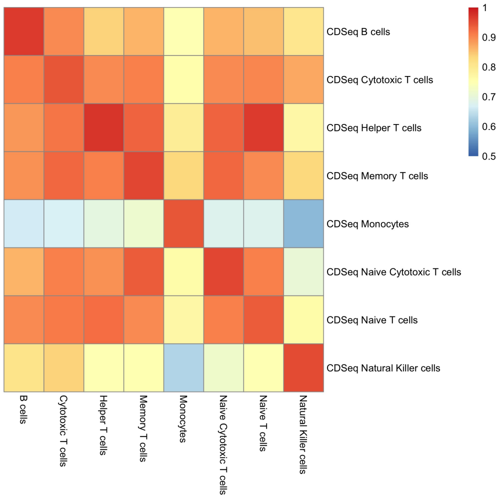


(k) Correlation between CDSeq-estimated cell-type-specific GEPs and true GEPs. Cell type number is 40.

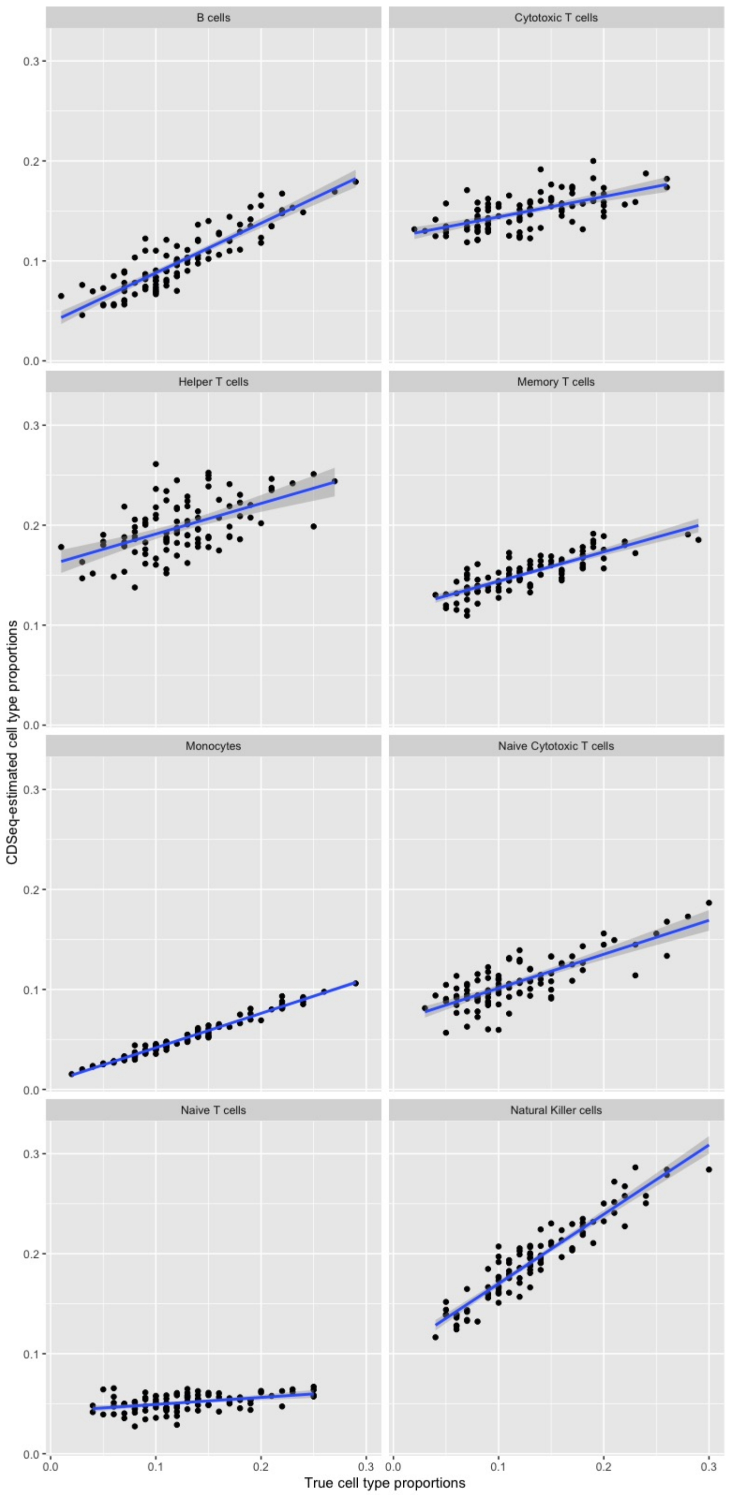


(l) Comparison between CDSeq-estimated cell type proportions and true proportion. Cell type number is 40.

Figure S22: Results of PBMC immune cell subtypes deconvolution using Reduce-Recover strategy. We created synthetic mixtures using 8 PBMC immune subtypes: B cells, Monocytes, Helper T cells, Natural Killer cells, Cytotoxic T cells, Memory T cells, Naive Cytotoxic T cells,Naive T cells [21]. We show results of using cell type number 2 (Fig.S22a and Fig.S22b), 4 (Fig.S22c and Fig.S22d), 6 (Fig.S22e and Fig.S22f), 10 (Fig.S22g and Fig.S22h), 20 (Fig.S22i and Fig.S22j), and 40 (Fig.S22k and Fig.S22l). For each case, we show the heatmap of correlation between CDSeq-estimated cell-type-specific GEPs and scatter plot comparing the CDSeq-estimated cell type proportions and true cell proportions. The blue lines are regression line. The results indicates CDSeq is able to detect the PBMC immune subtypes accurately. In this example, the number of cell types CDSeq estimated was 4, and the result suggests that for celll subtype detection, it may help to provide a larger number of cell types and then annotate the CDSeq-estimated cell types using a reference dataset. Note that CDSeq provides RNA proportion estimation whereas the true proportion is the cell proportion.

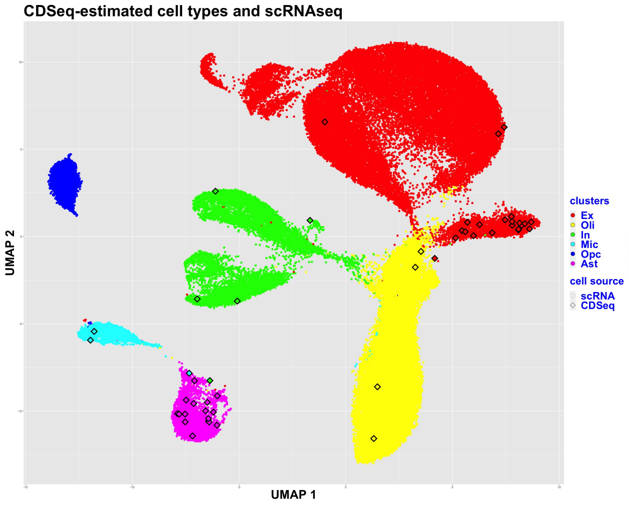


(a) UMAP embedding of CDSeq-estimated cell types and true cell types. Cell type number is 50.

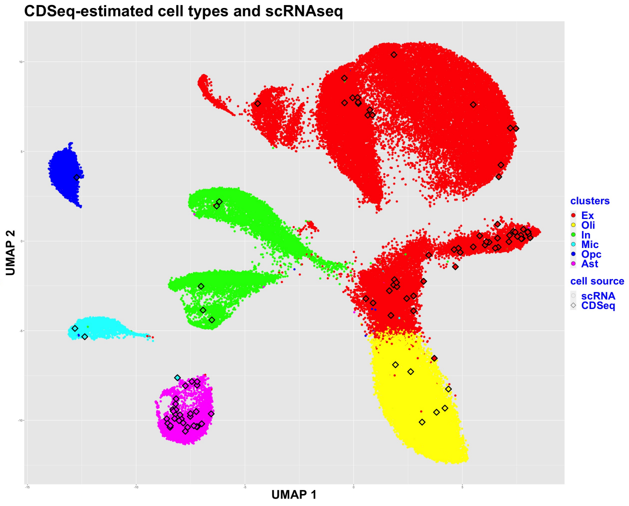


(b) UMAP embedding of CDSeq-estimated cell types and true cell types. Cell type number is 100.

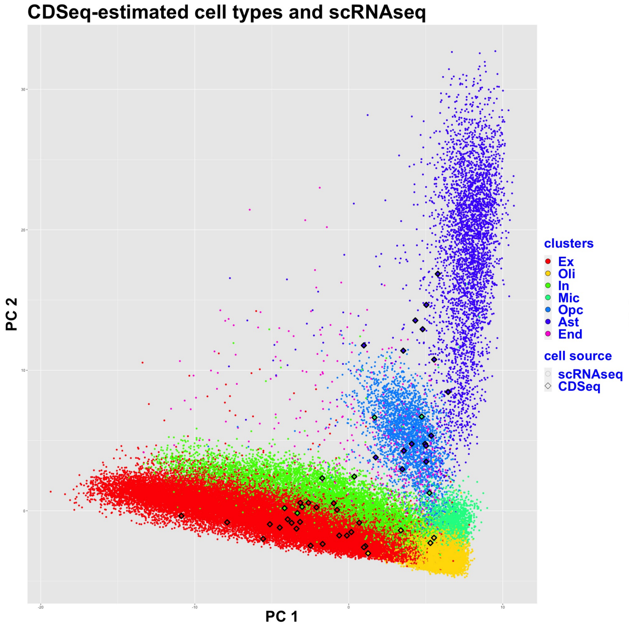


(c) PCA embedding of CDSeq-estimated cell types and true cell types. Cell type number is 50.

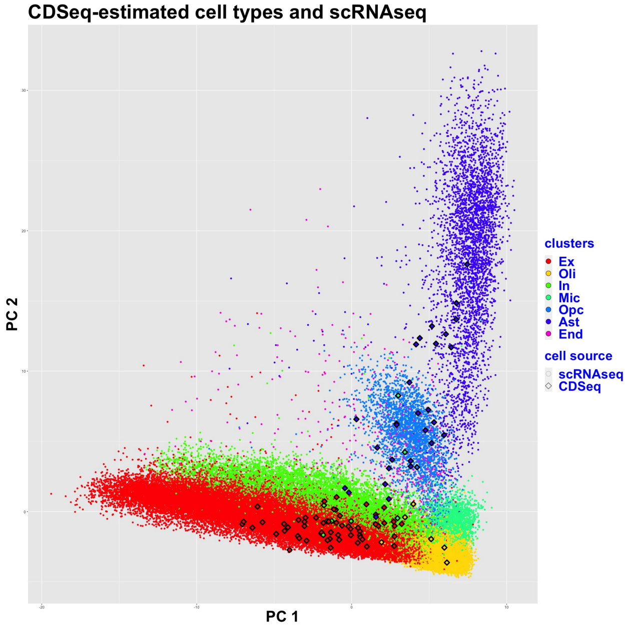


(d) PCA embedding of CDSeq-estimated cell types and true cell types. Cell type number is 100.

Figure S23: UMAP plot of the CDSeq-identified cell types in 209 GTEx frontal cortex RNA-seq bulk tissue samples and annotated individual cells from scRNA-seq data [15]. In the UMAP plot, the colored dots represent single cells of brain prefrontal cortex tissues from 48 individuals [15]. The colored squares denote the CDSeq-identified cell types. We show the results of deconvolving 209 GTEx frontal cortex samples using different number of cell types. The cell type annotation procedure suggests that when cell type number is 50, oligodendrocyte progenitor cell (opc) was not detected (Fig.S23a)probably due to its relatively low proportion, i.e. weaker signals compared to other cell types across the sample tissues. With cell type number 100, CDSeq is able to detect all 7 major cell types as shown in Fig.S23b. As shown, CDSeq-estimated cell types overlapped with all major clusters of single cells. CDSeqR identified the following cell types: excitatory neurons (Ex), inhibitory neurons (In), Oligodendrocyte (Oli), microglia (mic), oligodendrocyte progenitor cell (opc), astrocytes (ast), and endothelia (endo).

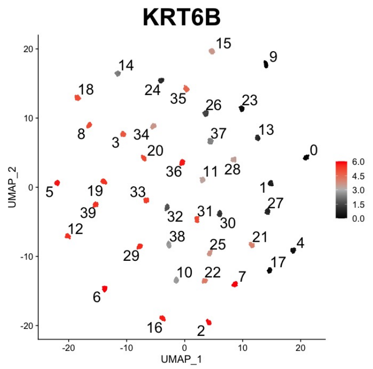


(a) Expression level of Epithelia cell marker KRT6B.

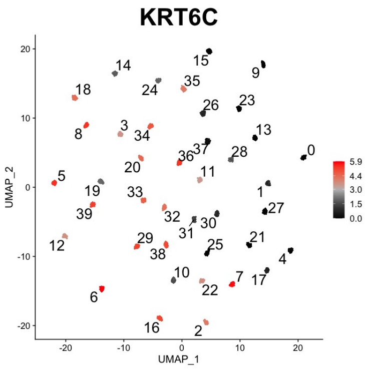


(b) Expression level of Epithelia cell marker KRT6C.

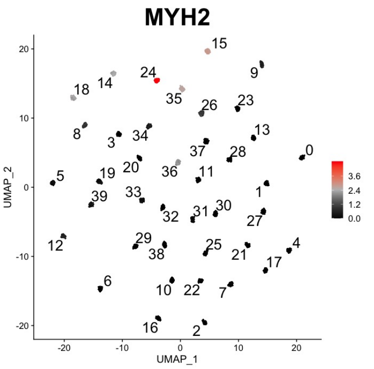


(c) Expression level of Myocytes cell marker MYH2.

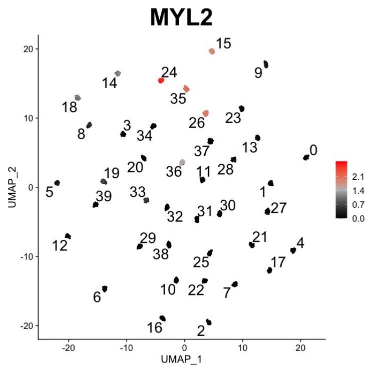


(d) Expression level of Myocytes cell marker MYL2.

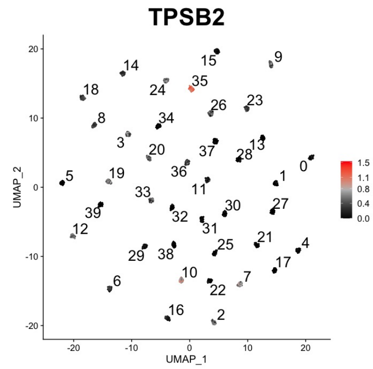


(e) Expression level of Mast cell marker TPSB2.

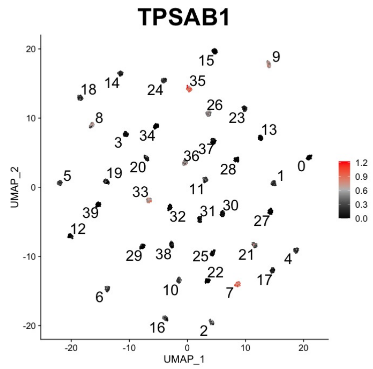


(f) Expression level of Mast cell marker TPSAB1.

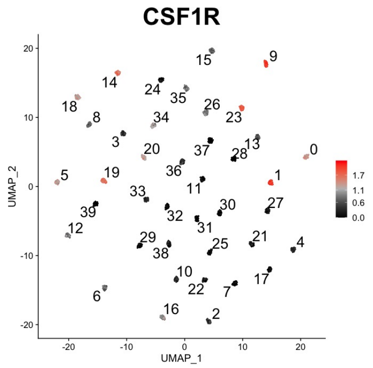


(g) Expression level of Macrophages cell marker CSF1R.

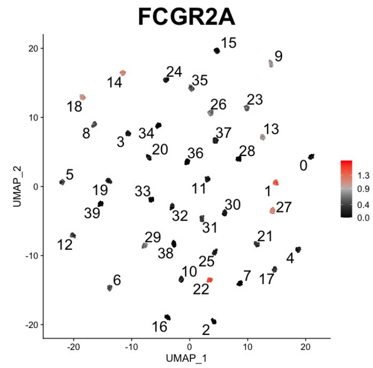


(h) Expression level of Macrophages cell marker FCGR2A.

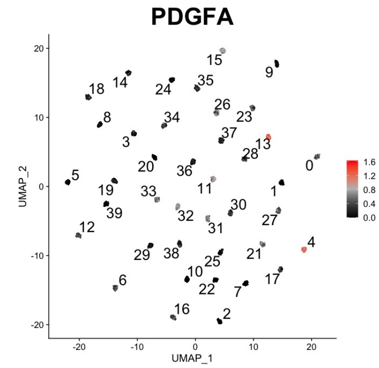


(i) Expression level of Myofibroblast cell marker PDGFA.

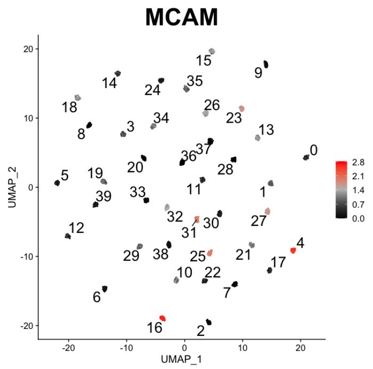


(j) Expression level of Myofibroblast cell marker MCAM.

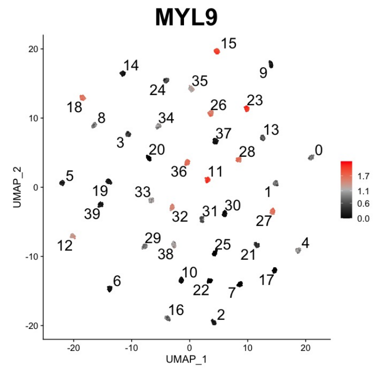


(k) Expression level of Myofibroblast cell marker CD3G.

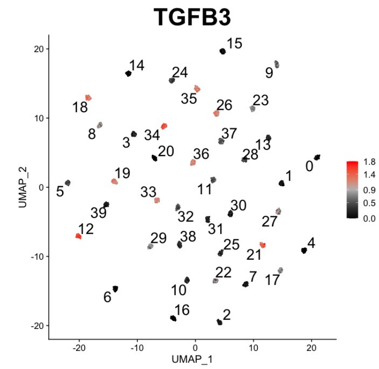


(l) Expression level of CAF cell marker TGFB3.

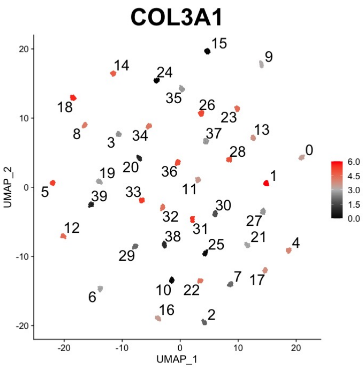


(m) Expression level of Fibroblast cell marker COL3A1.

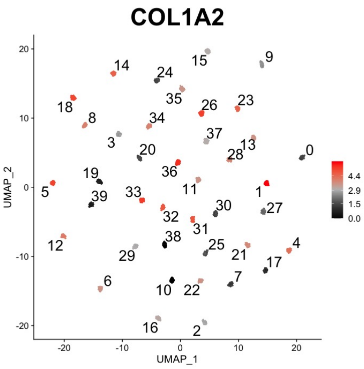


(n) Expression level of Fibroblast cell marker COL1A2.

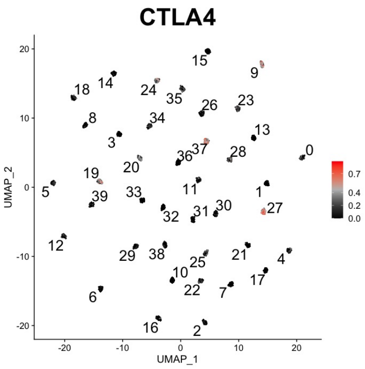


(o) Expression level of T cell marker CTLA4.

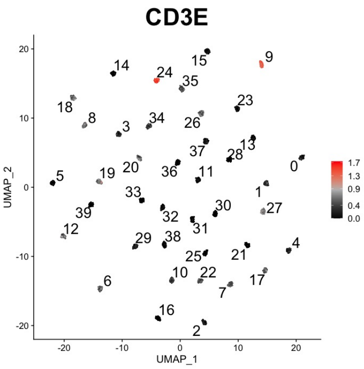


(p) Expression level of T cell marker CD3E.

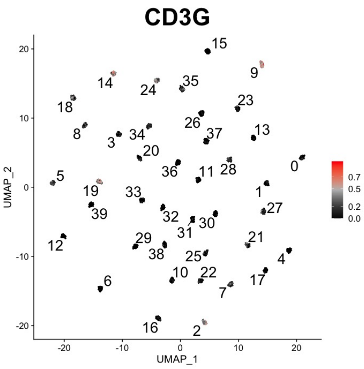


(q) Expression level of T cell marker CD3G.

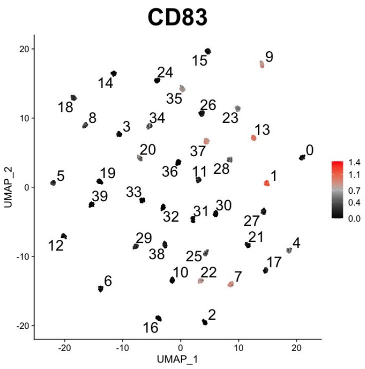


(r) Expression level of Dendritic cell marker CD83.

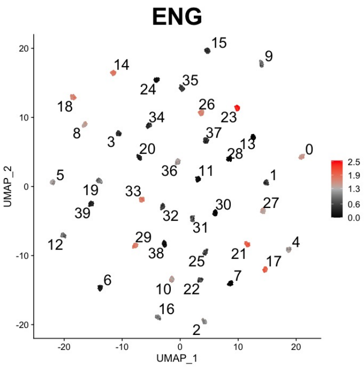


(s) Expression level of Endothelial cell marker ENG.

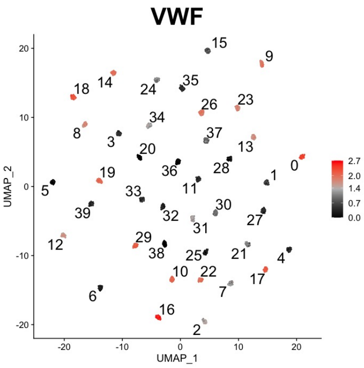


(t) Expression level of Endothelial cell marker VWF.

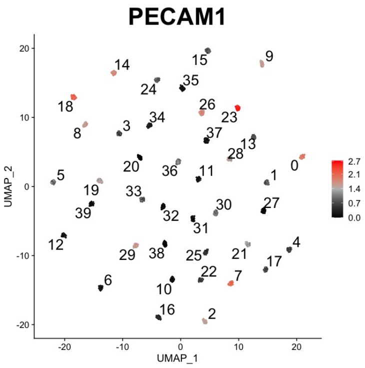


(u) Expression level of Endothelial cell marker PECAM1.

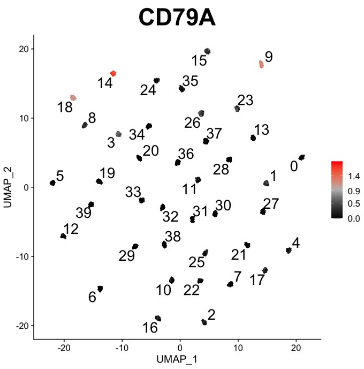


(v) Expression level of B cell marker CD79A.

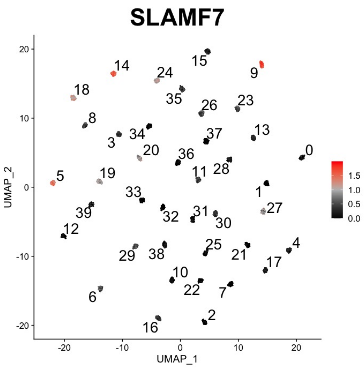


(w) Expression level of B cell marker SLAMF7.

Figure S24: Deconvolution of TCGA-HNSC (Head-Neck Squamous Cell Carcinoma) RNA-seq data. We randomly selected 20 tumor samples. Deconvolution was carried out using top 1000 variable genes in the bulk data. We show the results when cell type number is 40 considering the high level of heterogeneity suggested by single cell RNA-seq analysis of head and neck squamous cell carcinoma [22]. To assign cell types, we examined the expression levels of multiple marker genes as shown in the figures. We identified a diverse group of epithelial cells in CDSeq-estimated cell types: 2, 3, 5, 6, 7, 8, 12, 16, 18, 19, 20, 21, 22, 25, 29, 30, 31, 33, 34, 36, 38, 39. This finding is consistent with the findings in [22]. The study showed that malignant epithelial cells from each individual form their own clusters unlike normal cell type that form clusters based on their cell type identities. Based on the expression level of marker genes, we could potentially identify that cell type 0, 10 and 17 are endothelial cells; cell type 1 is macrophages; cell type 4, 11, 13, 15, 26, 28, 32 are Fibroblast cells (Myofibroblast or cancer-associated fibroblast (CAF) cells); cell type 9 and 27 are T cell; cell type 14 is B cell; cell type 24 is myocyte; cell type 35 is mast cell; cell type 37 is Dendritic cell.

### References


[1] Kang, K., Meng, Q., Shats, I., Umbach, D. M., Li, M., Li, Y., Li, X., and Li, L. (2019) CDSeq: A novel complete deconvolution method for dissecting heterogeneous samples using gene expression data. *PLoS Computational Biology,* **15**(12).

[2] Cobos, F. A., Alquicira-Hernandez, J., Powell, J. E., Mestdagh, P., and De Preter, K. (2020) Benchmarking of cell type deconvolution pipelines for transcriptomics data. *Nature communications,* **11**(1), 1–14.

[3] Newman, A. M., Liu, C. L., Green, M. R., Gentles, A. J., Feng, W., Xu, Y., Hoang, C. D., Diehn, M., and Alizadeh, A. A. (2015) Robust enumeration of cell subsets from tissue expression profiles. *Nature methods,* **12**(5), 453–457.

[4] Wang, X., Park, J., Susztak, K., Zhang, N. R., and Li, M. (2019) Bulk tissue cell type deconvolution with multi-subject single-cell expression reference. *Nature communications,* **10**(1),

1–9.

[5] Du, R., Carey, V., and Weiss, S. T. (2019) deconvSeq: deconvolution of cell mixture distribution in sequencing data. *Bioinformatics,* **35**(24), 5095–5102.

[6] Mullen, K. M. and Mullen, M. K. M. (2015) The nnls package.

[7] Dong, M., Thennavan, A., Urrutia, E., Li, Y., Perou, C. M., Zou, F., and Jiang, Y. (2020) SCDC: bulk gene expression deconvolution by multiple single-cell RNA sequencing references. *Briefings in bioinformatics,*.

[8] Altboum, Z., Steuerman, Y., David, E., Barnett-Itzhaki, Z., Valadarsky, L., Keren-Shaul, H., Meningher, T., Mendelson, E., Mandelboim, M., Gat-Viks, I., et al. (2014) Digital cell quantification identifies global immune cell dynamics during influenza infection. *Molecular systems biology,* **10**(2), 720.

[9] Friedman, J., Hastie, T., and Tibshirani, R. (2010) Regularization paths for generalized linear models via coordinate descent. *Journal of statistical software,* **33**(1), 1.

[10] Zaitsev, K., Bambouskova, M., Swain, A., and Artyomov, M. N. (2019) Complete deconvolution of cellular mixtures based on linearity of transcriptional signatures. *Nature communications,* **10**(1), 1–16.

[11] Jew, B., Alvarez, M., Rahmani, E., Miao, Z., Ko, A., Garske, K. M., Sul, J. H., Pietiläinen, K. H., Pajukanta, P., and Halperin, E. (2020) Accurate estimation of cell composition in bulk expression through robust integration of single-cell information. *Nature communications,* **11**(1), 1–11.

[12] Hao, Y., Yan, M., Heath, B. R., Lei, Y. L., and Xie, Y. (2019) Fast and robust deconvolution of tumor infiltrating lymphocyte from expression profiles using least trimmed squares. *PLoS computational biology,* **15**(5), e1006976.

[13] Ripley, B. et al. (2011) MASS: support functions and datasets for Venables and Ripley's MASS. *R package version,* **7**, 3–29.

[14] Chambers, J., Hastie, T., and Pregibon, D. (1990) Statistical models in S. In *Compstat* Springer pp. 317–321.

[15] Mathys, H., Davila-Velderrain, J., Peng, Z., Gao, F., Mohammadi, S., Young, J. Z., Menon, M., He, L., Abdurrob, F., Jiang, X., et al. (2019) Single-cell transcriptomic analysis of Alzheimer's disease. *Nature,* **570**(7761), 332–337.

[16] Ding, J., Adiconis, X., Simmons, S. K., Kowalczyk, M. S., Hession, C. C., Marjanovic, N. D., Hughes, T. K., Wadsworth, M. H., Burks, T., Nguyen, L. T., et al. (2020) Systematic comparison of single-cell and single-nucleus RNA-sequencing methods. *Nature biotechnology,* **38**(6), 737–746.

[17] Puram, S. V., Tirosh, I., Parikh, A. S., Patel, A. P., Yizhak, K., Gillespie, S., Rodman, C., Luo, C. L., Mroz, E. A., Emerick, K. S., et al. (2017) Single-cell transcriptomic analysis of primary and metastatic tumor ecosystems in head and neck cancer. *Cell,* **171**(7), 1611–1624.

[18] Zaitsev, K., Bambouskova, M., Swain, A., and Artyomov, M. N. (2019) Complete deconvolution of cellular mixtures based on linearity of transcriptional signatures. *Nature communications,* **10**(1), 1–16.

[19] Walker, D. G. (2020) Defining activation states of microglia in human brain tissue: an unresolved issue for Alzheimer's disease. *Neuroimmunology and Neuroinflammation,* **7**(3), 194–214.

[20] McKenzie, A. T., Wang, M., Hauberg, M. E., Fullard, J. F., Kozlenkov, A., Keenan, A., Hurd, Y. L., Dracheva, S., Casaccia, P., Roussos, P., et al. (2018) Brain cell type specific gene expression and co-expression network architectures. *Scientific reports,* **8**(1), 1–19.

[21] Zheng, G. X., Terry, J. M., Belgrader, P., Ryvkin, P., Bent, Z. W., Wilson, R., Ziraldo, S. B., Wheeler, T. D., McDermott, G. P., Zhu, J., et al. (2017) Massively parallel digital transcriptional profiling of single cells. *Nature communications,* **8**(1), 1–12.

[22] Puram, S. V., Tirosh, I., Parikh, A. S., Patel, A. P., Yizhak, K., Gillespie, S., Rodman, C., Luo, C. L., Mroz, E. A., Emerick, K. S., et al. (2017) Single-cell transcriptomic analysis of primary and metastatic tumor ecosystems in head and neck cancer. *Cell,* **171**(7), 1611–1624.
